# A cross-package Bioconductor workflow for analysing methylation array data

**DOI:** 10.1101/055087

**Authors:** Jovana Maksimovic, Belinda Phipson, Alicia Oshlack

## Abstract

Methylation in the human genome is known to be associated with development and disease. The Illumina Infinium methylation arrays are by far the most common way to interrogate methylation across the human genome. This paper provides a Bioconductor workflow using multiple packages for the analysis of methylation array data. Specifically, we demonstrate the steps involved in a typical differential methylation analysis pipeline including: quality control, filtering, normalization, data exploration and statistical testing for probe-wise differential methylation. We further outline other analyses such as differential methylation of regions, differential variability analysis, estimating cell type composition and gene ontology testing. Finally, we provide some examples of how to visualise methylation array data.

## Introduction

DNA methylation, the addition of a methyl group to a CG dinucleotide of the DNA, is the most extensively studied epigenetic mark due to its role in both development and disease (Bird 2002; Laird 2003). Although DNA methylation can be measured in several ways, the epigenetics community has enthusiastically embraced the Illumina HumanMethylation450 (450k) array (Bibikova et al. 2011) as a cost-effective way to assay methylation across the human genome. More recently, Illumina has increased the genomic coverage of the platform to >850,000 sites with the release of their MethylationEPIC (850k) array. As methylation arrays are likely to remain popular for measuring methylation for the foreseeable future, it is necessary to provide robust workflows for methylation array analysis.

Measurement of DNA methylation by Infinium technology (Infinium I) was first employed by Illumina on the HumanMethylation27 (27k) array (Bibikova et al. 2009), which measured methylation at approximately 27,000 CpGs, primarily in gene promoters. Like bisulfite sequencing, the Infinium assay detects methylation status at single base resolution. However, due to its relatively limited coverage the array platform was not truly considered “genome-wide” until the arrival of the 450k array. The 450k array increased the genomic coverage of the platform to over 450,000 gene-centric sites by combining the original Infinium I assay with the novel Infinium II probes. Both assay types employ 50bp probes that query a [C/T] polymorphism created by bisulfite conversion of unmethylated cytosines in the genome, however, the Infinium I and II assays differ in the number of beads required to detect methylation at a single locus. Infinium I uses two bead types per CpG, one for each of the methylated and unmethylated states (Figure ??a). In contrast, the Infinium II design uses one bead type and the methylated state is determined at the single base extension step after hybridization (Figure ??b). The 850k array also uses a combination of the Infinium I and II assays but achieves additional coverage by increasing the size of each array; a 450k slide contains 12 arrays whilst the 850k has only 8.

Regardless of the Illumina array version, for each CpG, there are two measurements: a methylated intensity (denoted by *M*) and an unmethylated intensity (denoted by *U*). These intensity values can be used to determine the proportion of methylation at each CpG locus. Methylation levels are commonly reported as either beta values (*β* = *M*/(*M* + *U* + *α*)) or M-values (*M value* = *log2*(*M*/*U*)). Beta values and M-values are related through a logit transformation. Beta values are generally preferable for describing the level of methylation at a locus or for graphical presentation because percentage methylation is easily interpretable. However, due to their distributional properties, M-values are more appropriate for statistical testing (Du et al. 2010).

**Figure 1:**
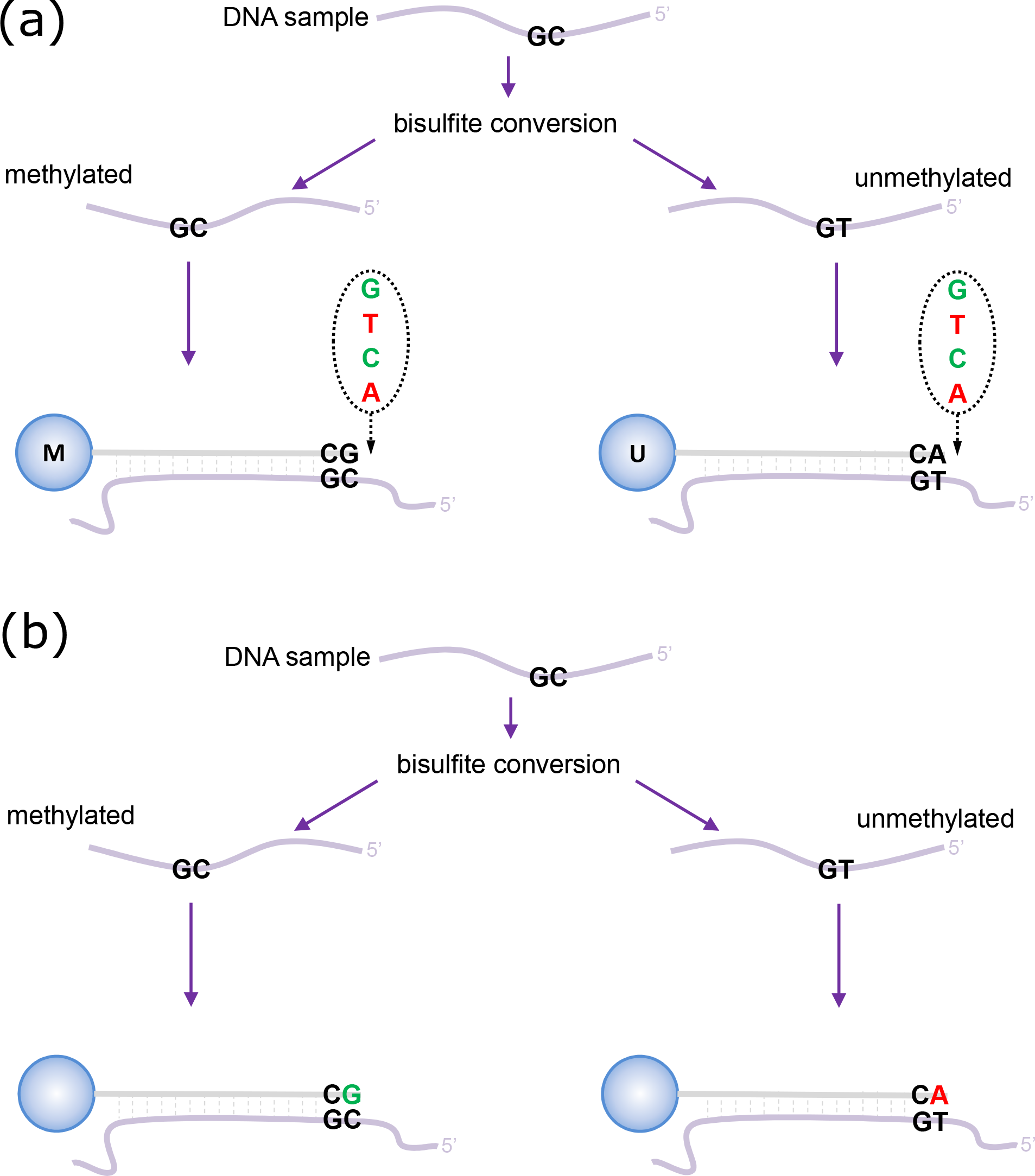
Illumina Infinium HumanMethylation450 assay, reproduced from Maksimovic, Gordon and Oshlack 2012. (a) Infinium I assay. Each individual CpG is interrogated using two bead types: methylated (M) and unmethylated (U). Both bead types will incorporate the same labeled nucleotide for the same target CpG, thereby producing the same color fluorescence. The nucleotide that is added is determined by the base downstream of the ‘C’ of the target CpG. The proportion of methylation can be calculated by comparing the intensities from the two different probes in the same color. (b) Infinium II assay. Each target CpG is interrogated using a single bead type. Methylation state is detected by single base extension at the position of the ‘C’ of the target CpG, which always results in the addition of a labeled ‘G’ or ‘A’ nucleotide, complementary to either the ‘methylated’ C or ‘unmethylated’ T, respectively. Each locus is detected in two colors, and methylation status is determined by comparing the two colors from the one position.

In this workflow, we will provide examples of the steps involved in analysing methylation array data using R (R Core Team 2014) and Bioconductor (Huber et al. 2015), including: quality control, filtering, normalization, data exploration and probe-wise differential methylation analysis. We will also cover other approaches such as differential methylation analysis of regions, differential variability analysis, gene ontology analysis and estimating cell type composition. Finally, we will provide some examples of useful ways to visualise methylation array data.

## Differential Methylation Analysis

To demonstrate the various aspects of analysing methylation data, we will be using a small, publicly available 450k methylation dataset (Y. Zhang et al. 2013). The dataset contains 10 samples in total; there are 4 different sorted T-cell types (naive, rTreg, act_naive, act_rTreg), collected from 3 different individuals (M28, M29, M30). For details describing sample collection and preparation, see Y. Zhang et al. (2013). An additional birth sample (individual VICS-72098-18-B) is included from another study (Cruickshank et al. 2013) to illustrate approaches for identifying and excluding poor quality samples.

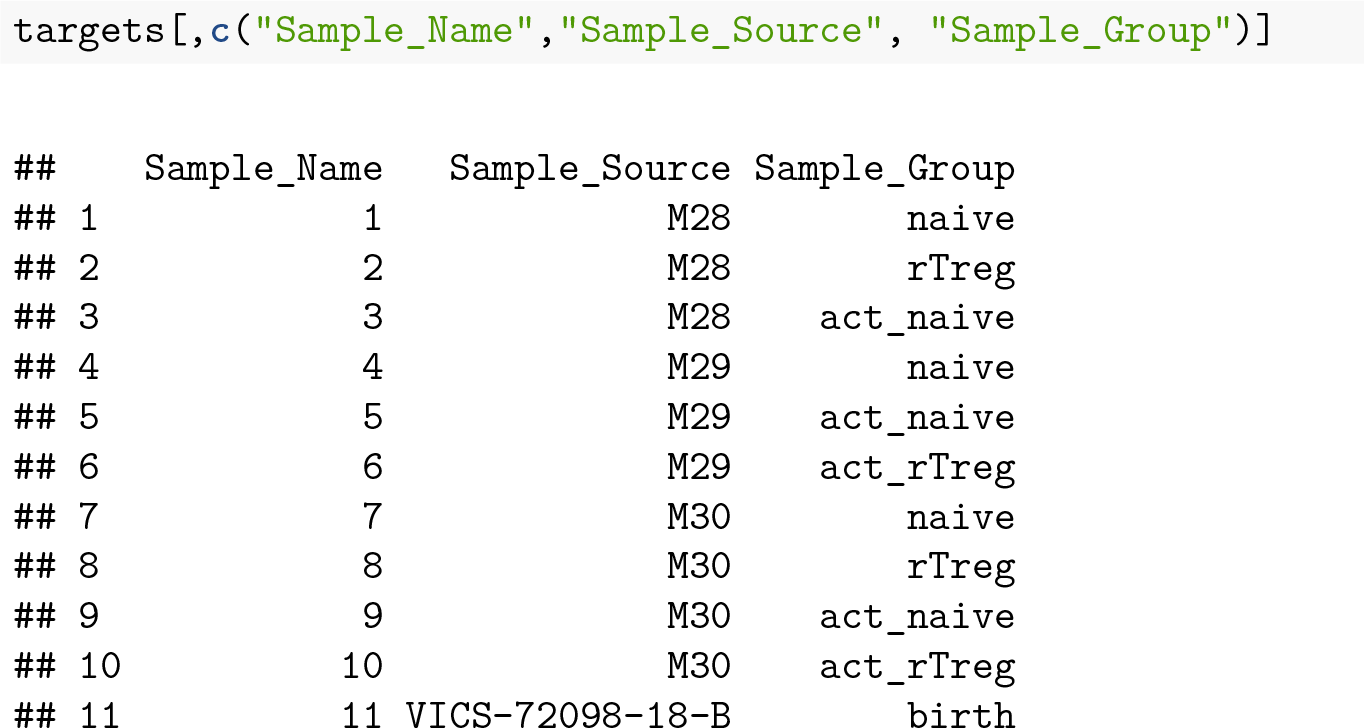

There are several R Bioconductor packages available that have been developed for analysing methylation array data, including *minfi* (Aryee et al. 2014), *missMethyl* (B. Phipson, Maksimovic, and Oshlack 2016), *wateRmelon* (Pidsley et al. 2013), *methylumi* (S. Davis et al. 2015), *ChAMP* (Morris et al. 2014) and *charm* (Aryee et al. 2011). Some of the packages, such as *minfi* and *methylumi* include a framework for reading in the raw data from IDAT files and various specialised objects for storing and manipulating the data throughout the course of an analysis. Other packages provide specialised analysis methods for normalisation and statistical testing that rely on either *minfi* or *methylumi* objects. It is possible to convert between *minfi* and *methylumi* data types, however, this is not always trivial. Thus, it is advisable to consider the methods that you are interested in using and the data types that are most appropriate before you begin your analysis. Another popular method for analysing methylation array data is *limma* (Ritchie et al. 2015), which was originally developed for gene expression microarray analysis. As *limma* operates on a matrix of values, it is easily applied to any data that can be converted to a **`matrix`** in R.

We will begin with an example of a **probe-wise** differential methylation analysis using *minfi* and *limma*. By **probe-wise** analysis we mean each individual CpG probe will be tested for differential methylation for the comparisons of interest and p-values and moderated t-statistics will be generated for each CpG probe.

## Loading the Data

It is useful to begin an analysis in R by loading all the package libraries that are likely to be required.

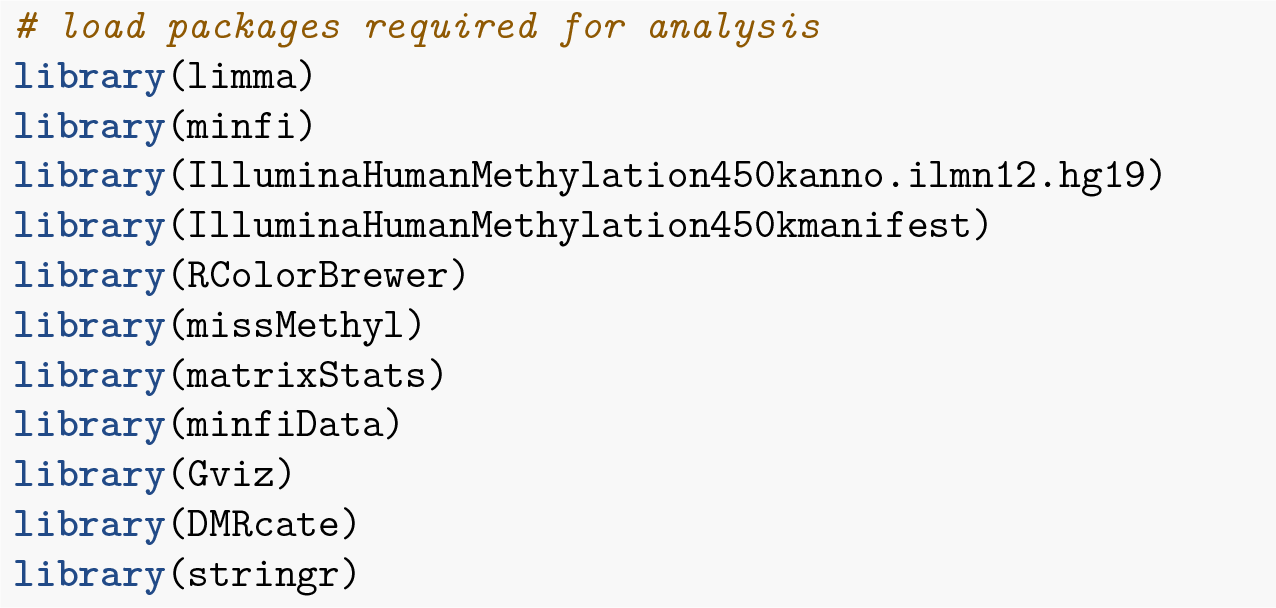

The *minfi* package provides the Illumina manifest as an R object which can easily be loaded into the environment. The manifest contains all of the annotation information for each of the CpG probes on the 450k array. This is useful for determining where any differentially methylated probes are located in a genomic context.

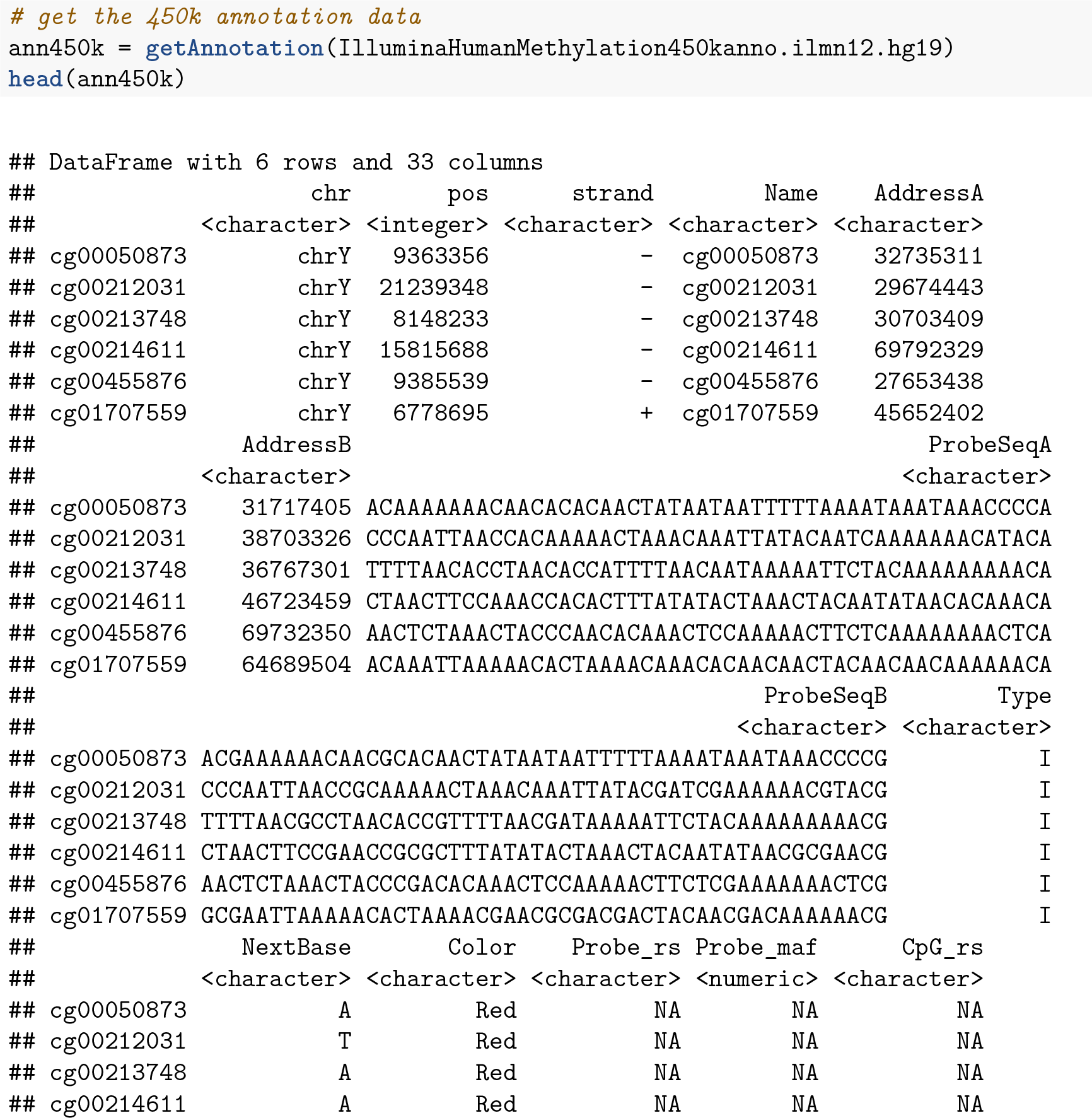

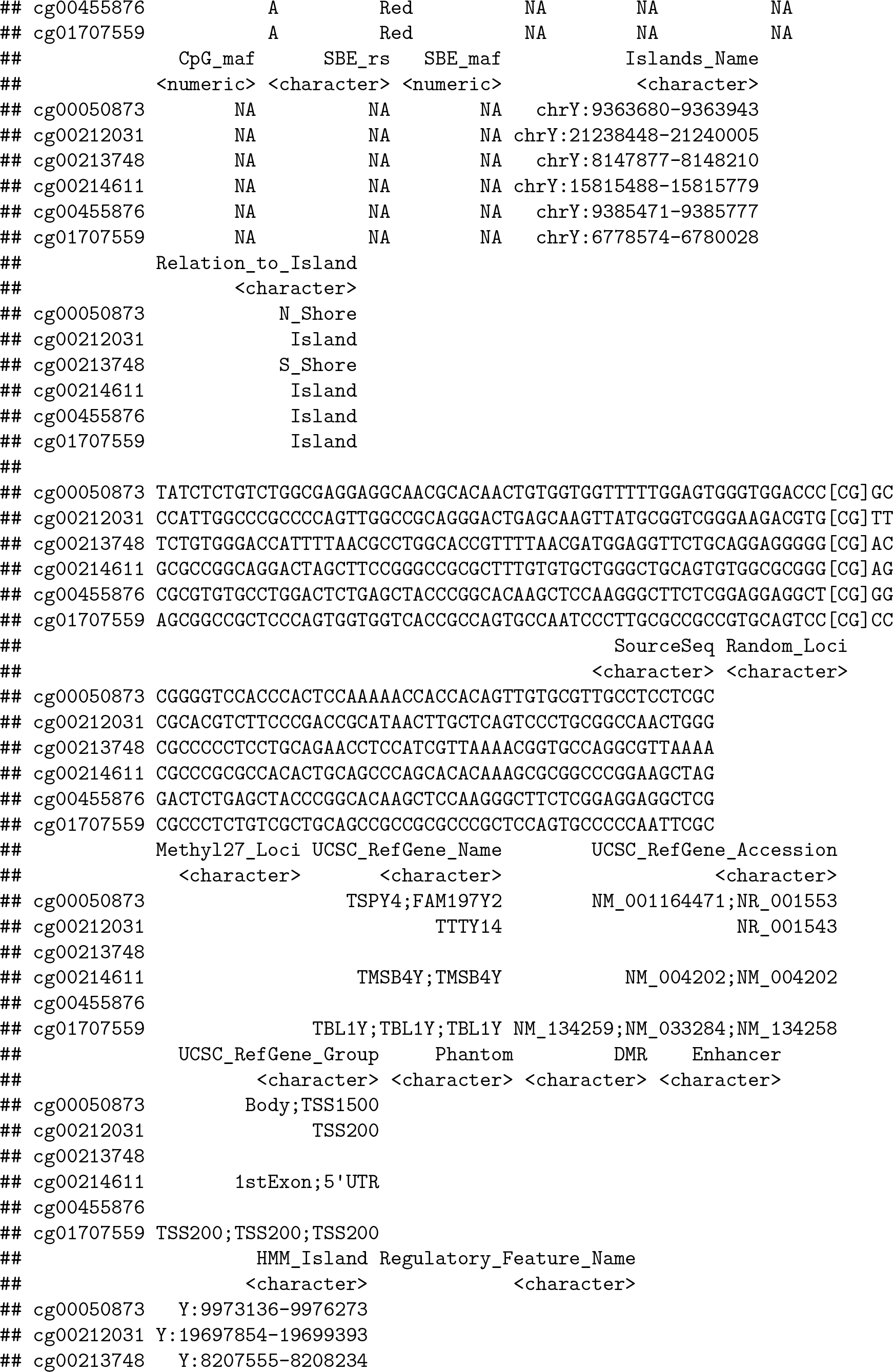

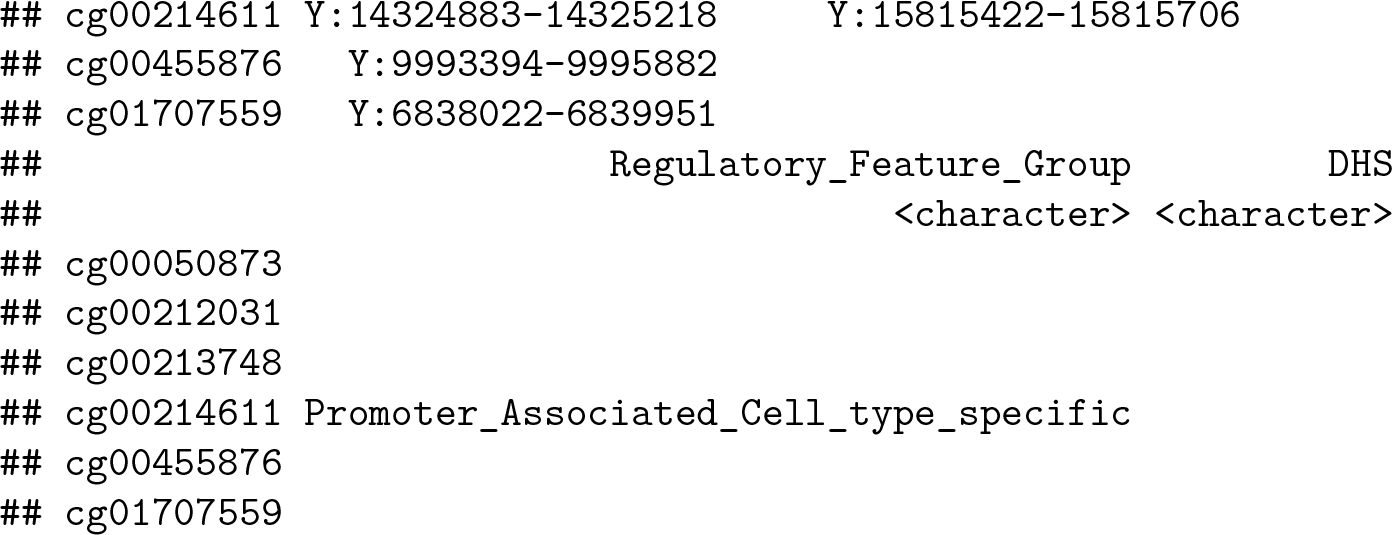

The simplest way to import the raw methylation data into R is using the *minfi* function **`read.450k.sheet`**, along with the path to the IDAT files and a sample sheet. The sample sheet is a CSV (comma-separated) file containing one line per sample, with a number of columns describing each sample. The format expected by the **`read.450k.sheet`** function is based on the sample sheet file that usually accompanies Illumina methylation array data. It is also very similar to the targets file described by the *limma* package. Importing the sample sheet into R creates a **`data.frame`** with one row for each sample and several columns. The **`read.450k.sheet`** function uses the specified path and other information from the sample sheet to create a column called Basename which specifies the location of each individual IDAT file in the experiment.

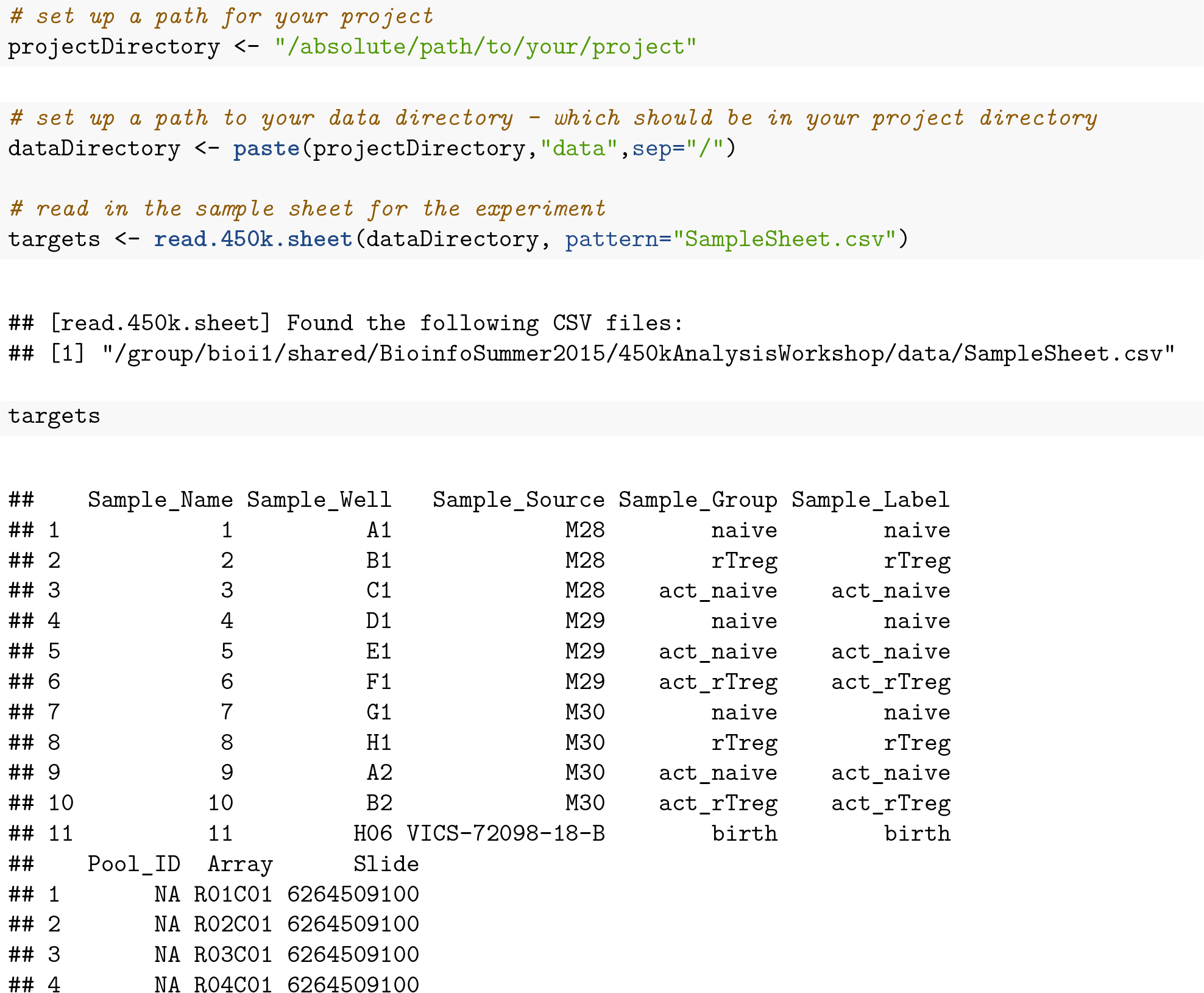

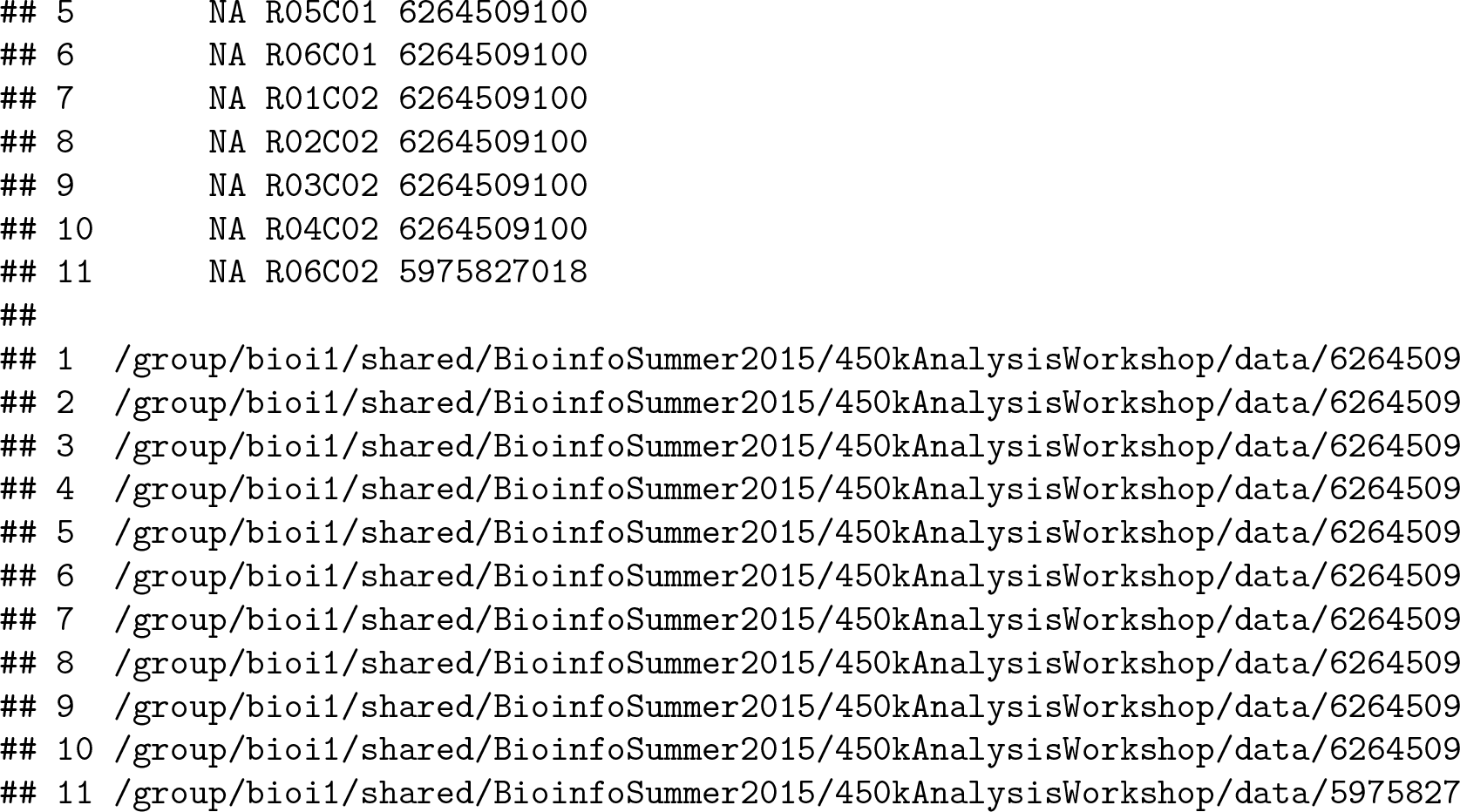

Now that we have imported the information about the samples and where the data is located, we can import the raw intensity signals into R from the IDAT files. This creates an **`RGChannelSet`** object that contains all the raw intensity data, from both the red and green colour channels, for each of the samples. At this stage, it can be useful to rename the samples with more descriptive names.

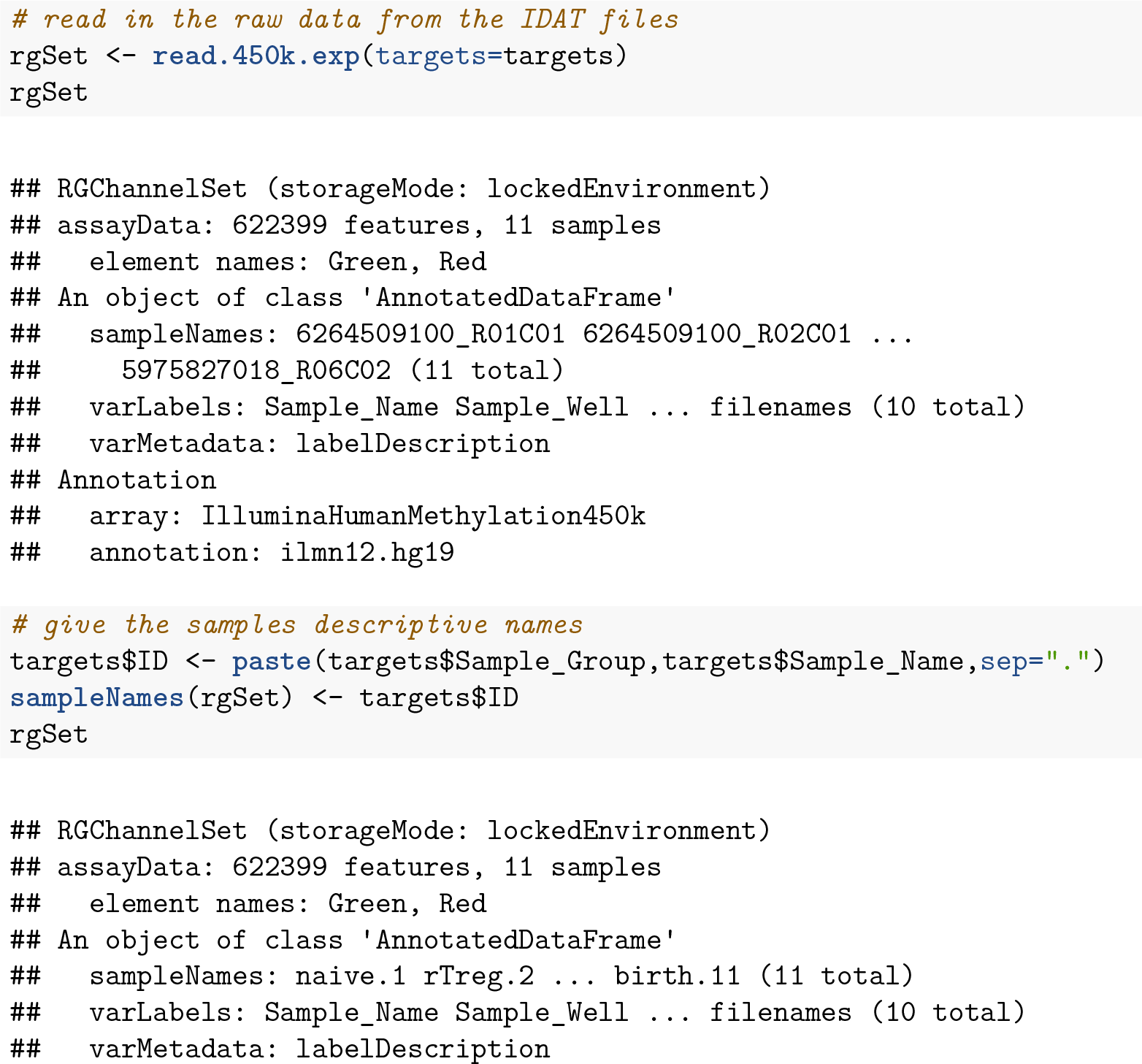

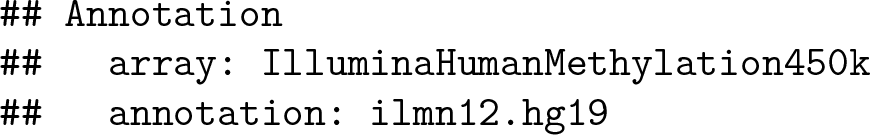

## Quality Control

Once the data has been imported into R, we can evaluate its quality. Firstly, we need to calculate detection p-values. We can generate a detection p-value for every CpG in every sample, which is indicative of the quality of the signal. The method used by *minfi* to calculate detection p-values compares the total signal *(M + U*) for each probe to the background signal level, which is estimated from the negative control probes. Very small p-values are indicative of a reliable signal whilst large p-values, for example >0.01, generally indicate a poor quality signal.

Plotting the mean detection p-value for each sample allows us to gauge the general quality of the samples in terms of the overall signal reliability. Samples that have many failed probes will have relatively large mean detection p-values.

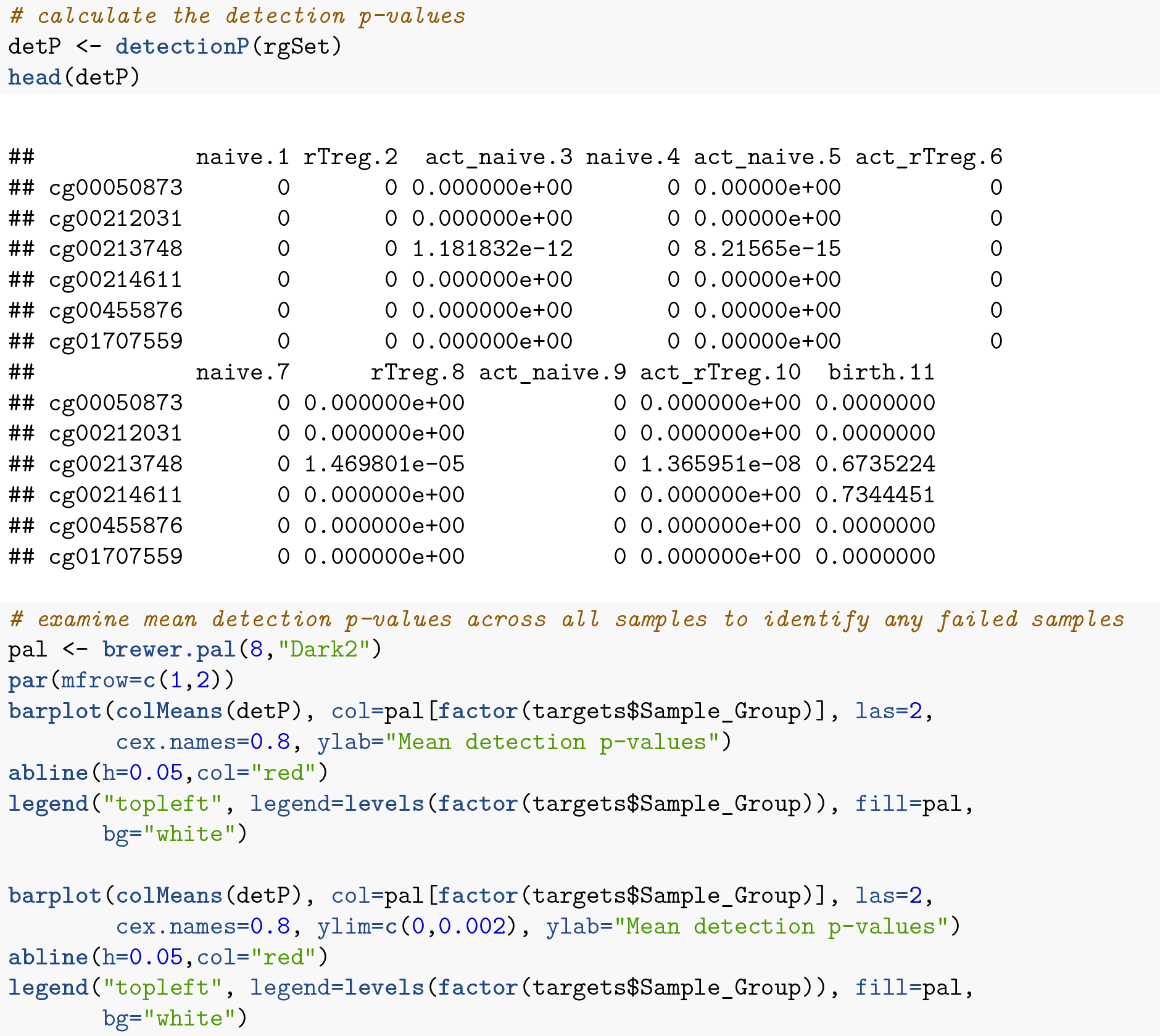

The *minfi* **`qcReport`** function generates many other useful quality control plots. The *minfi* vignette describes the various plots and how they should be interpreted in detail. Generally, samples that look poor based on mean detection p-value will also look poor using other metrics and it is usually advisable to exclude them from further analysis.

**Figure 2:**
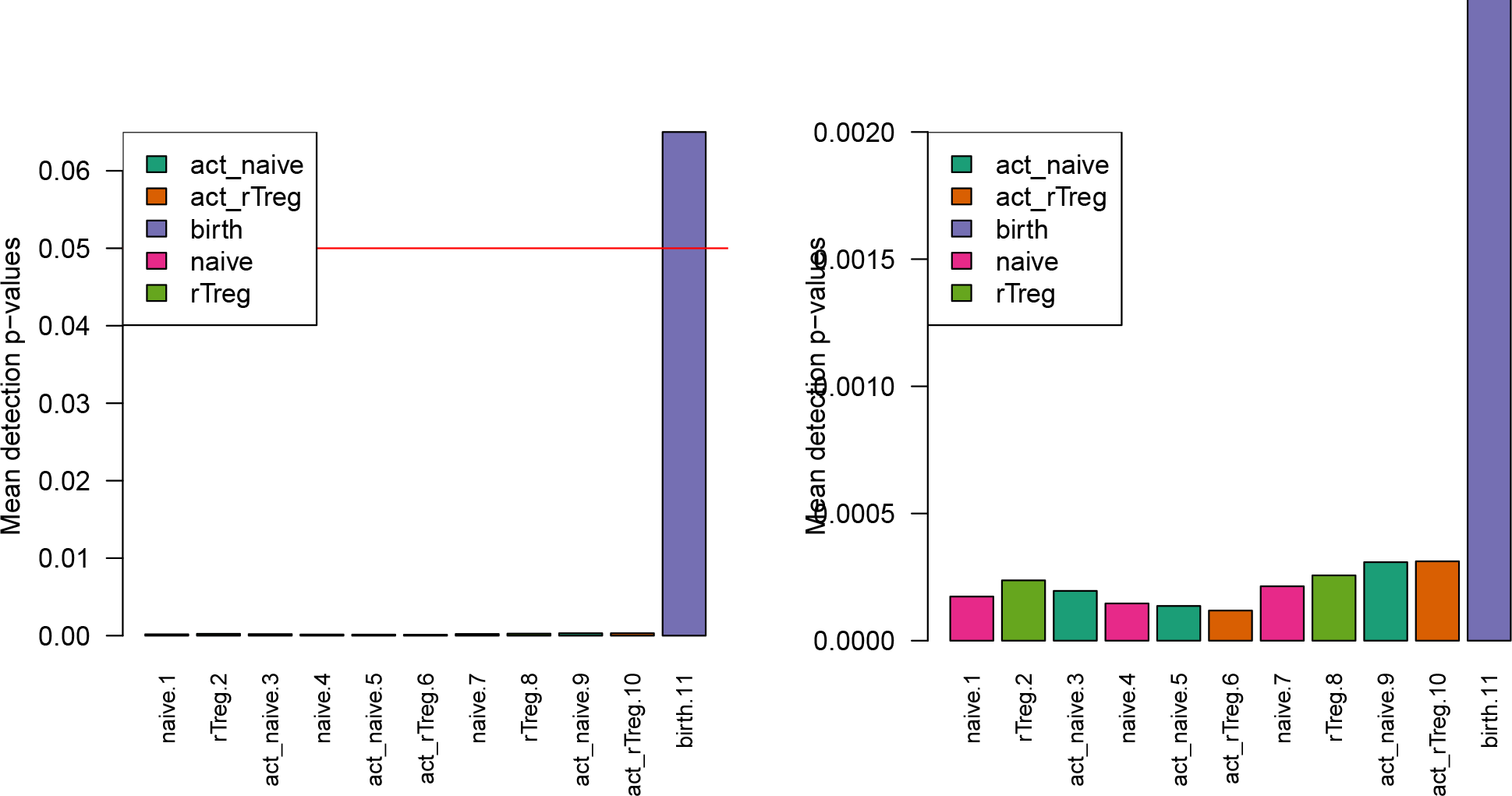
Mean detection p-values summarise the quality of the signal across all the probes in each sample.

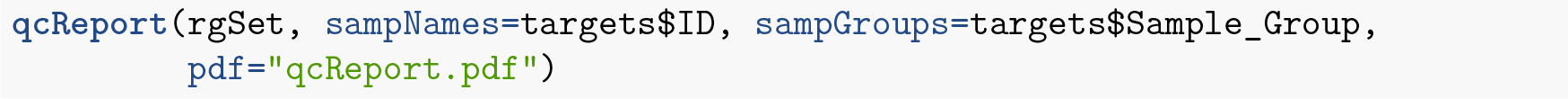

Poor quality samples can be easily excluded from the analysis using a detection p-value cutoff, for example >0.05. For this particular dataset, the **`birth`** sample shows a very high mean detection p-value, and hence it is excluded from subsequent analysis.

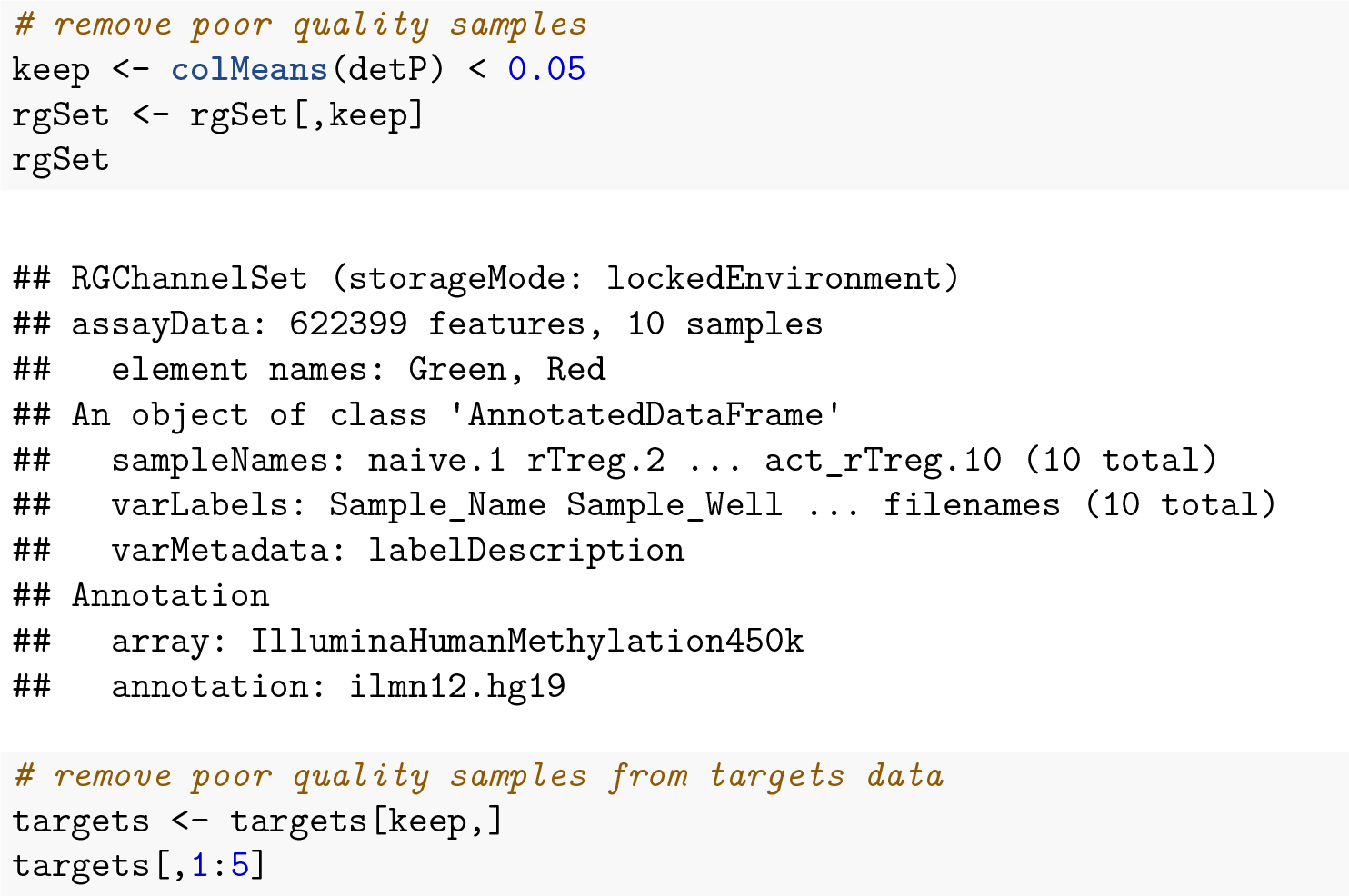

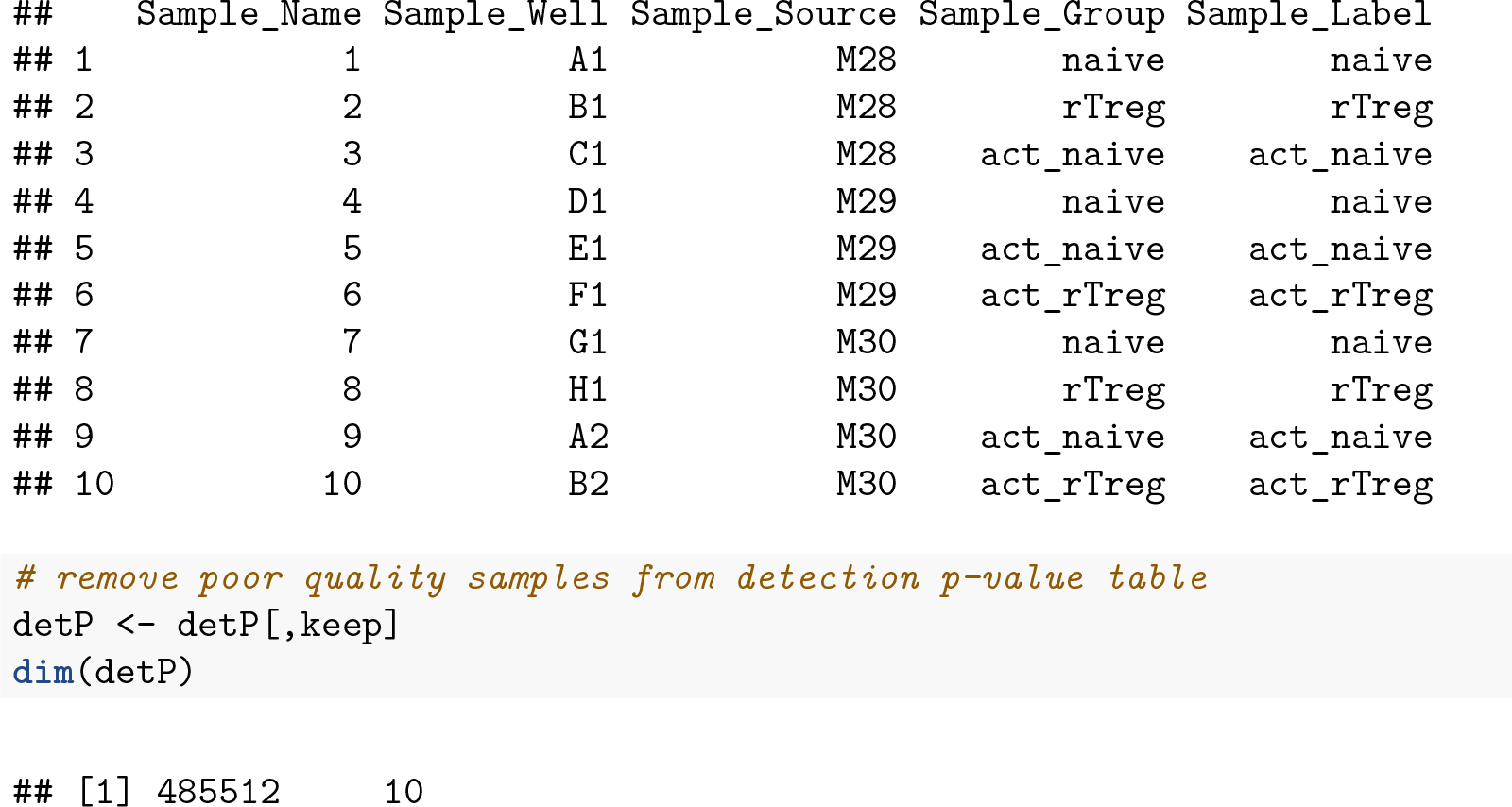

## Normalization

To minimise the unwanted variation within and between samples, various data normalizations can be applied. Many different types of normalization have been developed for methylation arrays and it is beyond the scope of this workflow to compare and contrast all of them (J. Fortin et al. 2014; M. C. Wu et al. 2014; Sun et al. 2011; D. Wang et al. 2012; Maksimovic, Gordon, and Oshlack 2012; Mancuso et al. 2011; Touleimat and Tost 2012; Teschendorff et al. 2013; Pidsley et al. 2013; T. J. Triche et al. 2013). Several methods have been built into *minfi* and can be directly applied within its framework (J. Fortin et al. 2014; T. J. Triche et al. 2013; Maksimovic, Gordon, and Oshlack 2012; Touleimat and Tost 2012), whilst others are *methylumi-specific* or require custom data types (M. C. Wu et al. 2014; Sun et al. 2011; D. Wang et al. 2012; Mancuso et al. 2011; Teschendorff et al. 2013; Pidsley et al. 2013). Although there is no single normalisation method that is universally considered best, a recent study by J. Fortin et al. (2014) has suggested that a good rule of thumb within the *minfi* framework is that the **`preprocessFunnorm`** (J. Fortin et al. 2014) function is most appropriate for datasets with global methylation differences such as cancer/normal or vastly different tissue types, whilst the **`preprocessQuantile`** function (Touleimat and Tost 2012) is more suited for datasets where you do not expect global differences between your samples, for example a single tissue. As we are comparing different blood cell types, which are globally relatively similar, we will apply the **`preprocessQuantile`** method to our data. Note that after normalization, the data is housed in a **`GenomicRatioSet`** object. This is a much more compact representation of the data as the colour channel information has been discarded and the *M* and *U* intensity information has been converted to M-values and beta values, together with associated genomic coordinates.

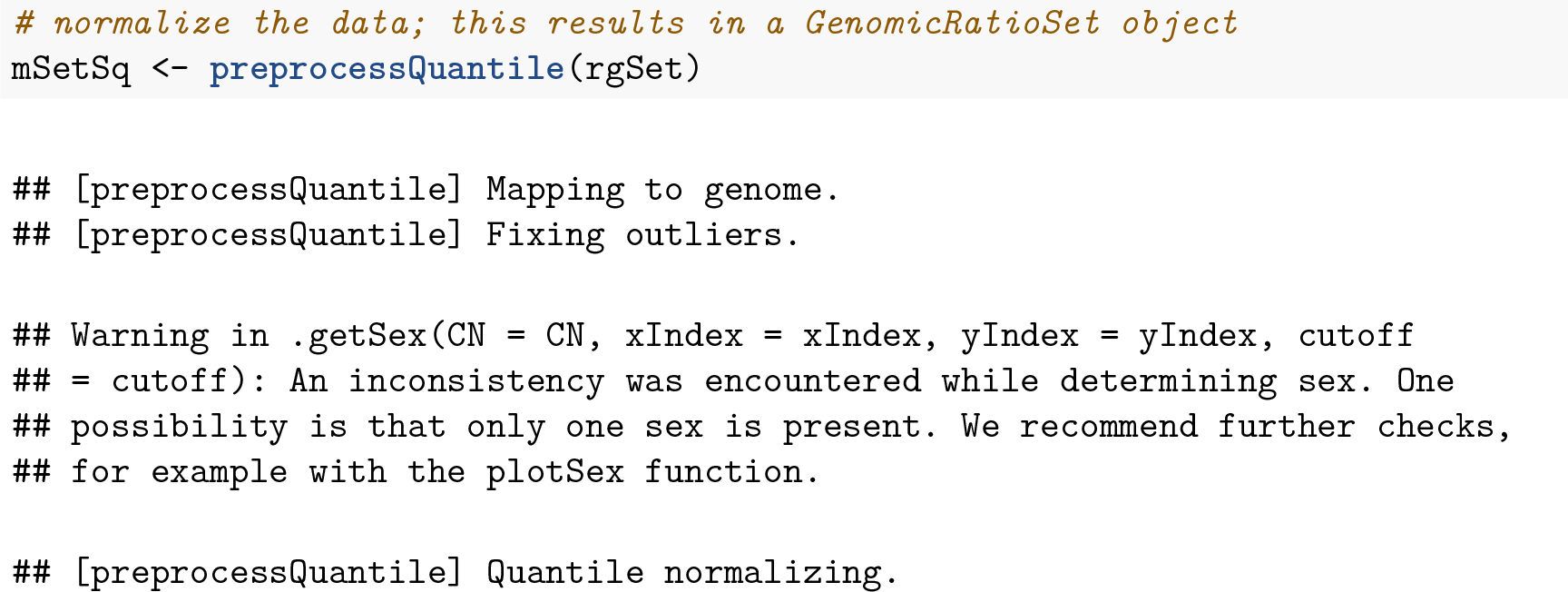

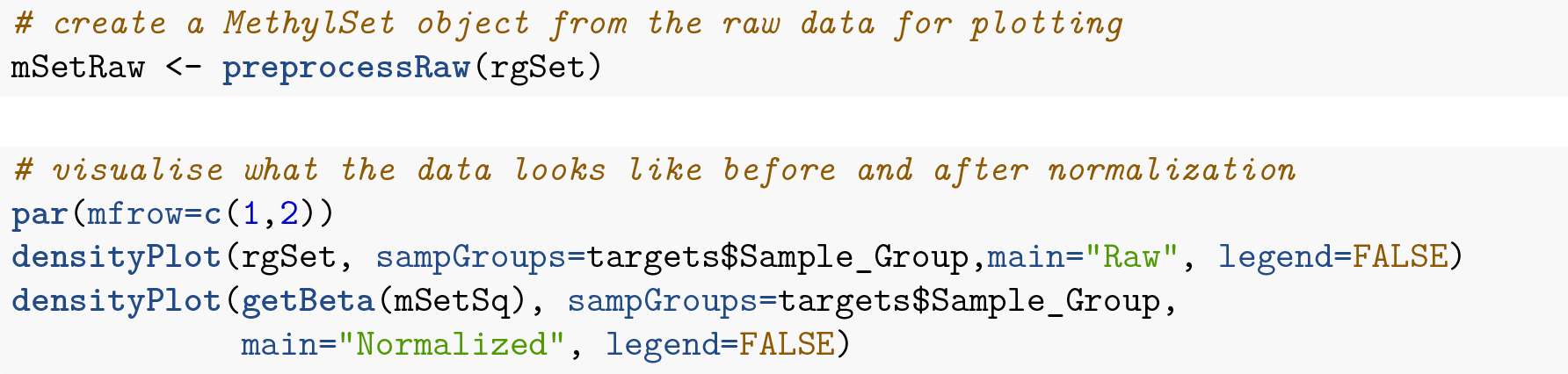

**Figure 3:**
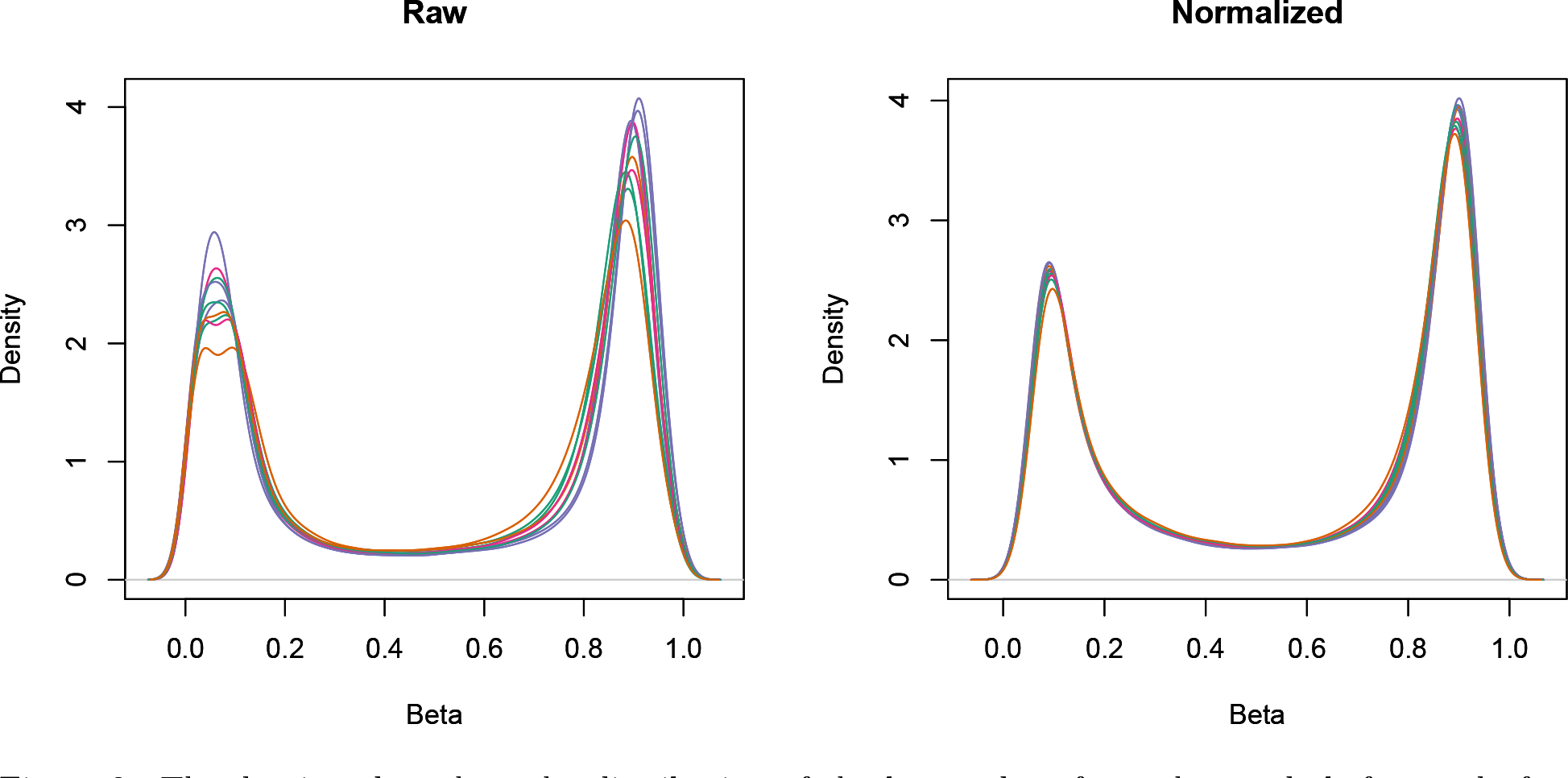
The density plots show the distribution of the beta values for each sample before and after normalization.

## Data Exploration

Multi-dimensional scaling (MDS) plots are excellent for visualising data, and are usually some of the first plots that should be made when exploring the data. MDS plots are based on principle components analysis and are an unsupervised method for looking at the similarities and differences between the various samples. Samples that are more similar to each other should cluster together, and samples that are very different should be further apart on the plot. Dimension one (or principle component one) captures the greatest source of variation in the data, dimension two captures the second greatest source of variation in the data and so on. Colouring the data points or labels by known factors of interest can often highlight exactly what the greatest sources of variation are in the data. It is also possible to use MDS plots to decipher sample mix-ups.

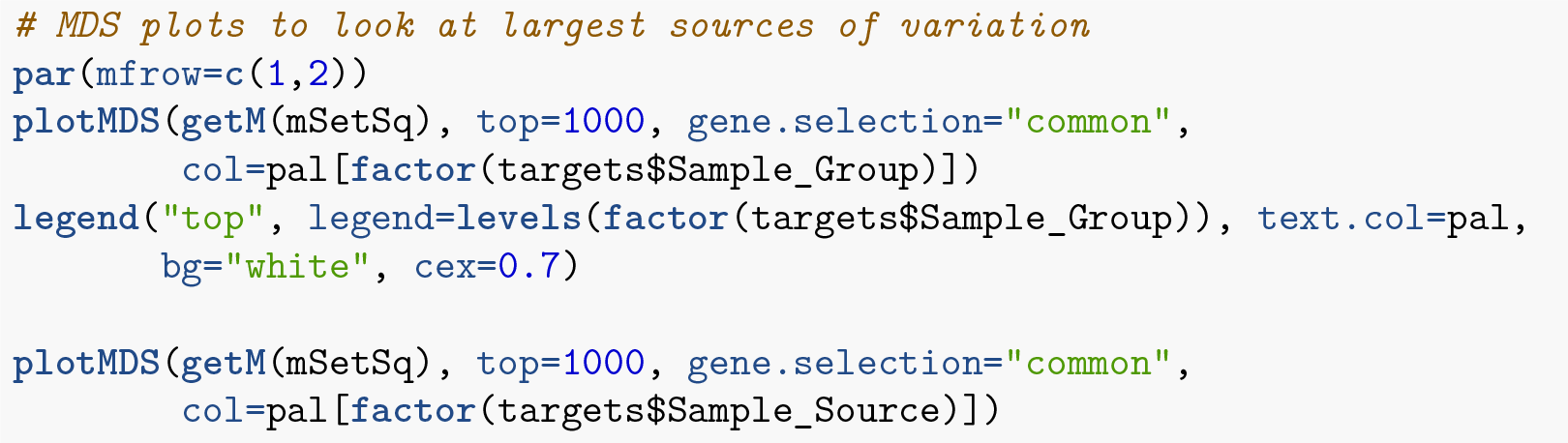

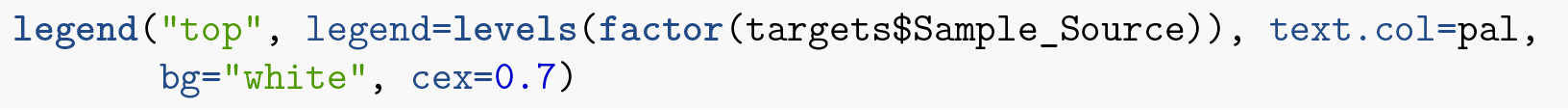

**Figure 4:**
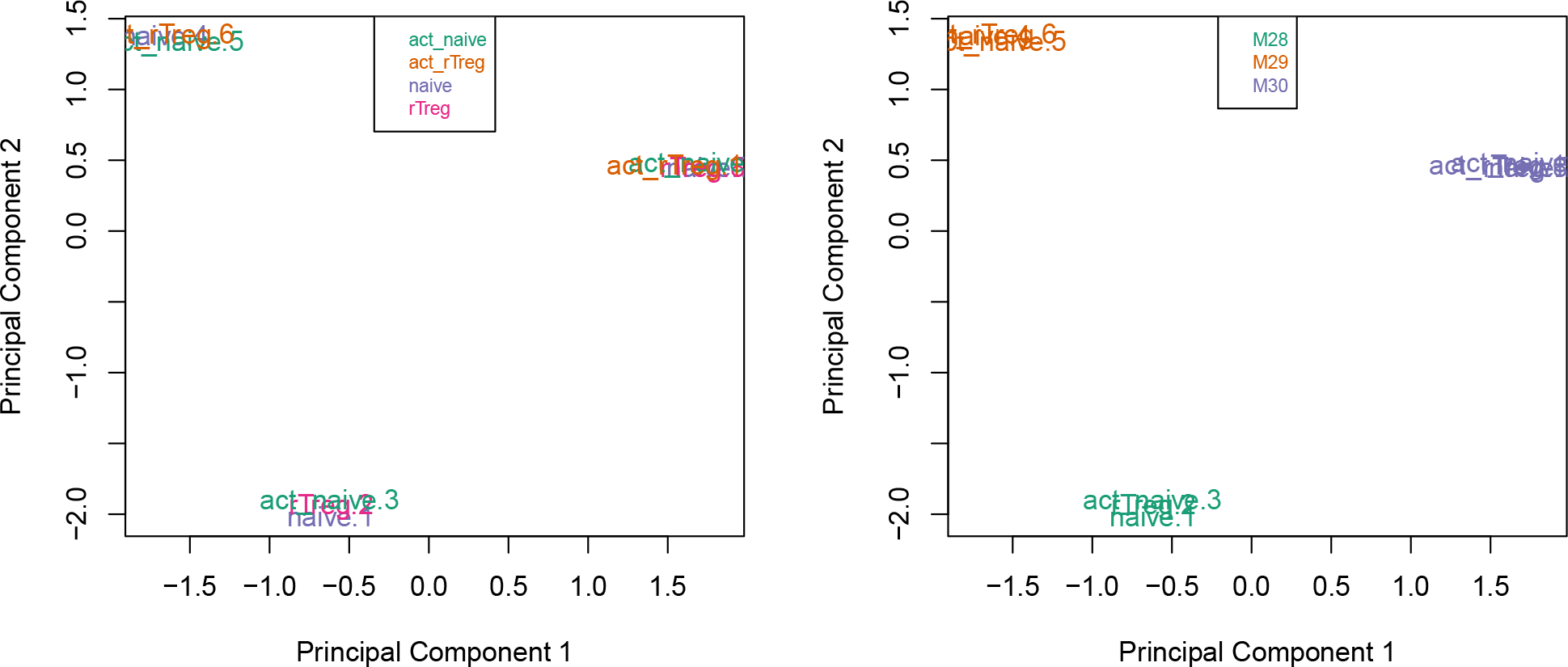
Multi-dimensional scaling plots are a good way to visualise the relationships between the samples in an experiment.

Examining the MDS plots for this dataset demonstrates that the largest source of variation is the difference between individuals. The higher dimensions reveal that the differences between cell types are largely captured by the third and fourth principal components. This type of information is useful in that it can inform downstream analysis. If obvious sources of unwanted variation are revealed by the MDS plots, we can include them in our statistical model to account for them. In the case of this particular dataset, we will include individual to individual variation in our statistical model.

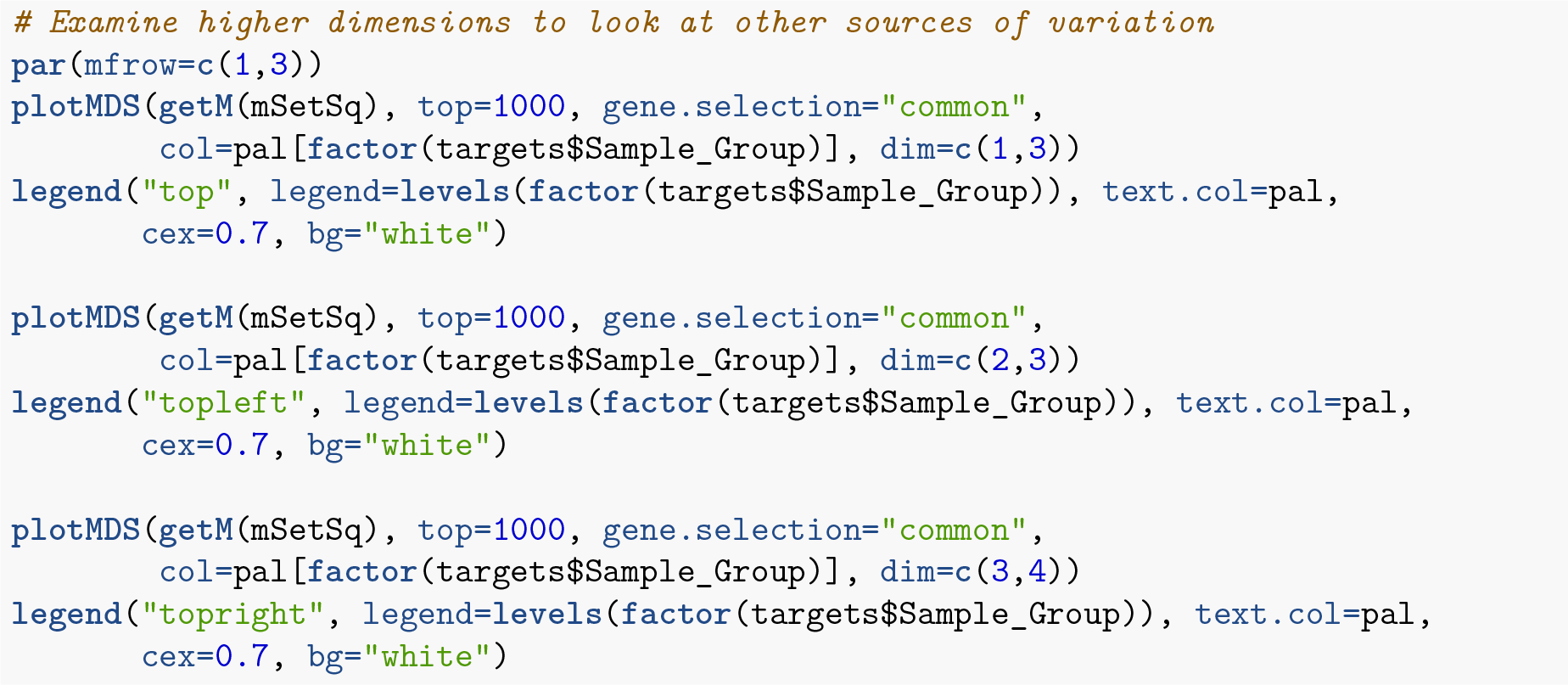

## Filtering

Poor performing probes are generally filtered out prior to differential methylation analysis. As the signal from these probes is unreliable, by removing them we perform fewer statistical tests and thus incur a reduced multiple testing penalty. We filter out probes that have failed in one or more samples based on detection p-value.

**Figure 5:**
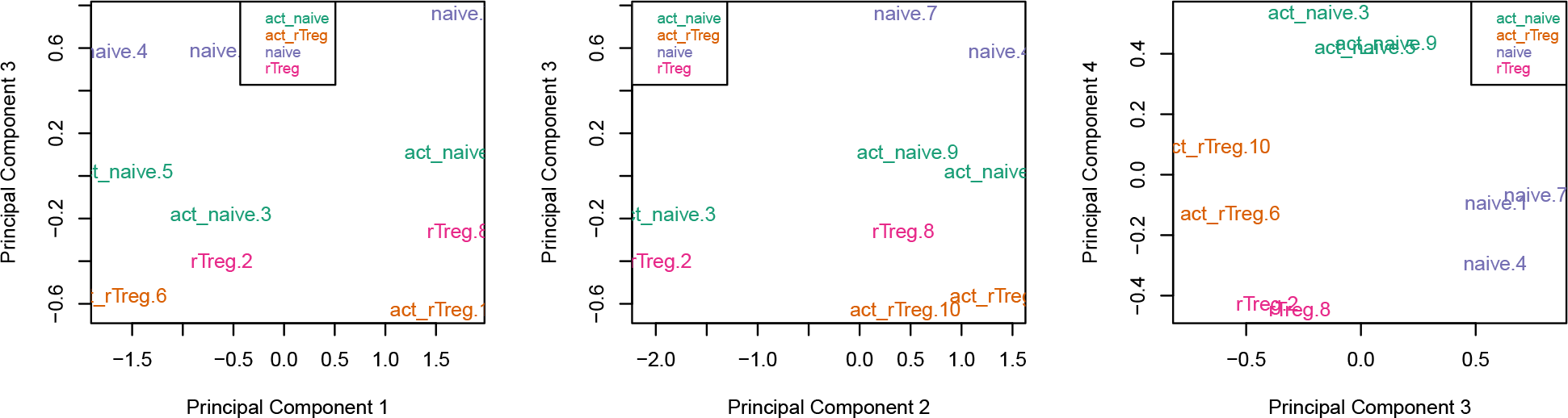
Examining the higher dimensions of an MDS plot can reaveal significant sources of variation in the data.

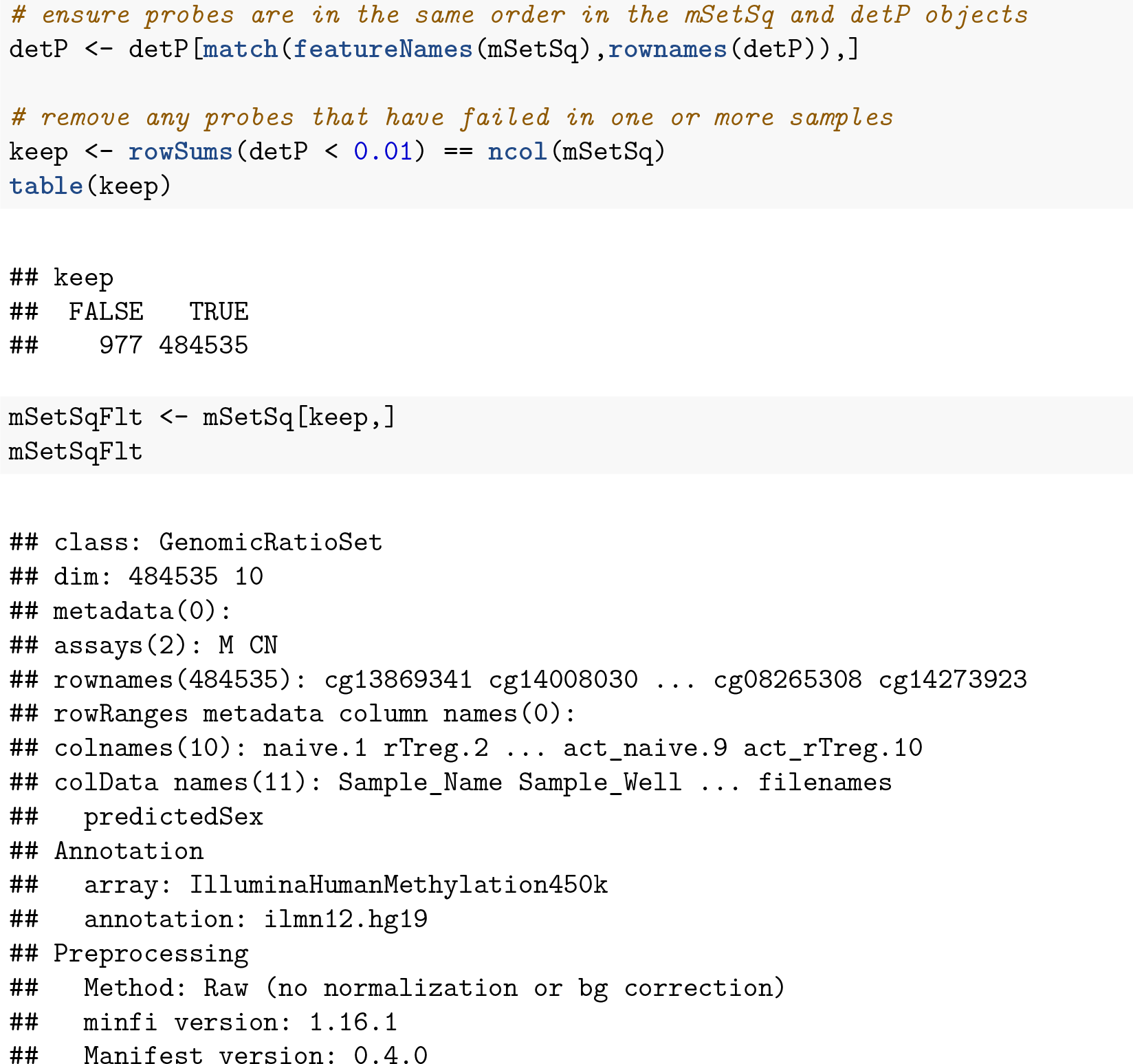

Depending on the nature of your samples and your biological question you may also choose to filter out the probes from the X and Y chromosomes or probes that are known to have common SNPs at the CpG site. As the samples in this dataset were all derived from male donors, we will not be removing the sex chromosome probes as part of this analysis, however example code is provided below. A different dataset, which contains both male and female samples, is used to demonstrate a Differential Variability analysis and provides an example of when sex chromosome removal is necessary.

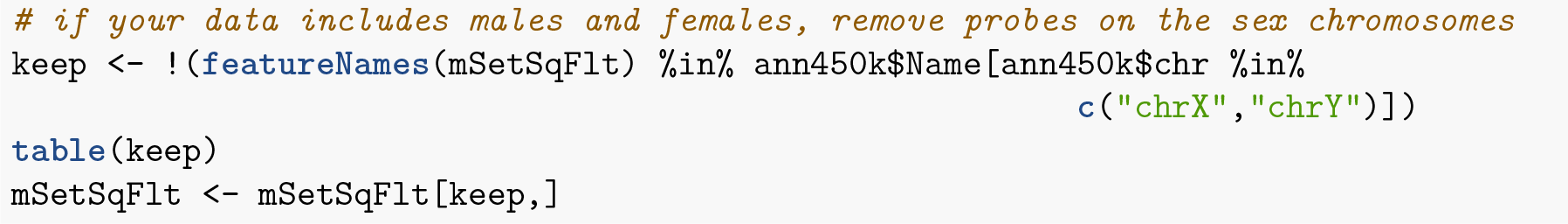
Box initially at rest on sled sliding across ice.

There is a function in *minfi* that provides a simple interface for the removal of probes where common SNPs may affect the CpG. You can either remove all probes affected by SNPs (default), or only those with minor allele frequencies greater than a specified value.

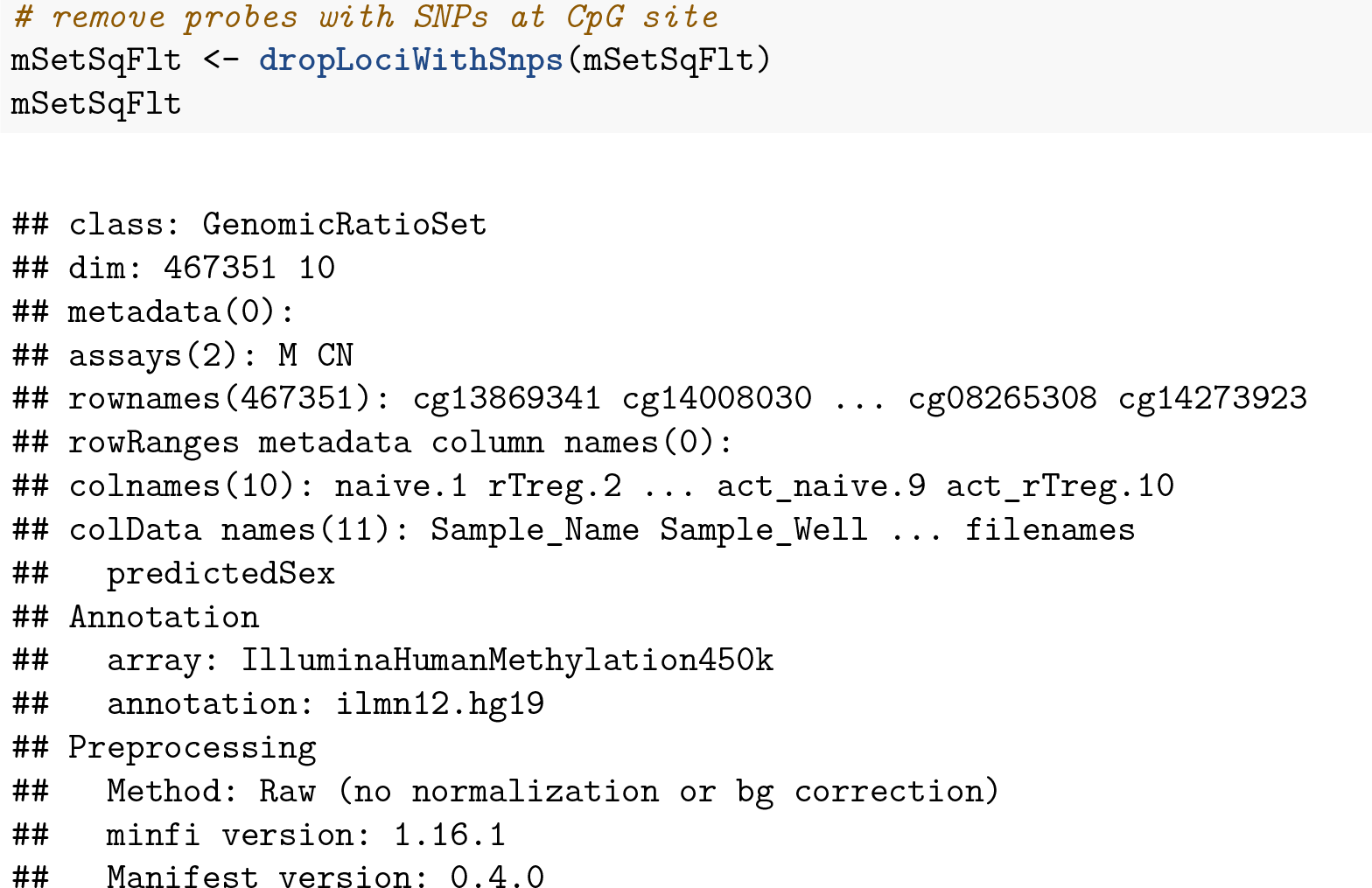

We will also filter out probes that have shown to be cross-reactive, that is, probes that have been demonstrated to map to multiple places in the genome. This list was originally published by Chen et al. (2013) and can be obtained from the authors’ website.

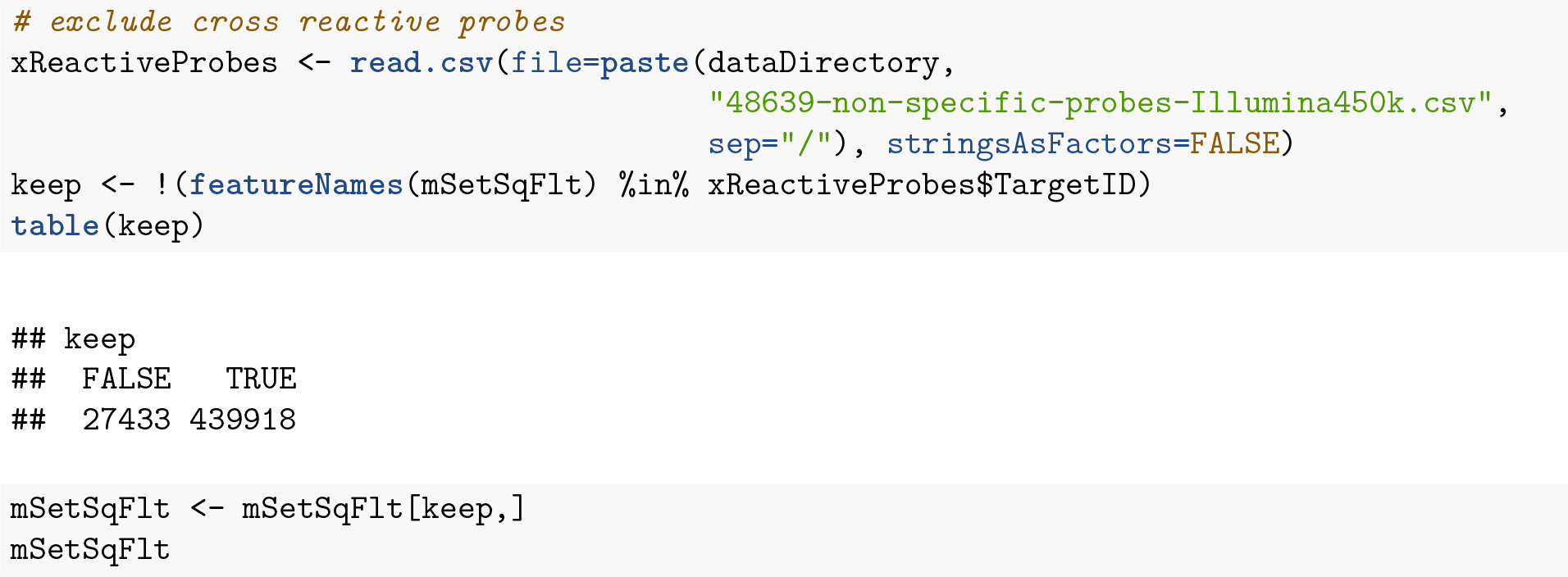

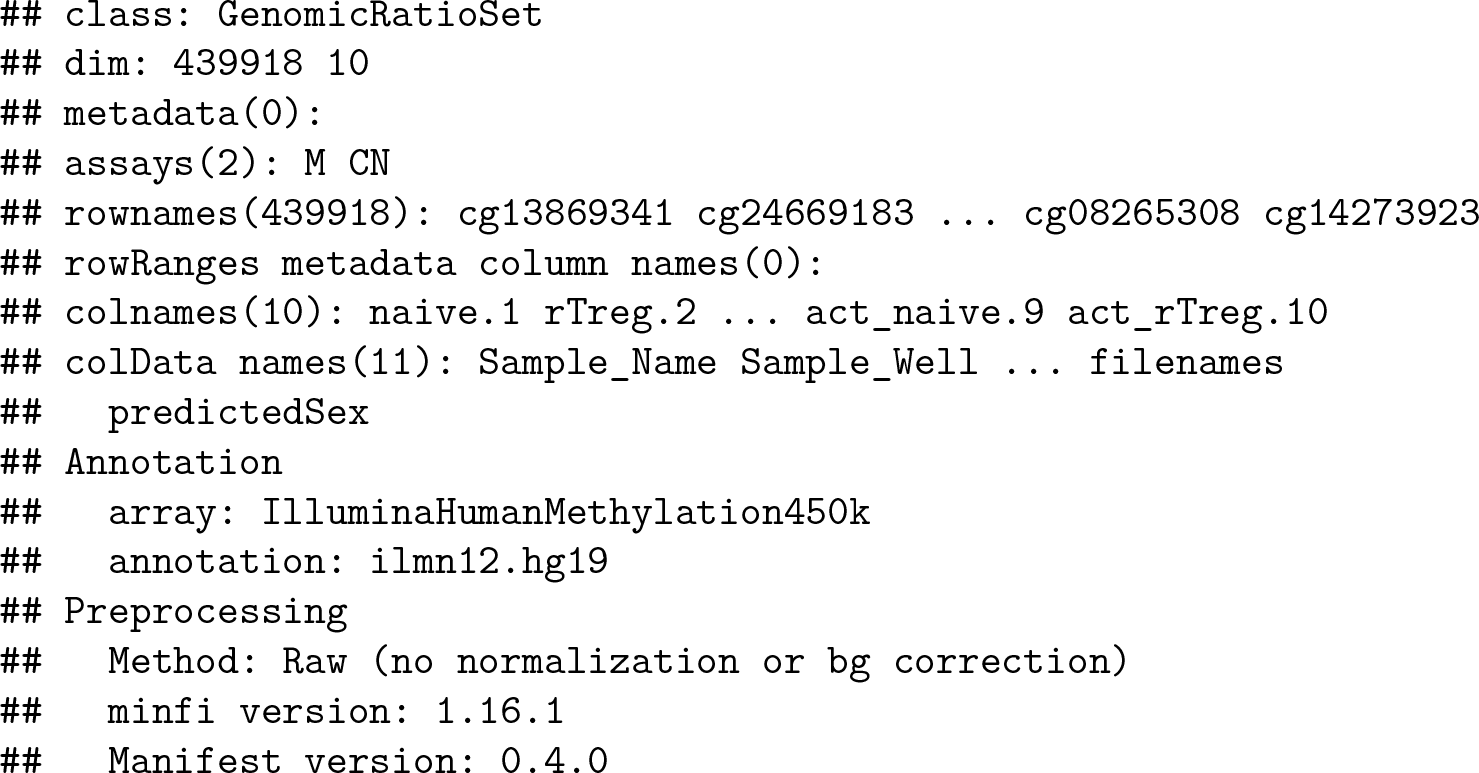

Once the data has been filtered and normalised, it is often useful to re-examine the MDS plots to see if the relationship between the samples has changed. It is apparent from the new MDS plots that much of the inter-individual variation has been removed as this is no longer the first principal component, likely due to the removal of the SNP-affected CpG probes. However, the samples do still cluster by individual in the second dimension and thus a factor for individual should still be included in the model.

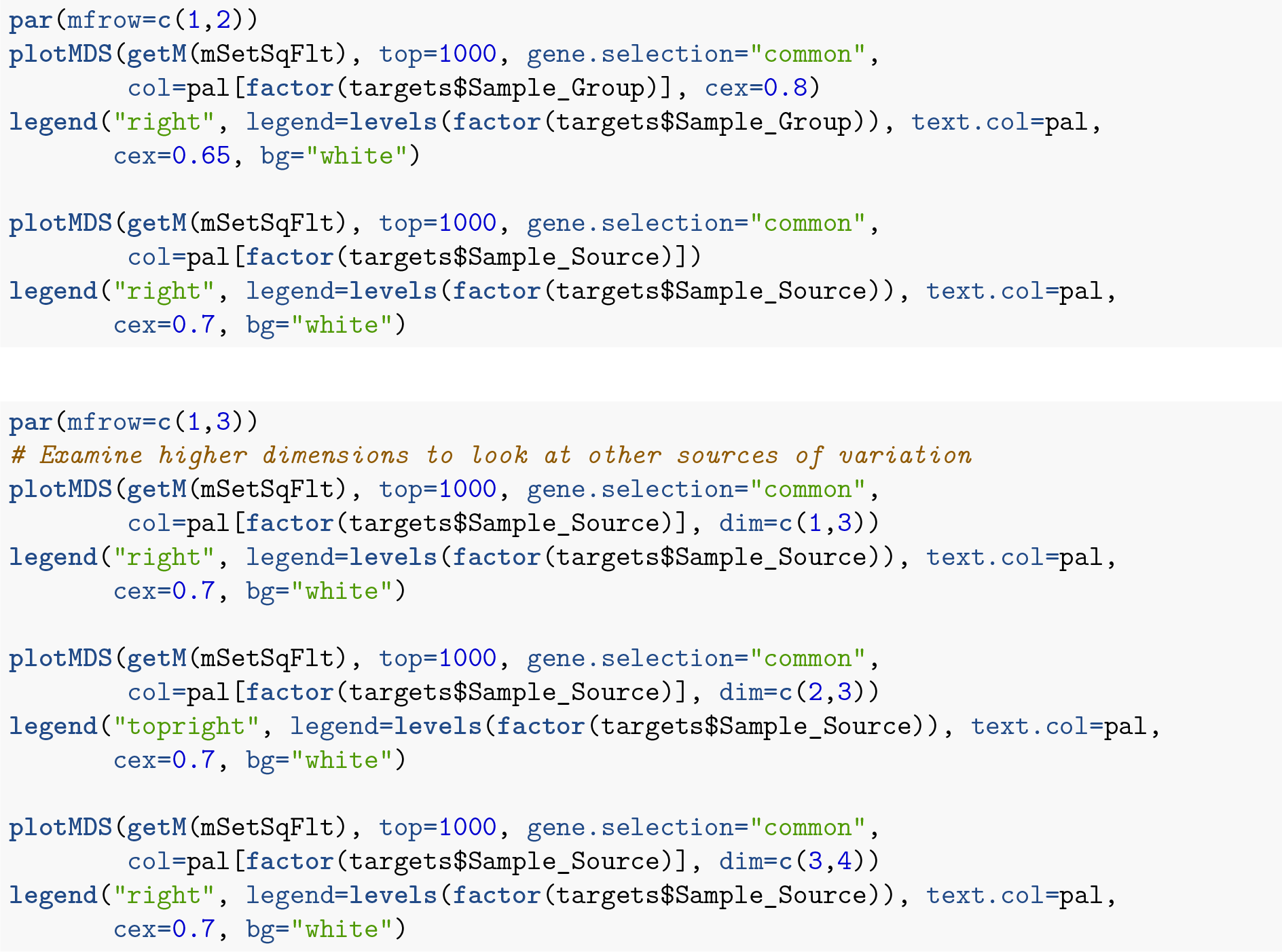

The next step is to calculate M-values and beta values. As previously mentioned, M-values have nicer statistical properties and are thus better for use in statistical analysis of methylation data whilst beta values are easy to interpret and are thus better for displaying data. A detailed comparison of M-values and beta values was published by Du et al. (2010).

**Figure 6:**
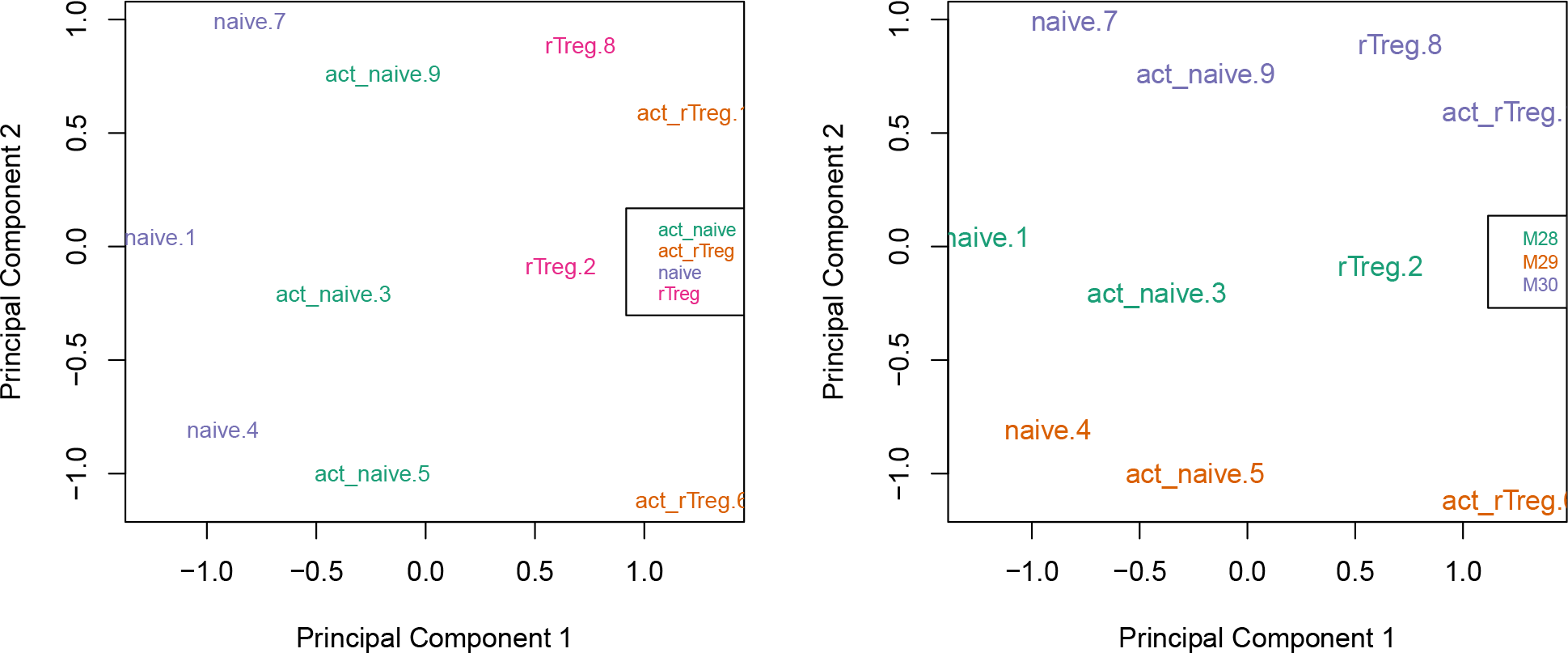
Removing SNP-affected CpGs probes from the data changes the sample clustering in the MDS plots.

**Figure 7:**
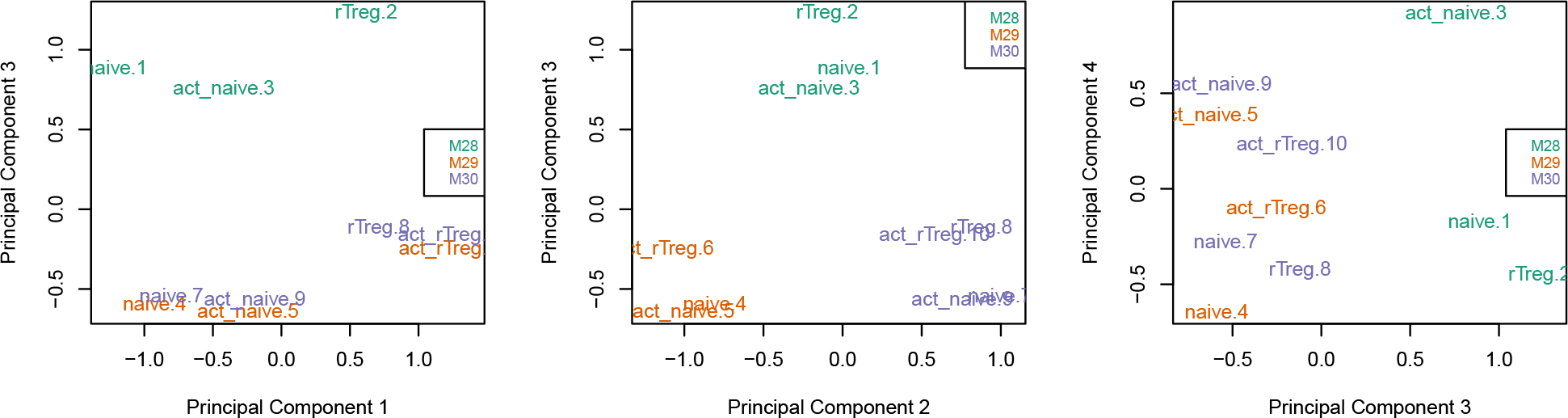
Examining the higher dimensions of the MDS plots shows that significant inter-individual variation still exists in the second and third principle components.

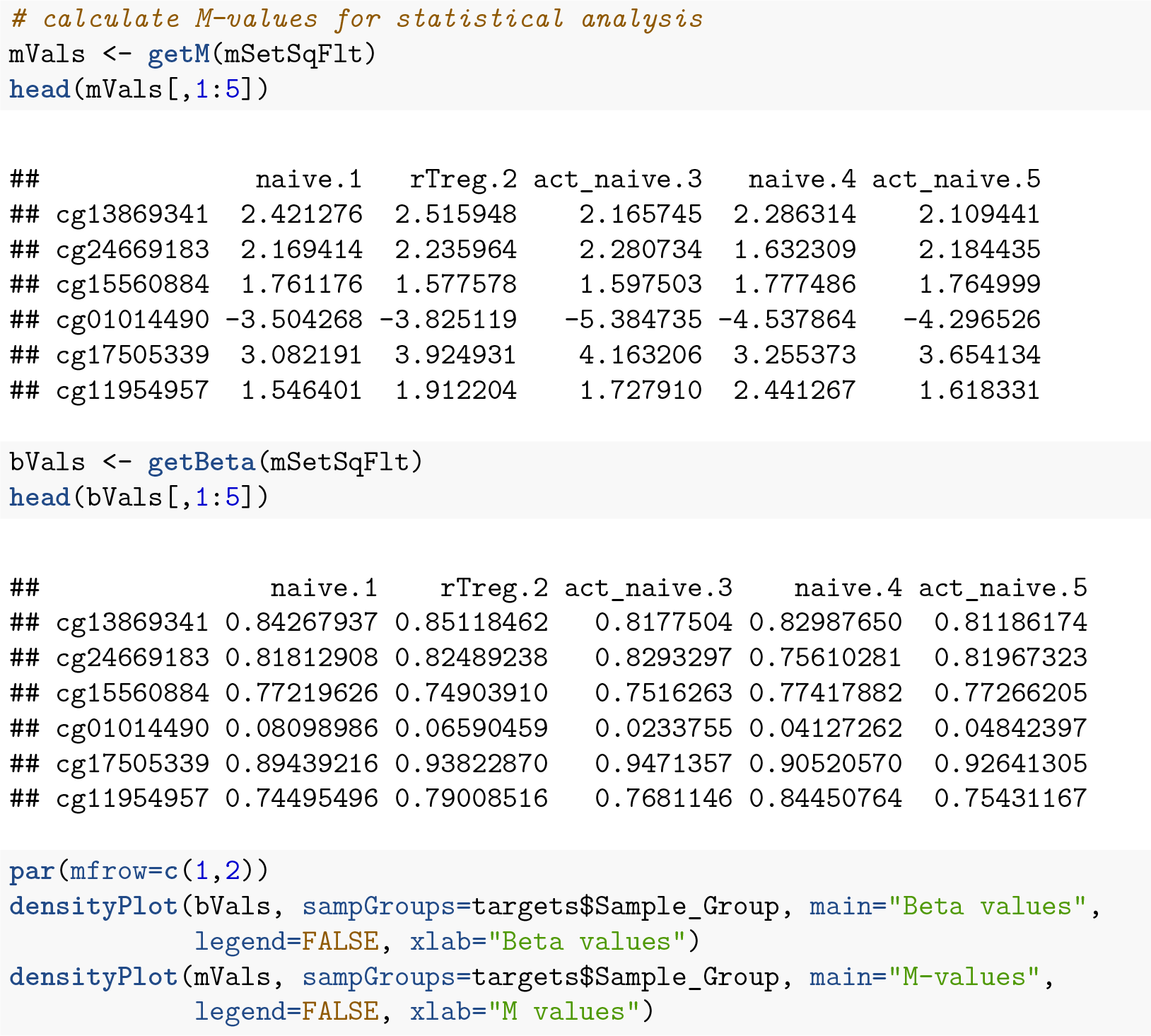

## Probe-wise differential methylation analysis

The biological question of interest for this particular dataset is to discover differentially methylated probes between the different cell types. However, as was apparent in the MDS plots, there is another factor that we need to take into account when we perform the statistical analysis. In the **`targets`** file, there is a column called **`Sample_Source`**, which refers to the individuals that the samples were collected from. In this dataset, each of the individuals contributes more than one cell type. For example, individual M28 contributes **`naive, rTreg and act_naive`** samples. Hence, when we specify our design matrix, we need to include two factors: individual and cell type. This style of analysis is called a paired analysis; differences between cell types are calculated *within* each individual, and then these differences are averaged *across* individuals to determine whether there is an overall significant difference in the mean methylation level for each CpG site. The *limma* User’s Guide extensively covers the different types of designs that are commonly used for microarray experiments and how to analyse them in R.

We are interested in pairwise comparisons between the four cell types, taking into account individual to individual variation. We perform this analysis on the matrix of M-values in *limma*, obtaining moderated t-statistics and associated p-values for each CpG site. The comparison that has the most significantly differentially methylated CpGs is **`naïve`** vs **`rTreg`** (n=3021 at 5% false discovery rate (FDR)), while **`rTreg`** vs **`act_rTreg`** doesn’t show any significant differential methylation.

**Figure 8:**
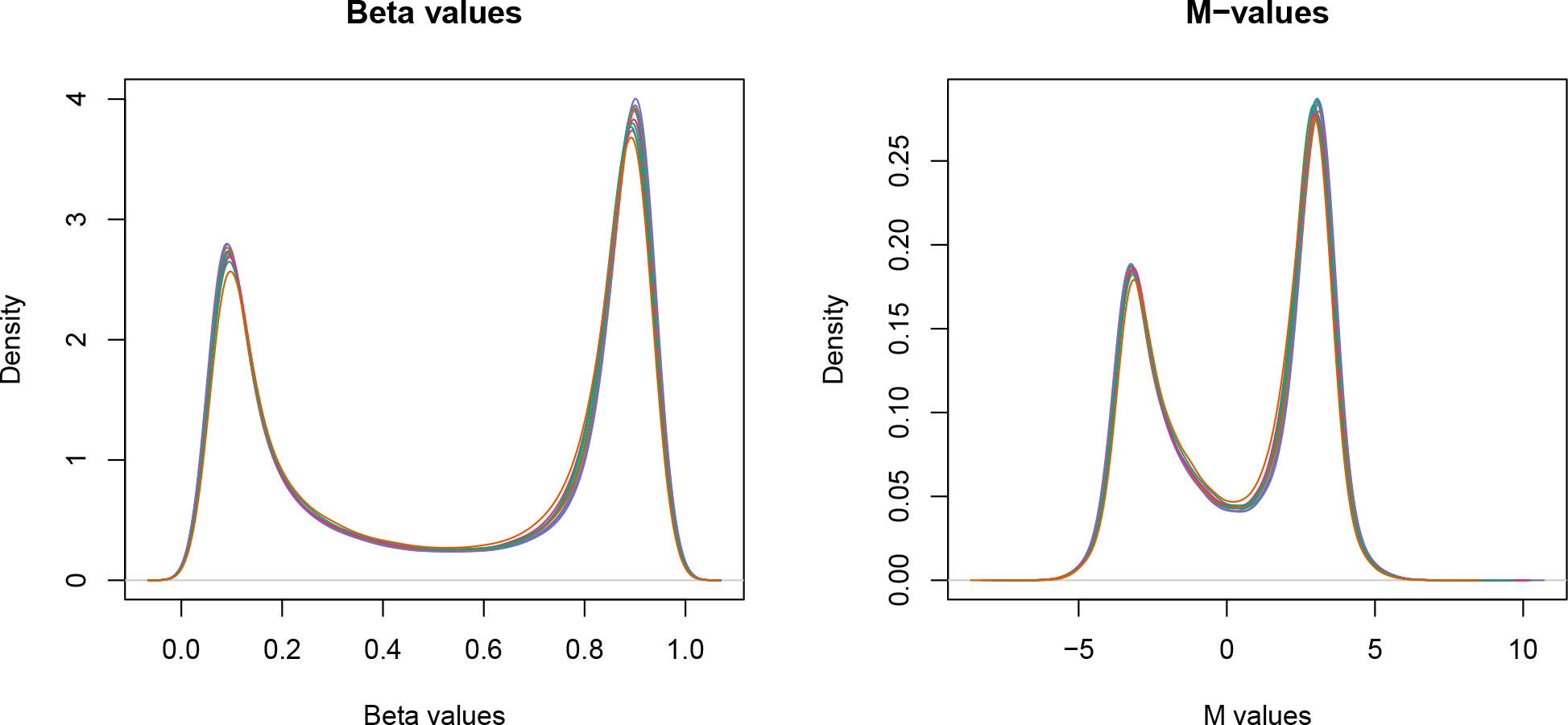
The distributions of beta and M-values are quite different; beta values are constrained between 0 and 1 whilst M-values range between-Inf and Inf.

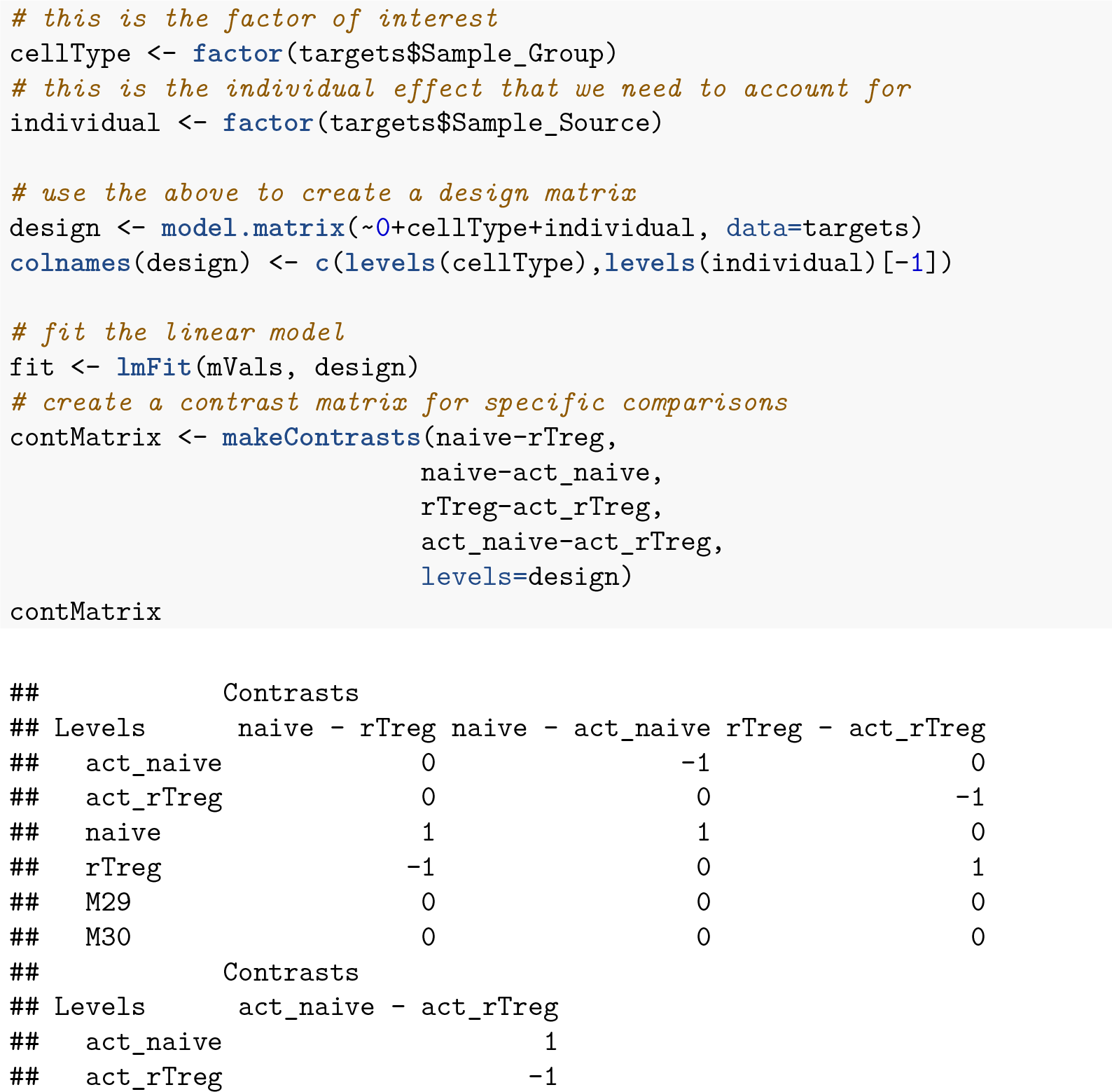

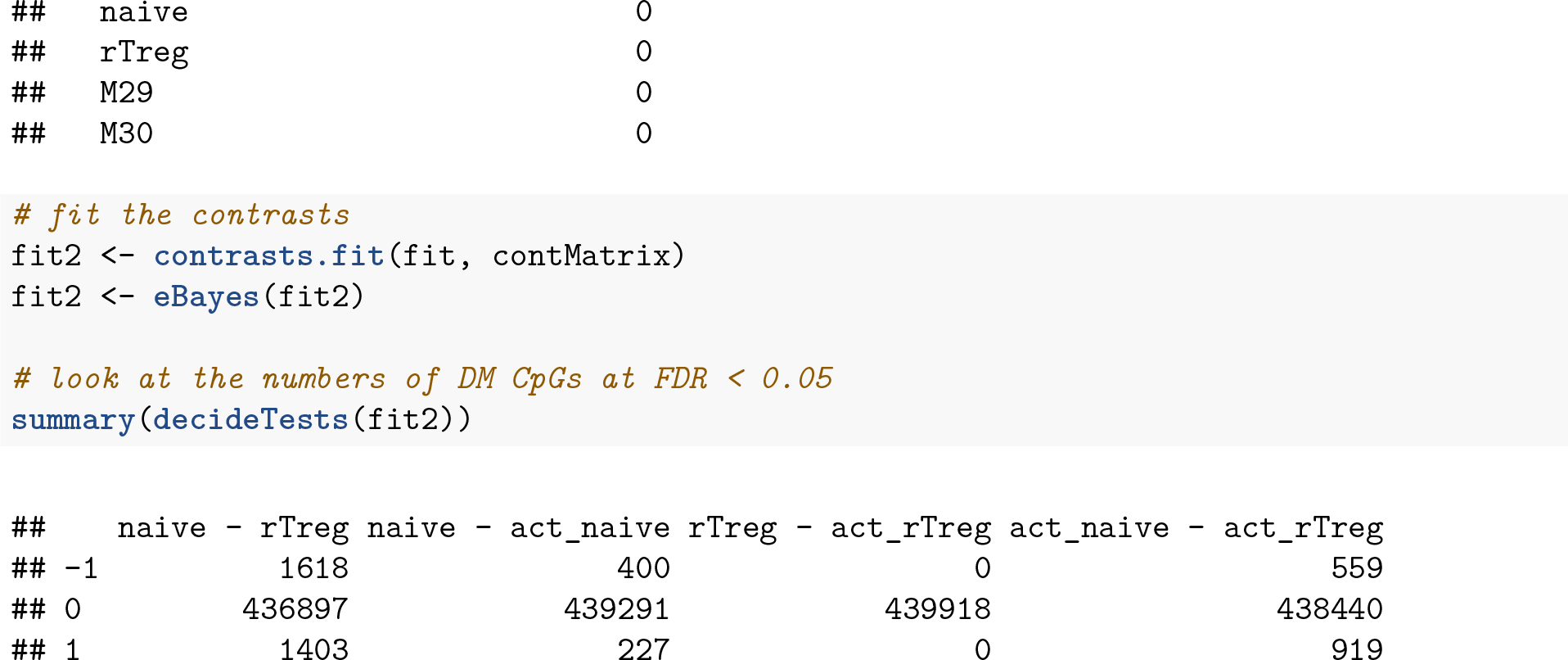

We can extract the tables of differentially expressed CpGs for each comparison, ordered by B-statistic by default, using the **`topTable`** function in *limma*. The results of the analysis for the first comparison, **`naive`** vs. **`rTreg`**, can be saved as a **`data.frame`** by setting **`coef=1`**.

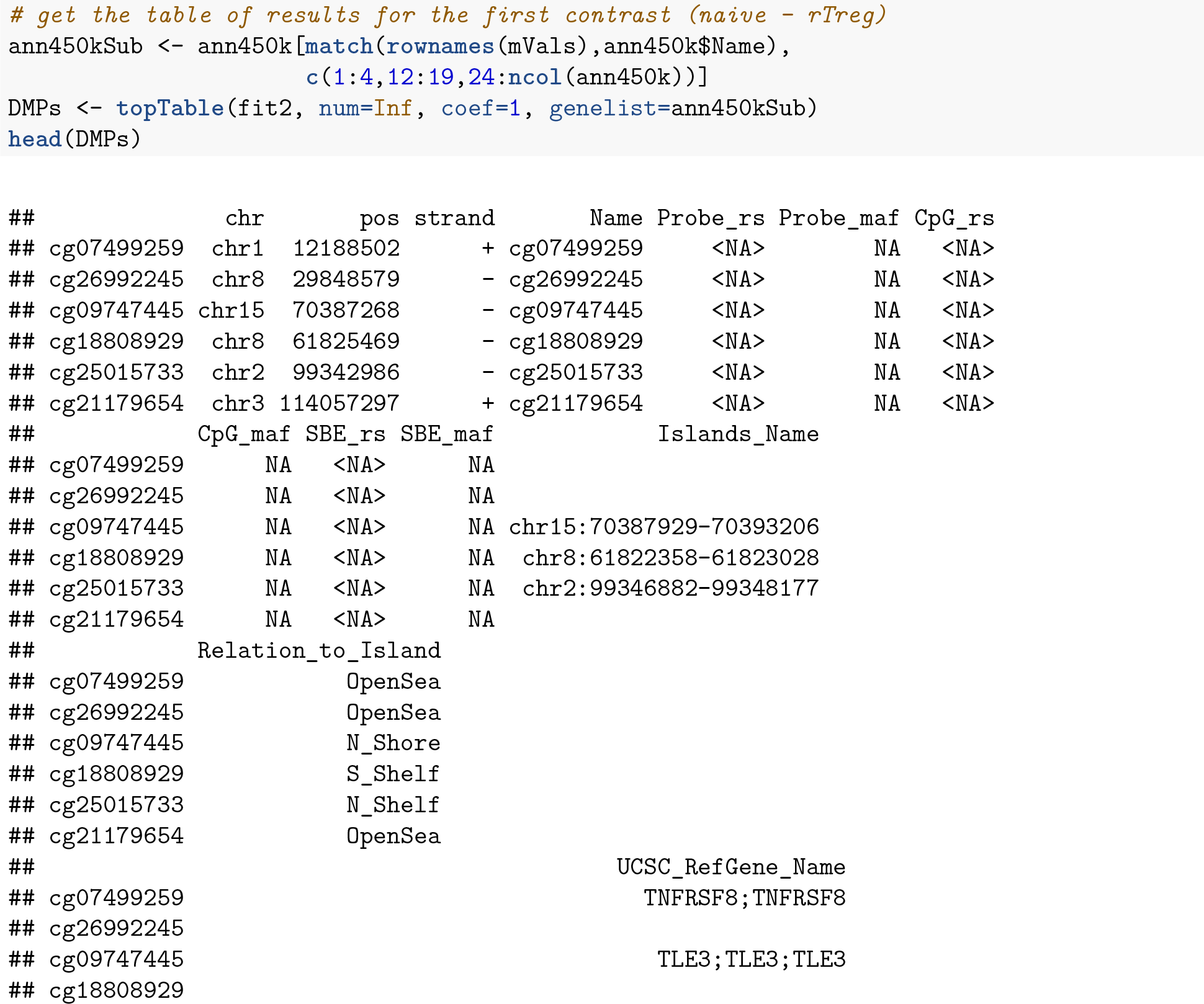

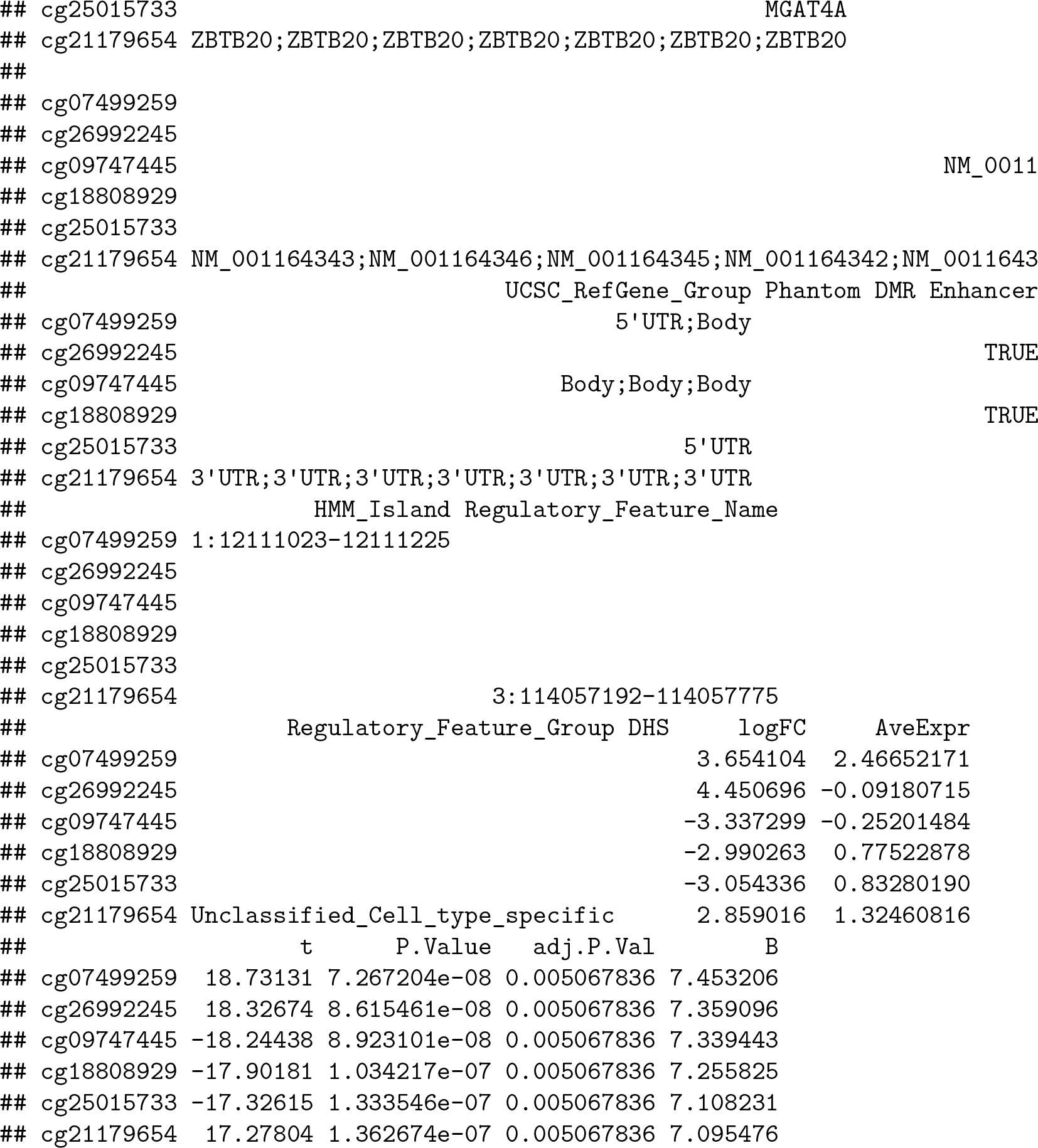

The resulting **`data.frame`** can easily be written to a CSV file, which can be opened in Excel.

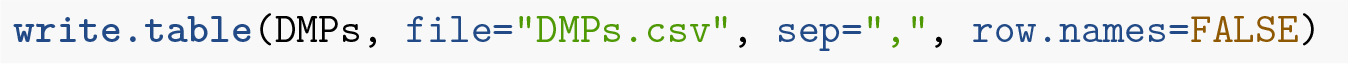

It is always useful to plot sample-wise methylation levels for the top differentially methylated CpG sites to quickly ensure the results make sense. If the plots do not look as expected, it is usually an indication of an error in the code, or in setting up the design matrix. It is easier to interpret methylation levels on the beta value scale, so although the analysis is performed on the M-value scale, we visualise data on the beta value scale. The **`plotCpg`** function in *minfi* is a convenient way to plot the sample-wise beta values stratified by the grouping variable.

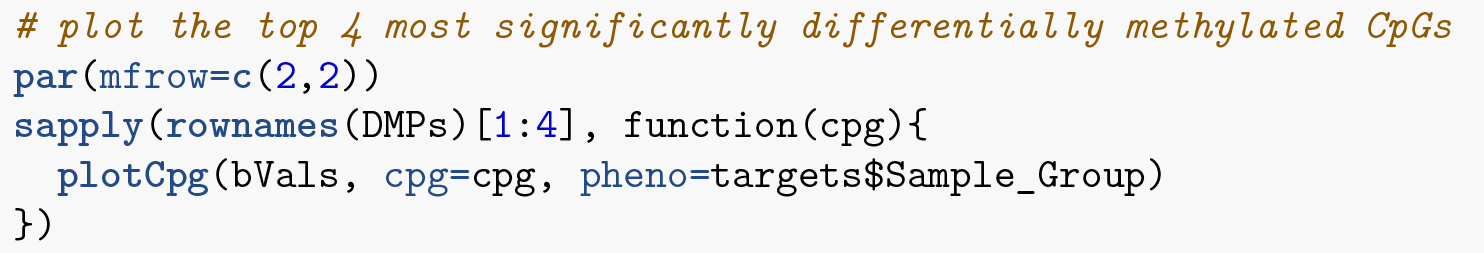

**Figure 9:**
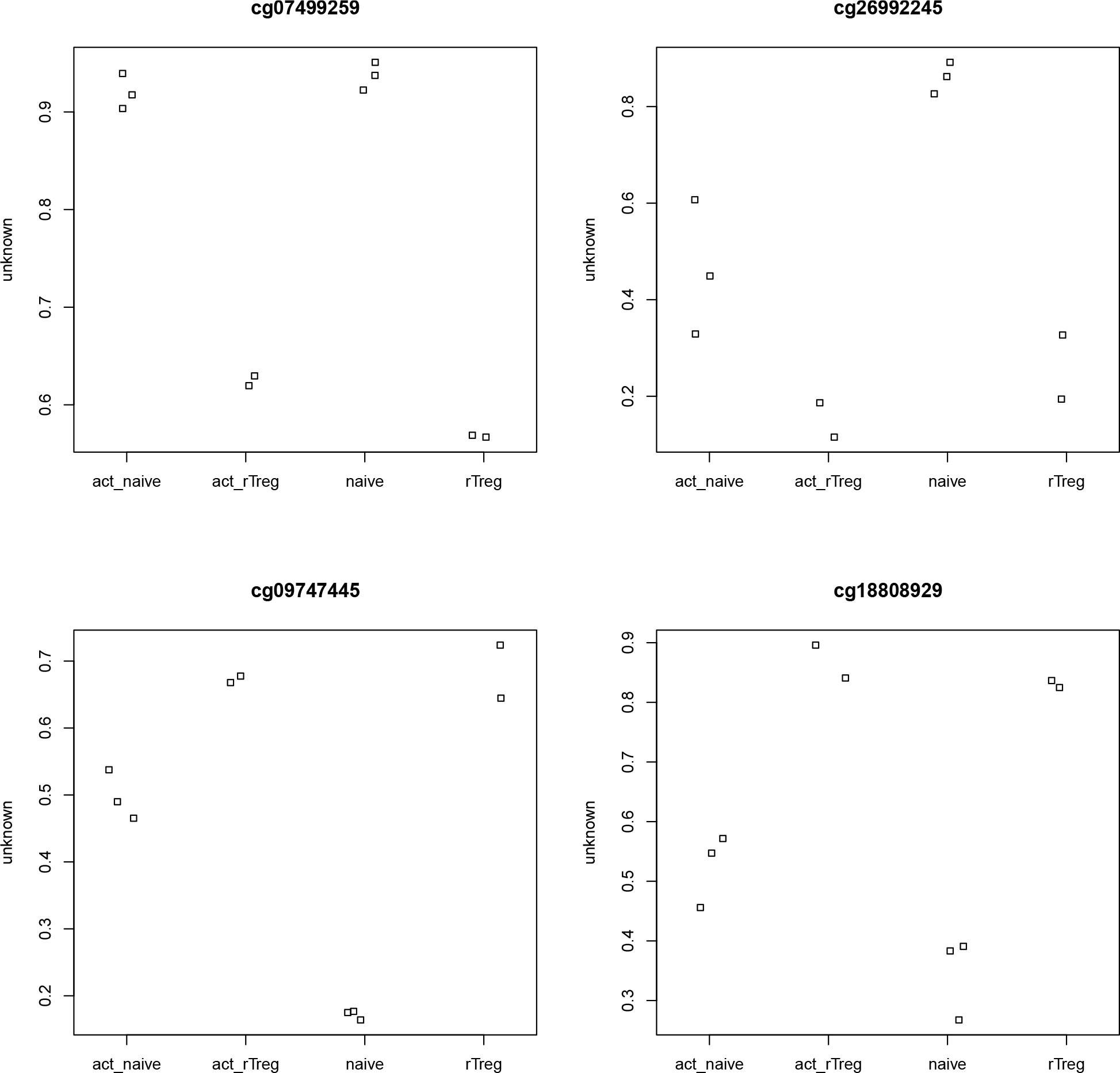
Plotting the top few differentially methylated CpGs is a good way to check whether the results make sense.

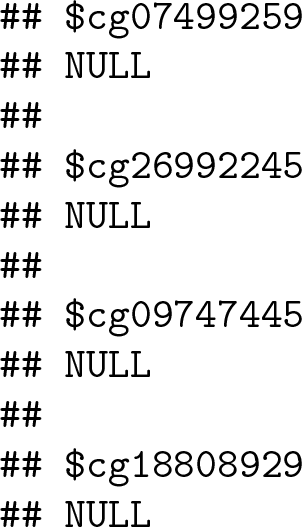

## Differential Methylation Analysis of Regions

Although performing a *probe-wise* analysis is useful and informative, sometimes we are interested in knowing whether several proximal CpGs are concordantly differentially methylated, that is, we want to identify differentially methylated *regions.* There are several Bioconductor packages that have functions for identifying differentially methylated regions from 450k data. Some of the most popular are the **`dmrFind`** function in the charm package, which has been somewhat superseded for 450k arrays by the **`bumphunter`** function in minfi(Jaffe et al. 2012; Aryee et al. 2014), and, the recently published **`dmrcate`** in the DMRcate package (Peters et al. 2015). They are each based on different statistical methods. In our experience, the **`bumphunter`** and **`dmrFind`** functions can be somewhat slow to run unless you have the computer infrastructure to parallelise them, as they use permutations to assign significance. In this workflow, we will perform an analysis using the **`dmrcate`**. As it is based on *limma*, we can directly use the **`design`** and **`contMatrix`** we previously defined.

Firstly, our matrix of M-values is annotated with the relevant information about the probes such as their genomic position, gene annotation, etc. By default, this is done using the **`ilmn12.hg19`** annotation, but this can be substituted for any argument compatible with the interface provided by the *minfi* package. The *limma* pipeline is then used for differential methylation analysis to calculate moderated t-statistics.

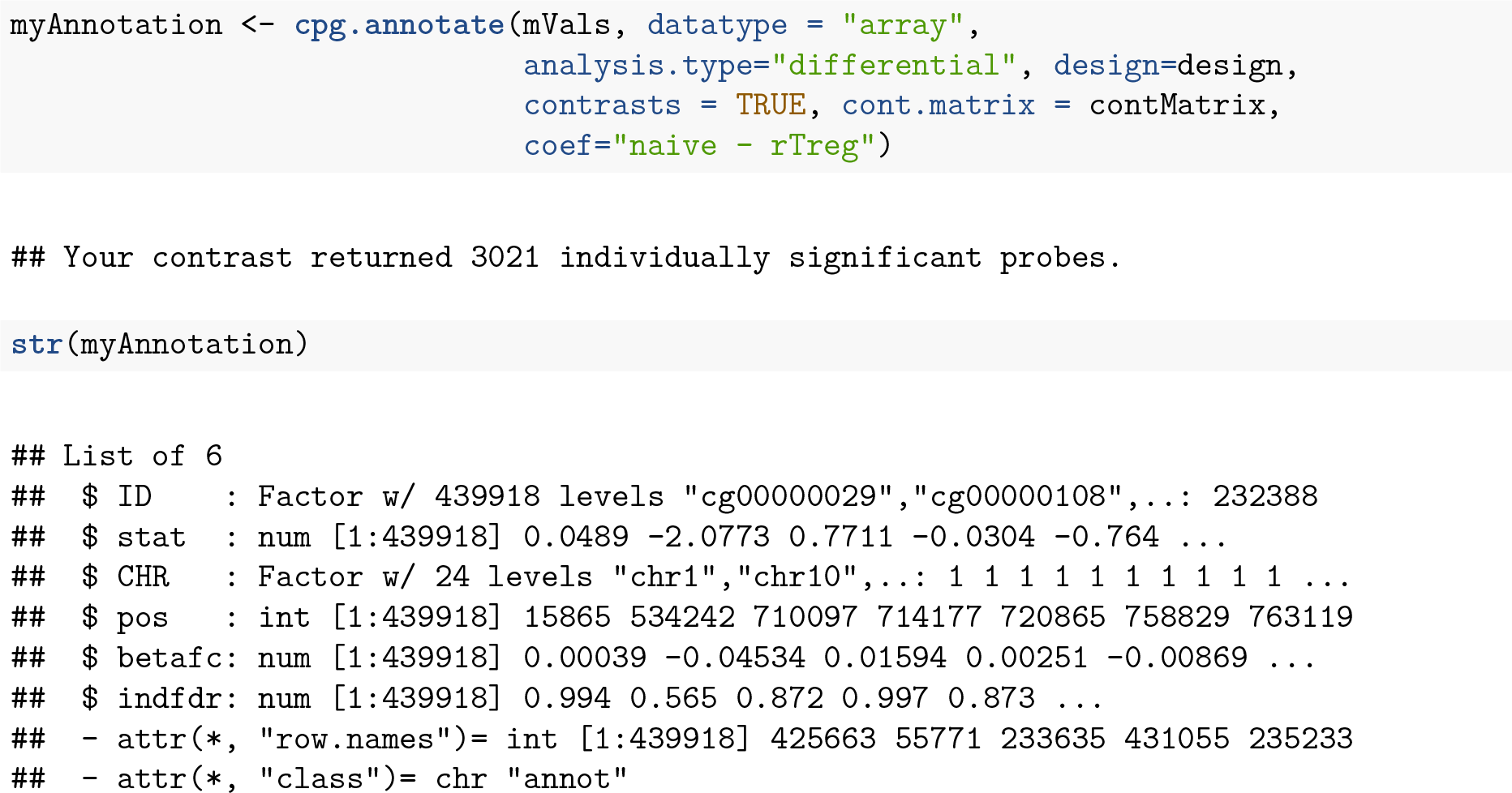

Once we have the relevant statistics for the individual CpGs, we can then use the **`dmrcate`** function to combine them to identify differentially methylated regions. The main output table **`DMRs$results`** contains all of the regions found, along with their genomic annotations and p-values.

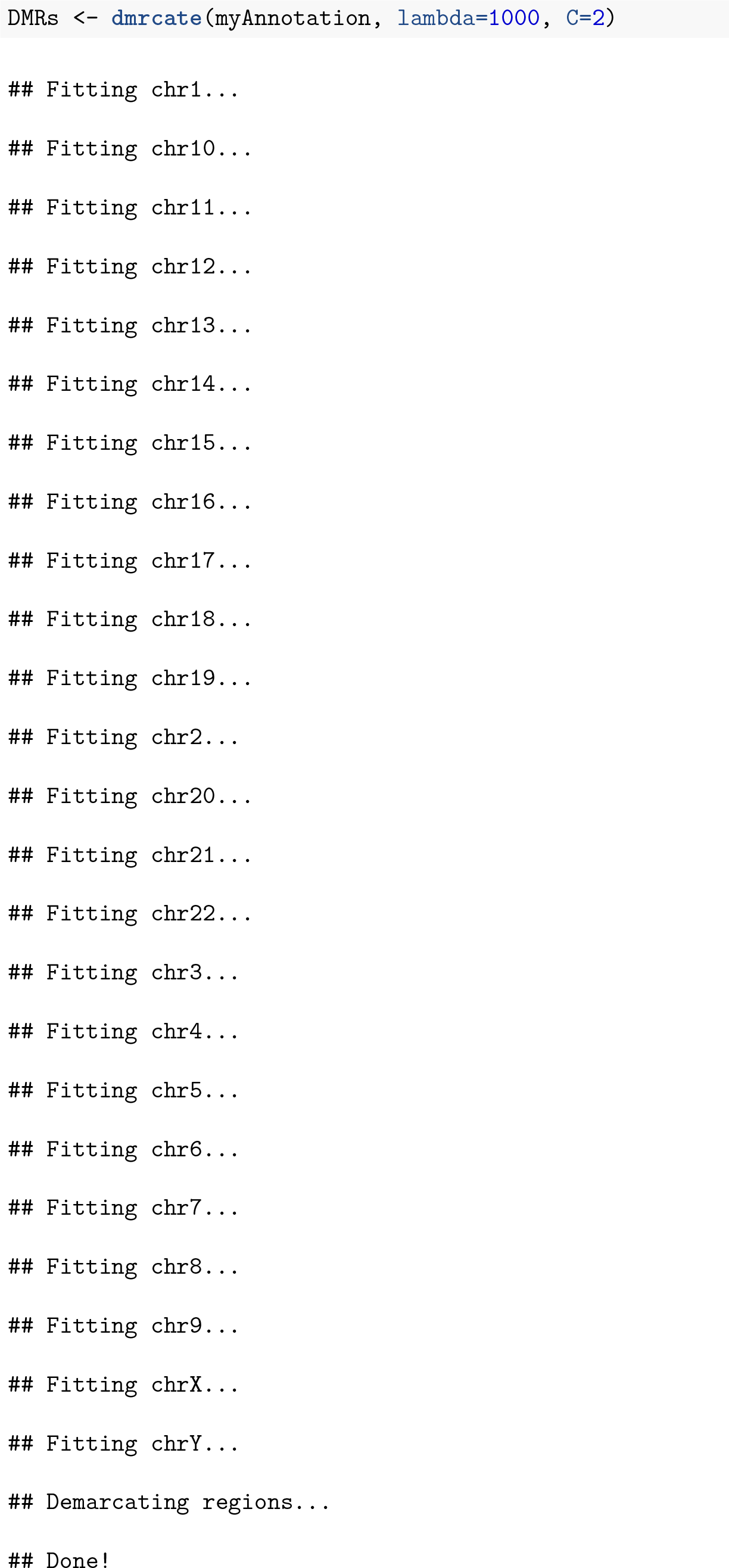

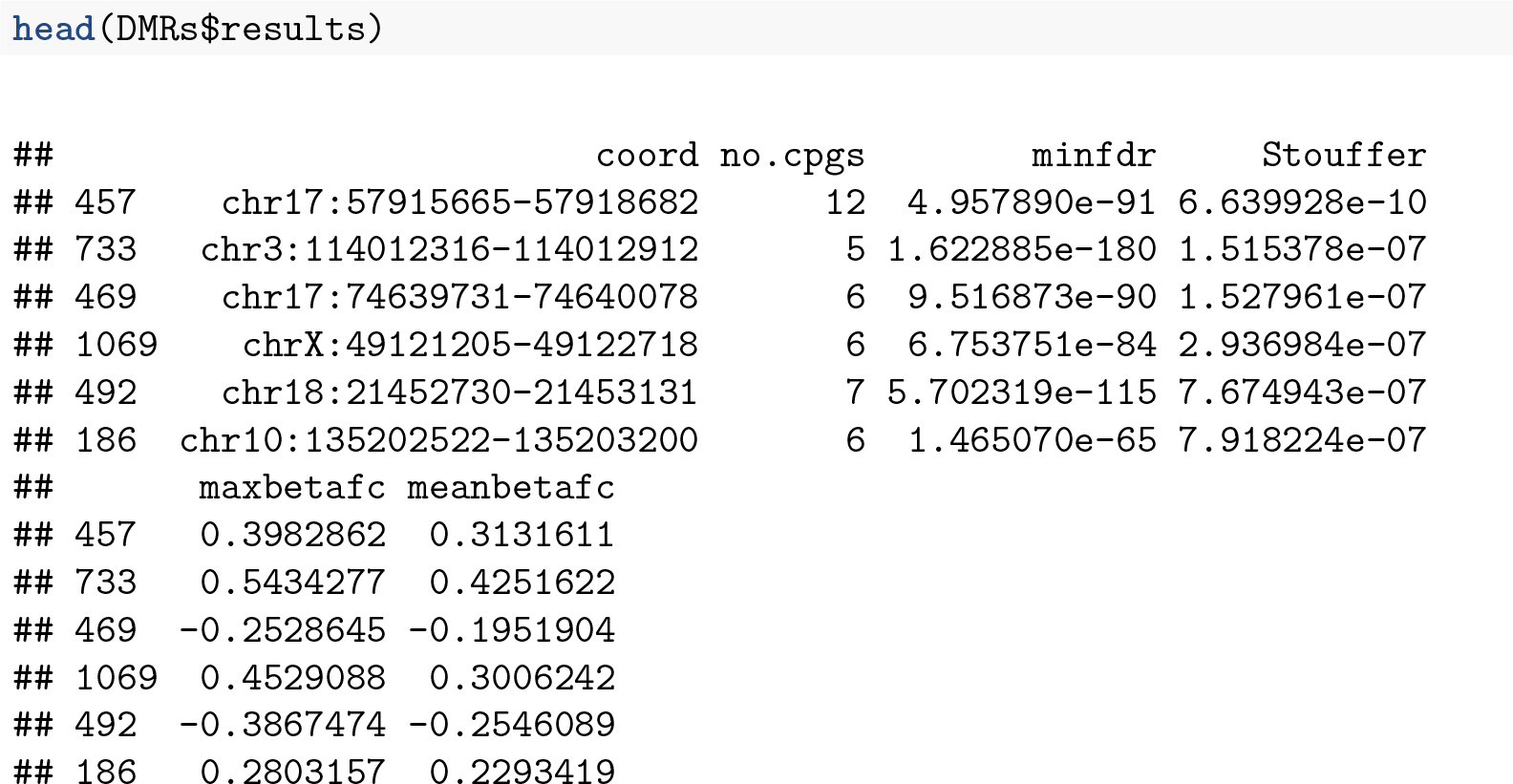

As for the probe-wise analysis, it is advisable to visualise the results to ensure that they make sense. The regions can easily be viewed using the **`DMR.plot`** function provided in the *DMRcate* package.

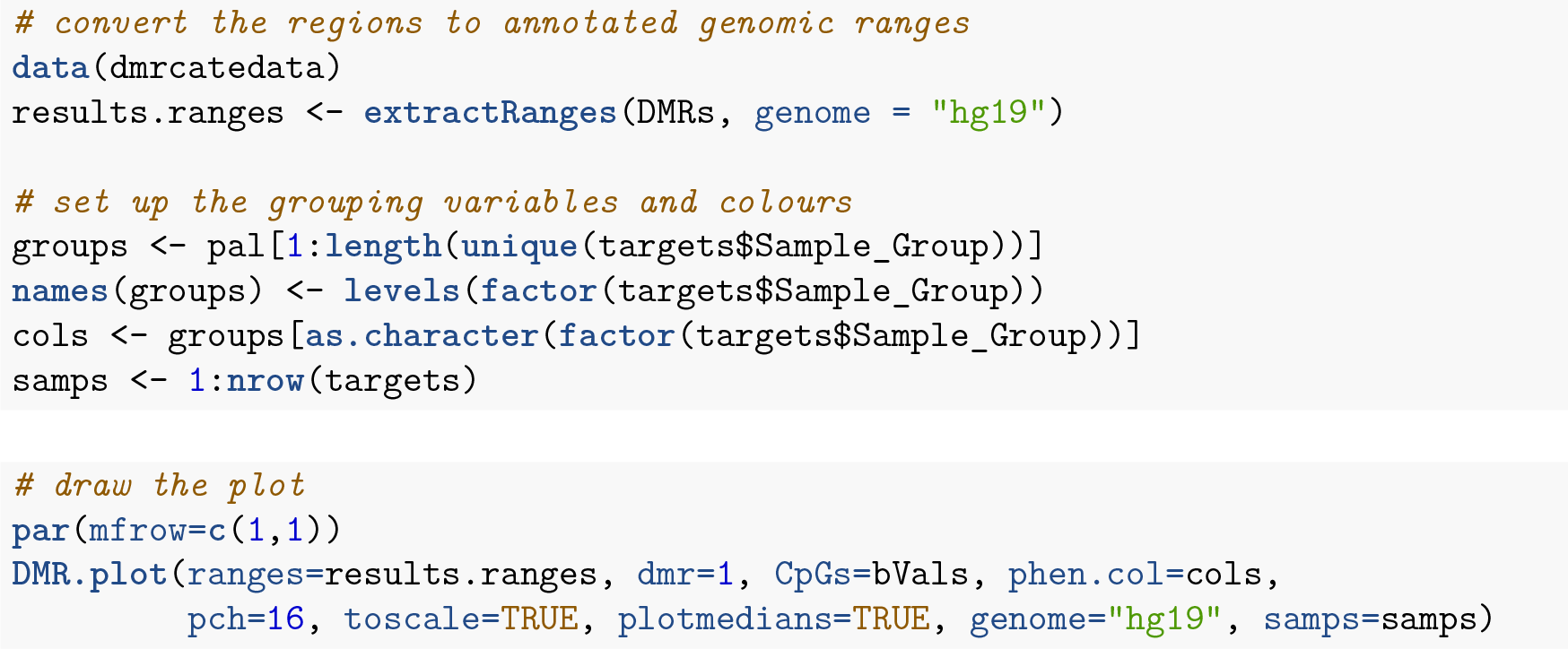

## Customising visualisations of methylation data

The *Gviz* package offers powerful functionality for plotting methylation data in its genomic context. The package vignette is very extensive and covers the various types of plots that can be produced using the *Gviz* framework. We will re-plot the top differentially methylated region from the *DMRcate* regional analysis to demonstrate the type of visualisations that can be created.

We will first set up the genomic region we would like to plot by extracting the genomic coordinates of the top differentially methylated region.

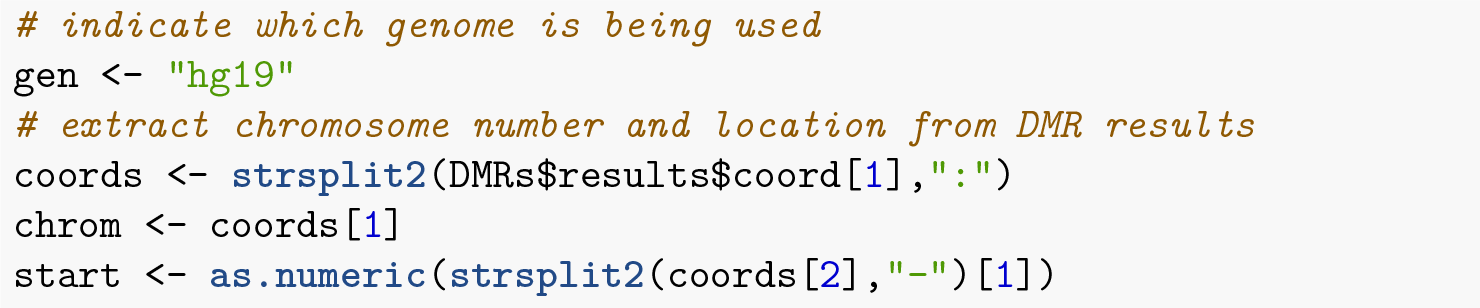

**Figure 10:**
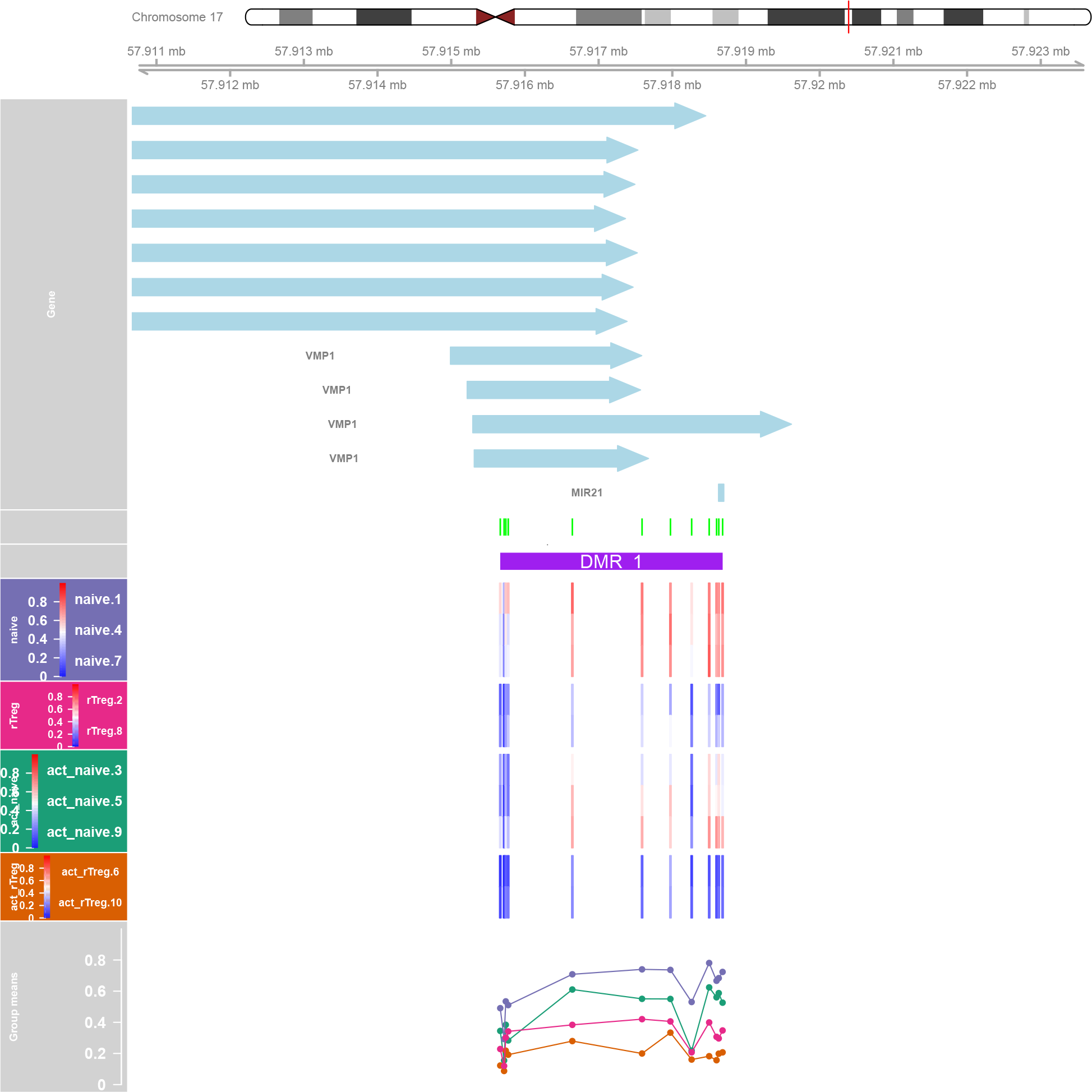
DMRcate provides a function for plotting differentially methylated regions in their genomic context.

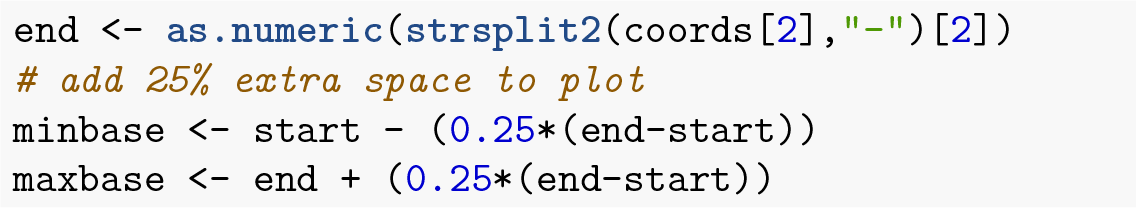

Next, we will add some genomic annotations of interest such as the locations of CpG islands and DNAsel hypersensitive sites; this can be any feature or genomic annotation of interest that you have data available for. The CpG islands data was generated using the method published by H. Wu et al. (2010); the DNAsel hypersensitive site data was obtained from the UCSC Genome Browser.

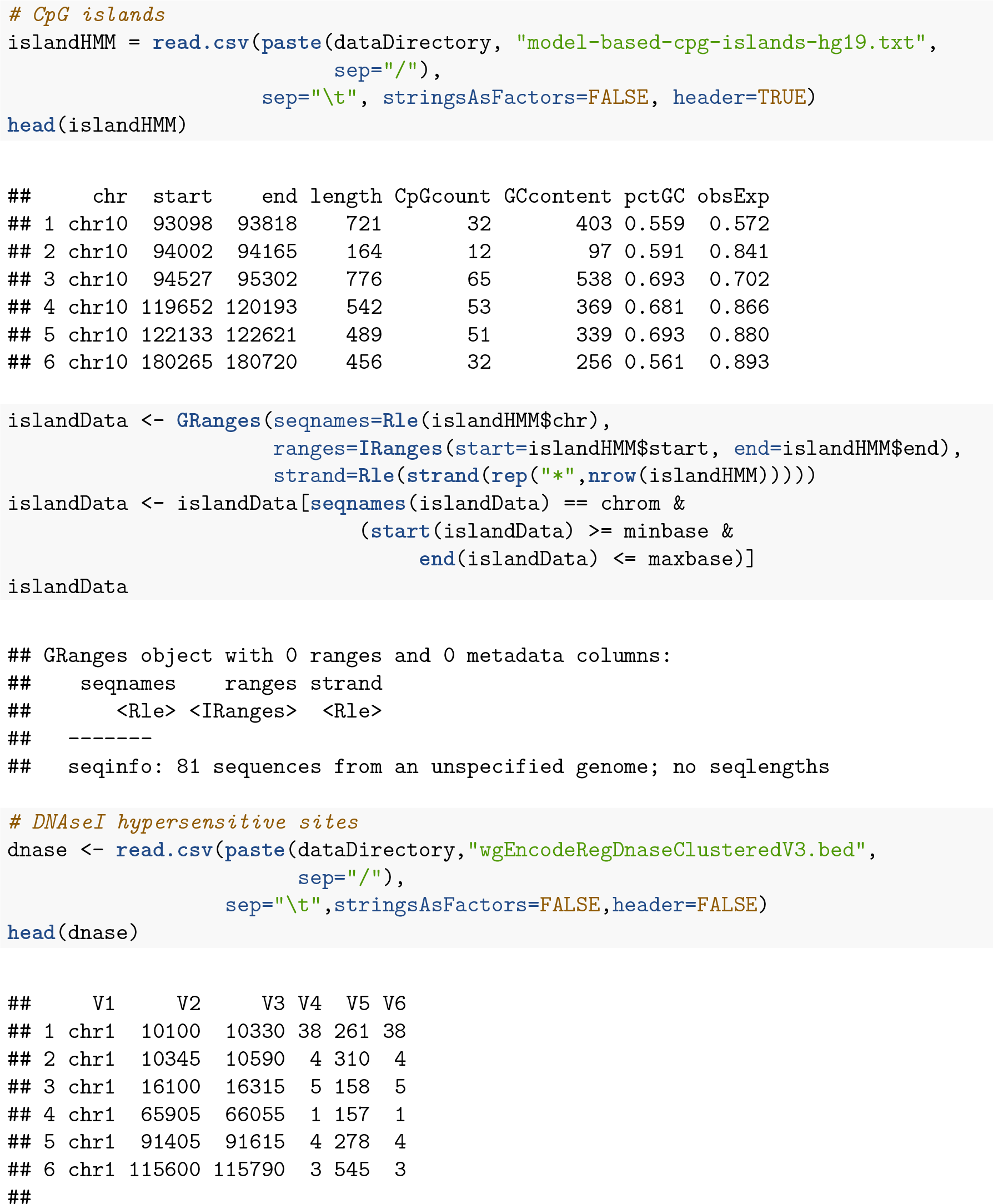

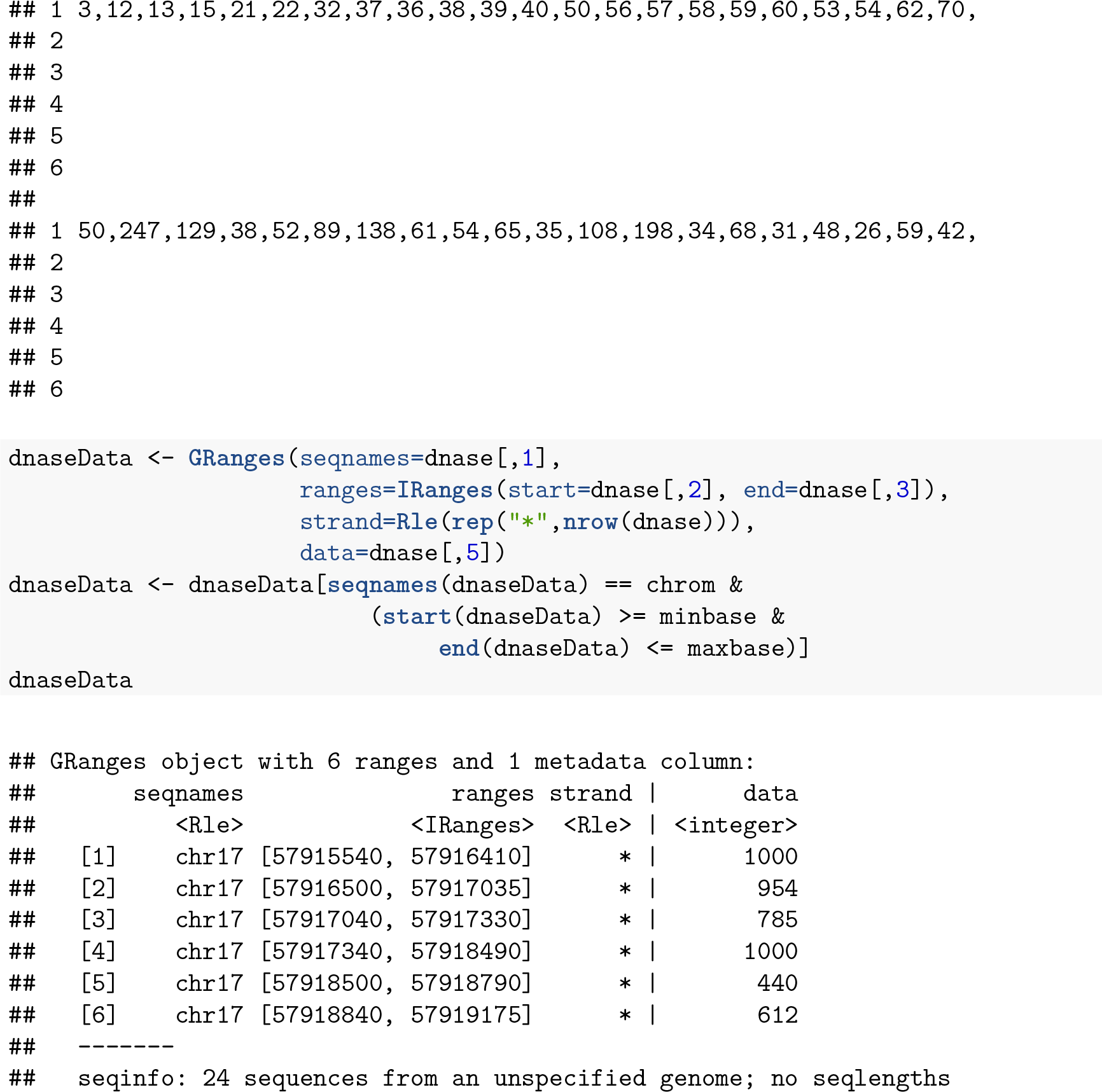

Now, set up the ideogram, genome and RefSeq tracks that will provide context for our methylation data.

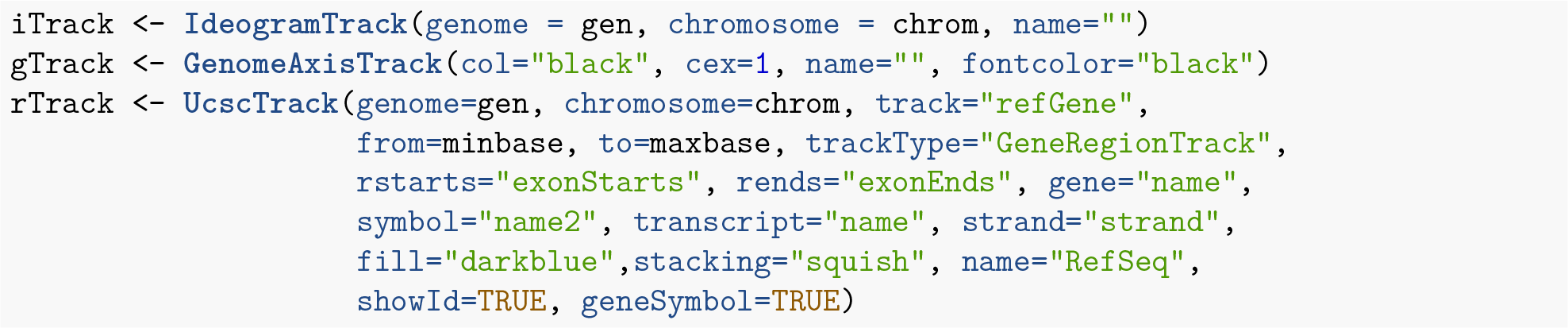

Ensure that the methylation data is ordered by chromosome and base position.

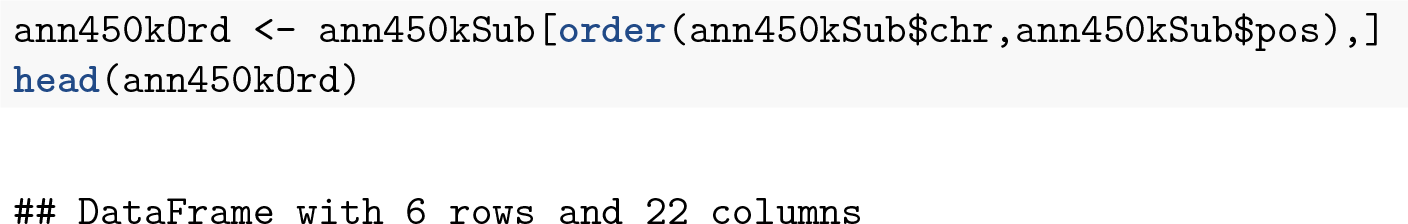

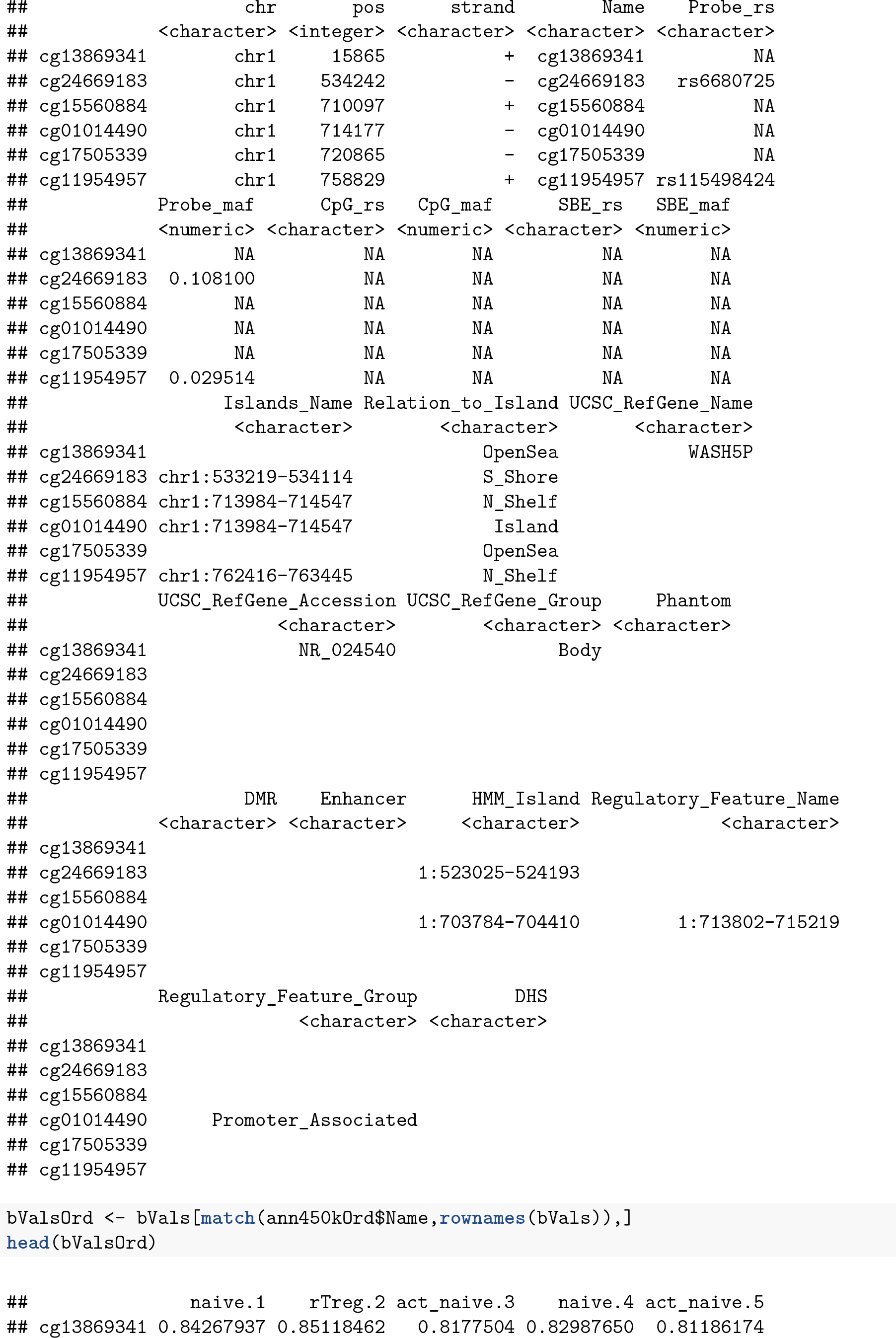

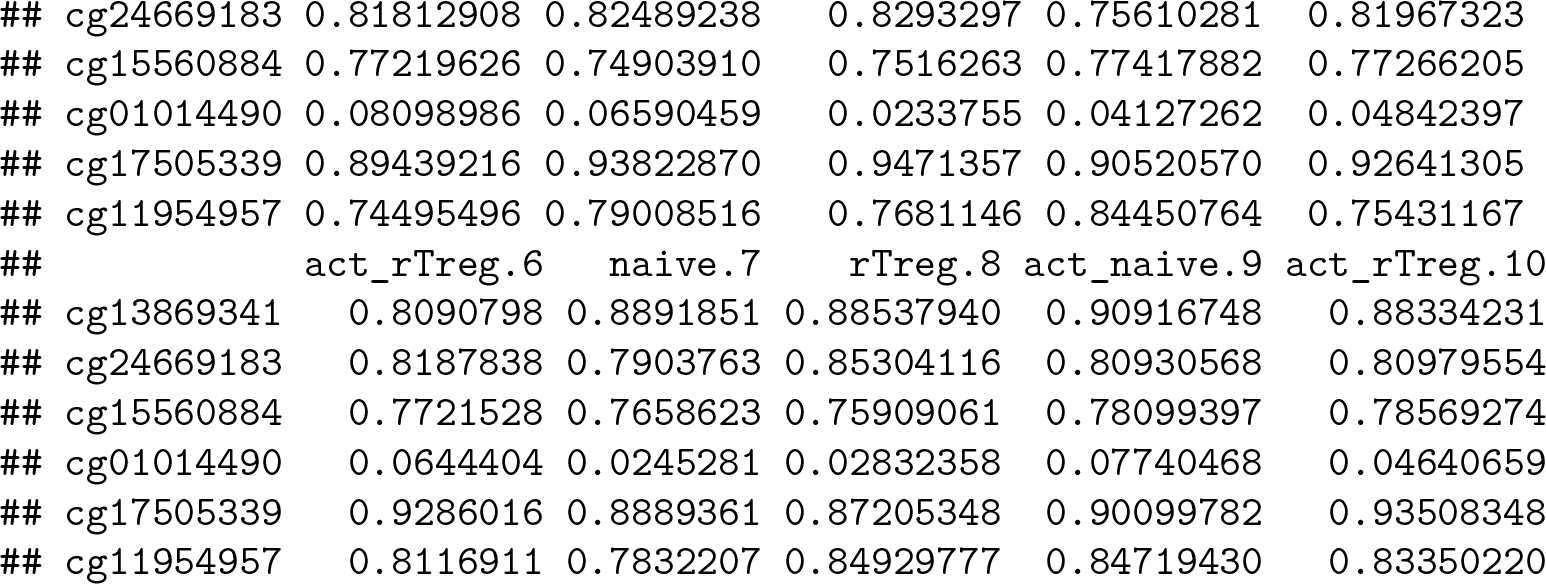

Create the data tracks using the appropriate track type for each data type.

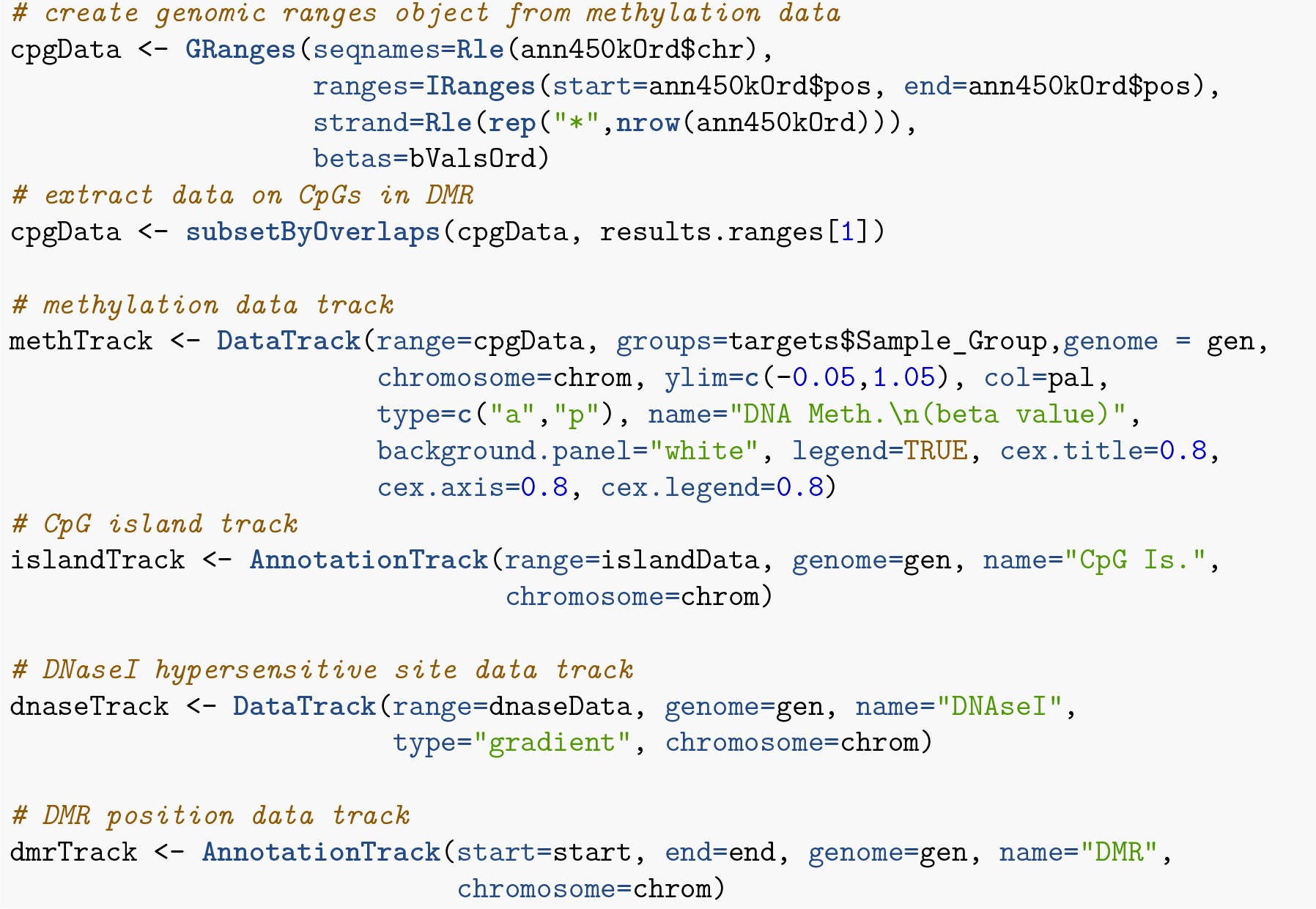

Set up the track list and indicate the relative sizes of the different tracks. Finally, draw the plot using the **`plotTracks`** function.

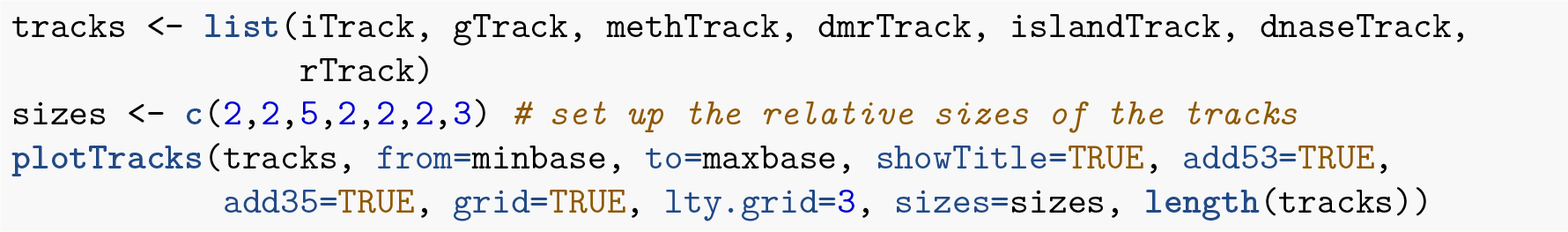

**Figure 11:**
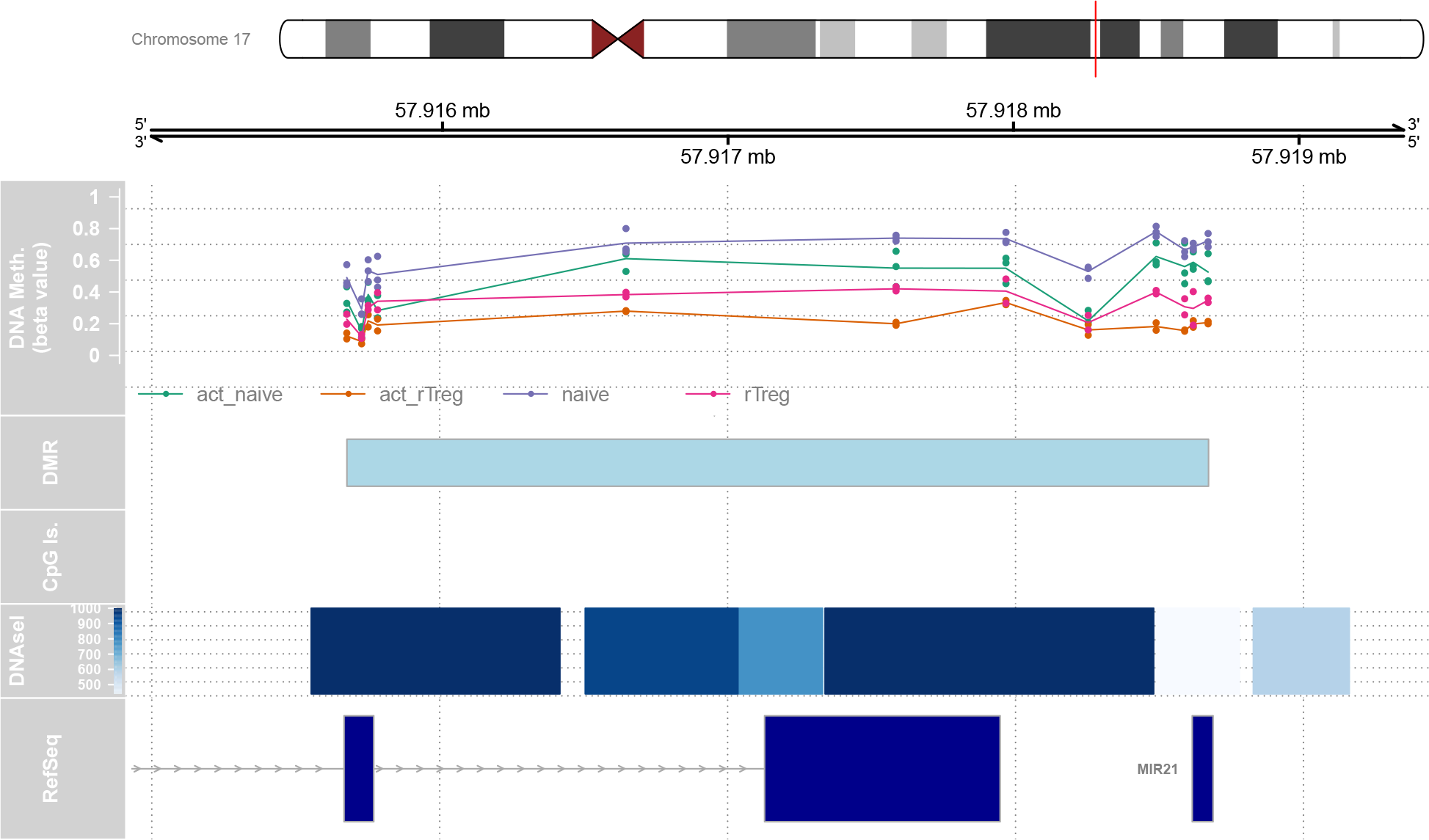
The Gviz package provides extensive functionality for customising plots of genomic regions.

## Additional analyses

### Gene ontology testing

Once you have performed a differential methylation analysis, there may be a very long list of significant CpG sites to interpret. One question a researcher may have is, “which gene pathways are over-represented for differentially methylated CpGs?” In some cases it is relatively straightforward to link the top differentially methylated CpGs to genes that make biological sense in terms of the cell types or samples being studied, but there may be many thousands of CpGs significantly differentially methylated. In order to gain an understanding of the biological processes that the differentially methylated CpGs may be involved in, we can perform gene ontology or KEGG pathway analysis using the **`gometh`** function in the *missMethyl* package (B. Phipson, Maksimovic, and Oshlack 2016).

Let us consider the first comparison, naive vs rTreg, with the results of the analysis in the **`DMPs`** table. The **`gometh`** function takes as input a character vector of the names (e.g. cg20832020) of the significant CpG sites, and optionally, a character vector of all CpGs tested. This is recommended particularly if extensive filtering of the CpGs has been performed prior to analysis. For gene ontology testing (default), the user can specify **`collection=“GO”`**; for KEGG testing **`collection=“KEGG”`**. In the **`DMPs`** table, the **`Name`** column corresponds to the CpG name. We will select all CpG sites that have adjusted p-value of less than 0.05.

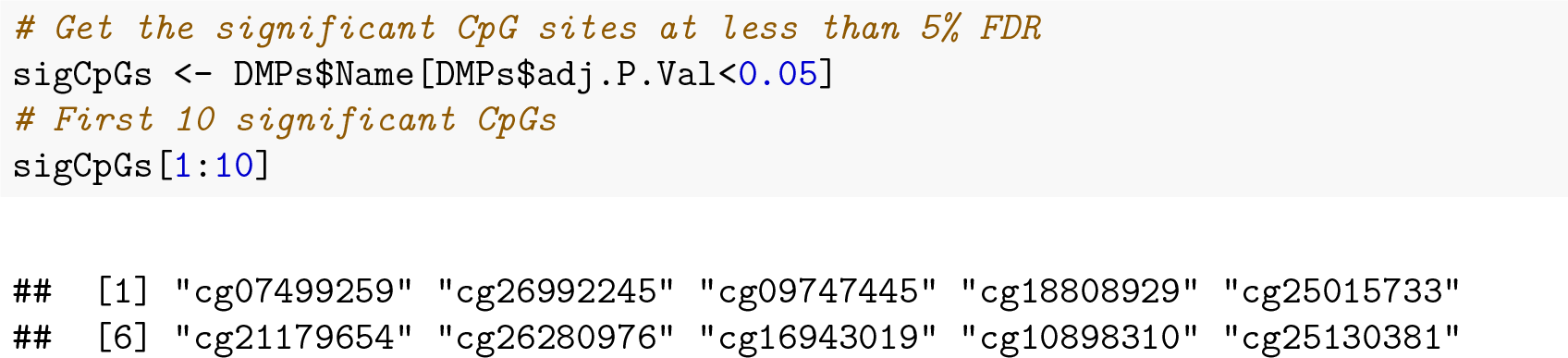

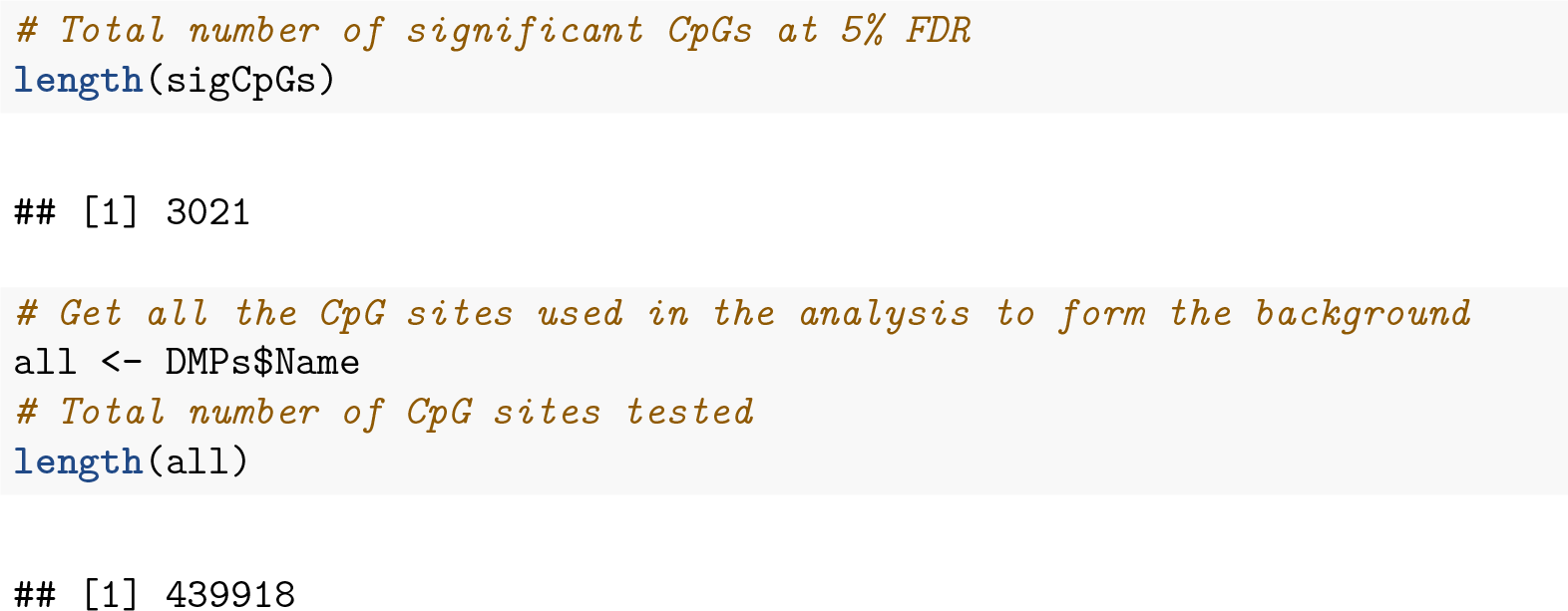

The **`gometh`** function takes into account the varying numbers of CpGs associated with each gene on the Illumina methylation arrays. For the 450k array, the numbers of CpGs mapping to genes can vary from as few as 1 to as many as 1200. The genes that have more CpGs associated with them will have a higher probability of being identified as differentially methylated compared to genes with fewer CpGs. We can look at this bias in the data by specifying **`plot=TRUE`** in the call to **`gometh`**.

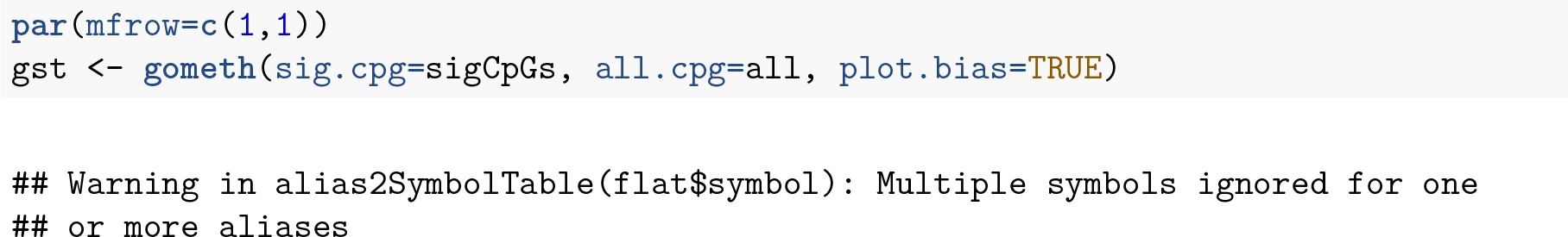

The **`gst`** object is a **`data.frame`** with each row corresponding to the GO category being tested. The top 20 gene ontology categories can be displayed using the topGO function. For KEGG pathway analysis, the **`topKEGG`** function can be called to display the top 20 enriched pathways.

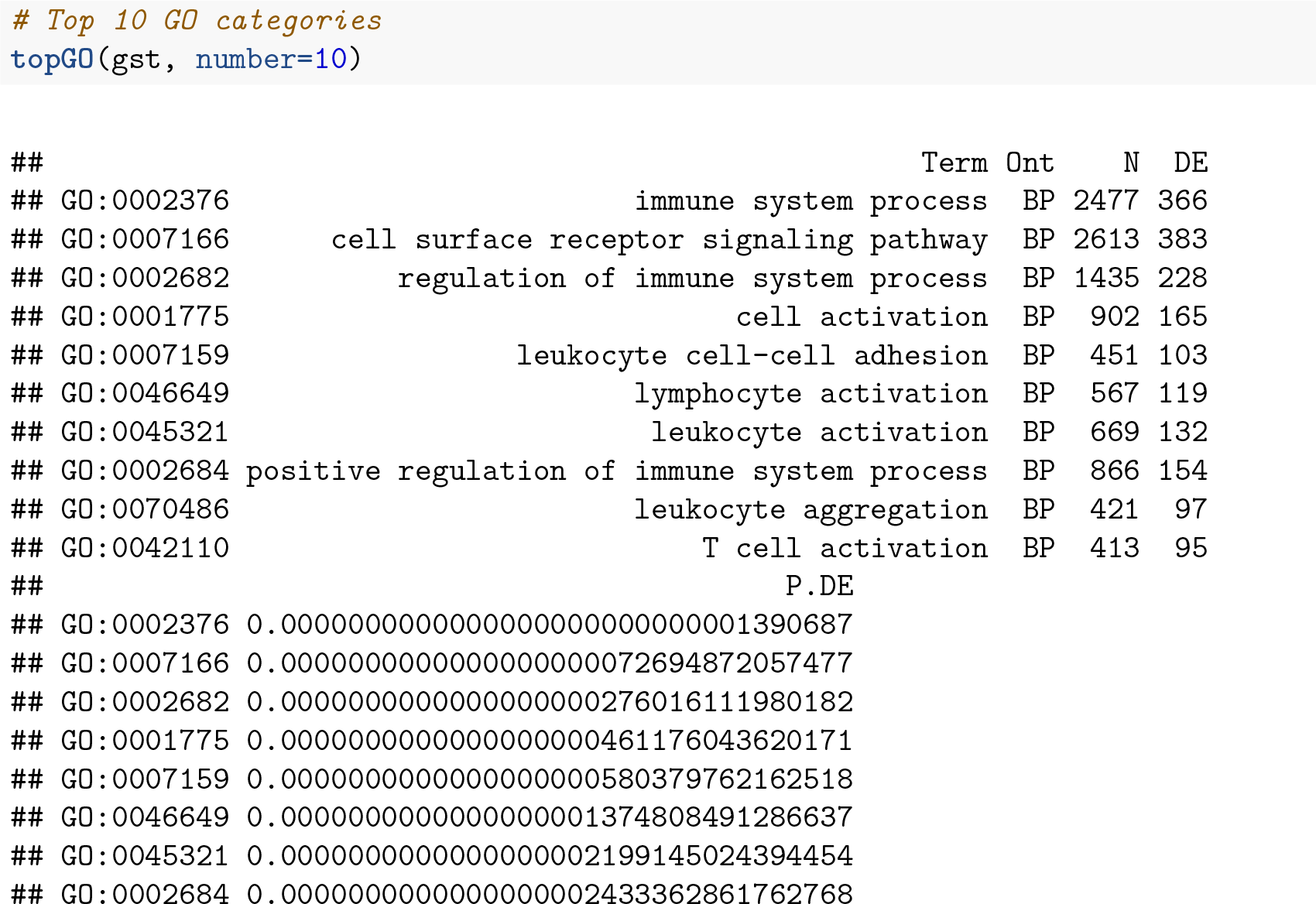

**Figure 12:**
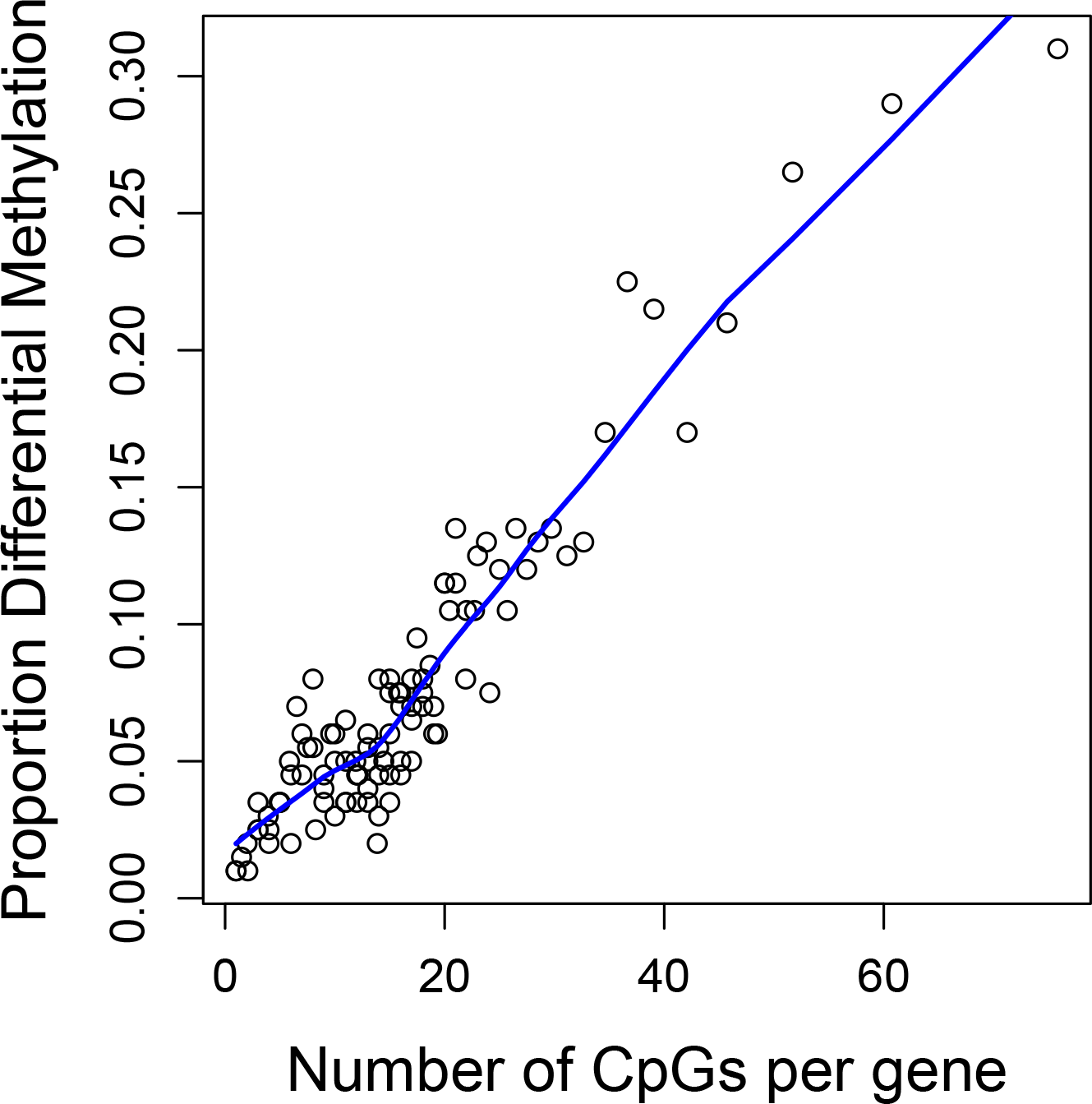
Bias resulting from different numbers of CpG probes in different genes.

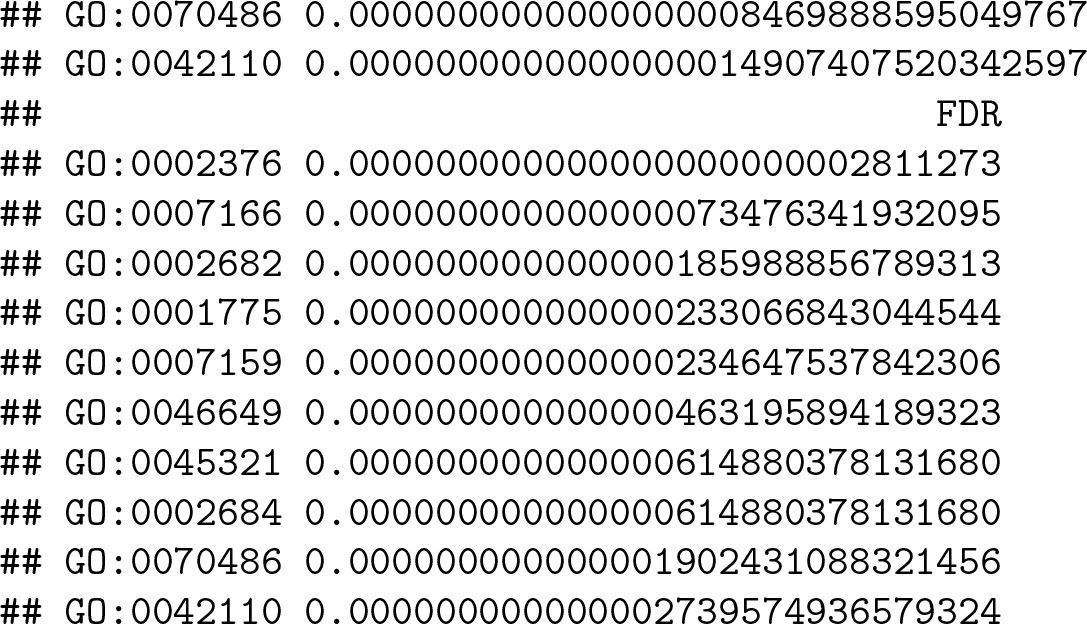

From the output we can see many of the top GO categories correspond to immune system and T cell processes, which is unsurprising as the cell types being studied form part of the immune system.

For a more generalised version of gene set testing for methylation data where the user can specify the gene set to be tested, the **`gsameth`** function can be used. To display the top 20 pathways, **`topGSA`** can be called. **`gsameth`** accepts a single gene set, or a list of gene sets. The gene identifiers in the gene set must be Entrez Gene IDs. To demonstrate **`gsameth`**, we are using the curated genesets (C2) from the Broad Institute Molecular signatures database. These can be downloaded as an **`RData`** object from the WEHI Bioinformatics website.

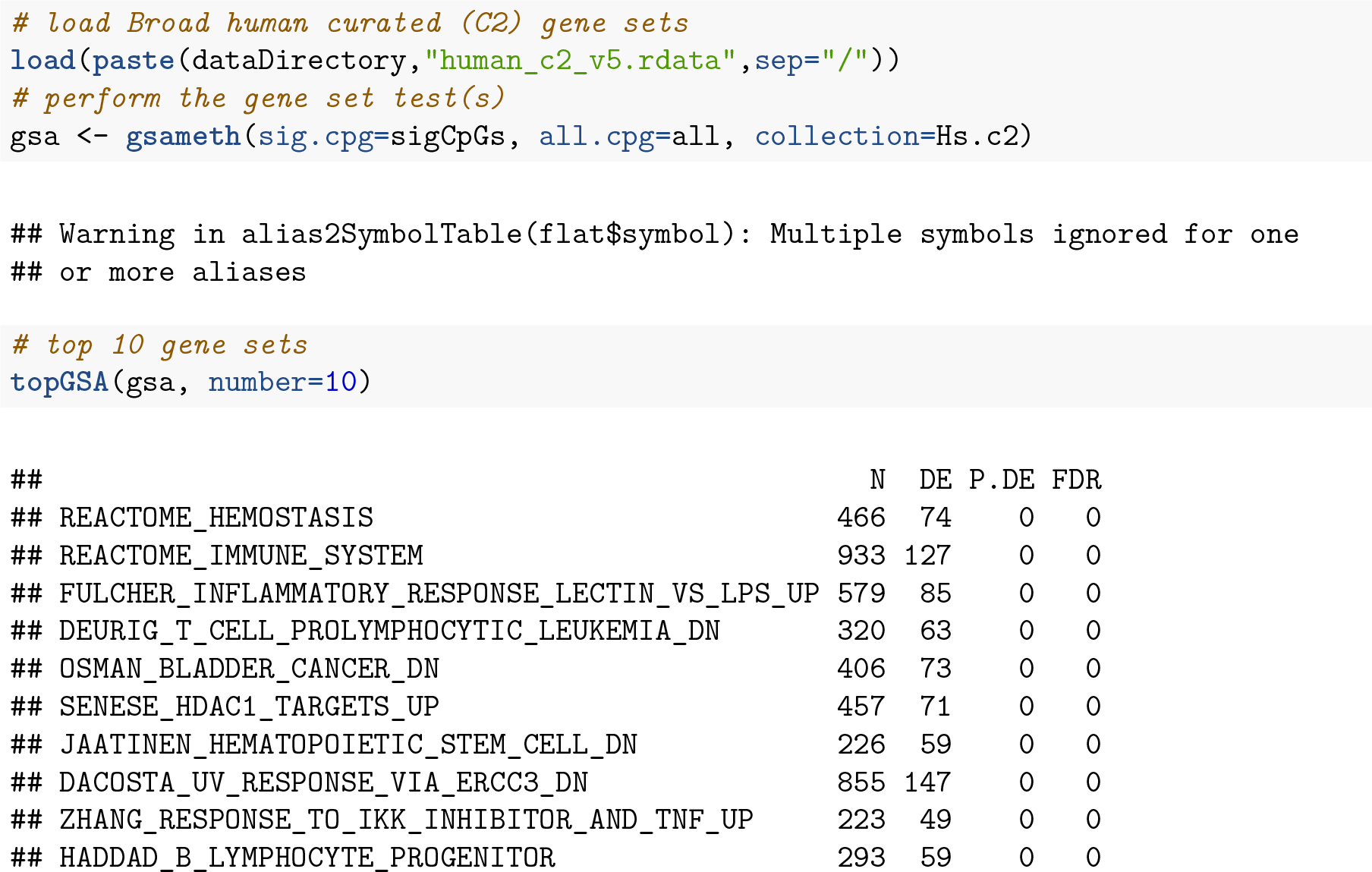

### Differential variability

Rather than testing for differences in mean methylation, we may be interested in testing for differences between group variances. For example, it has been hypothesised that highly variable CpGs in cancer are important for tumour progression. Hence we may be interested in CpG sites that are consistently methylated in one group, but variably methylated in another group.

Sample size is an important consideration when testing for differentially variable CpG sites. In order to get an accurate estimate of the group variances, larger sample sizes are required than for estimating group means. A good rule of thumb is to have at least ten samples in each group (B. Phipson and Oshlack 2014). To demonstrate testing for differentially variable CpG sites, we will use a publicly available dataset on ageing, where whole blood samples were collected from 18 centenarians and 18 newborns and profiled for methylation on the 450k array (Heyn et al. 2012). We will first need to load, normalise and filter the data as previously described.

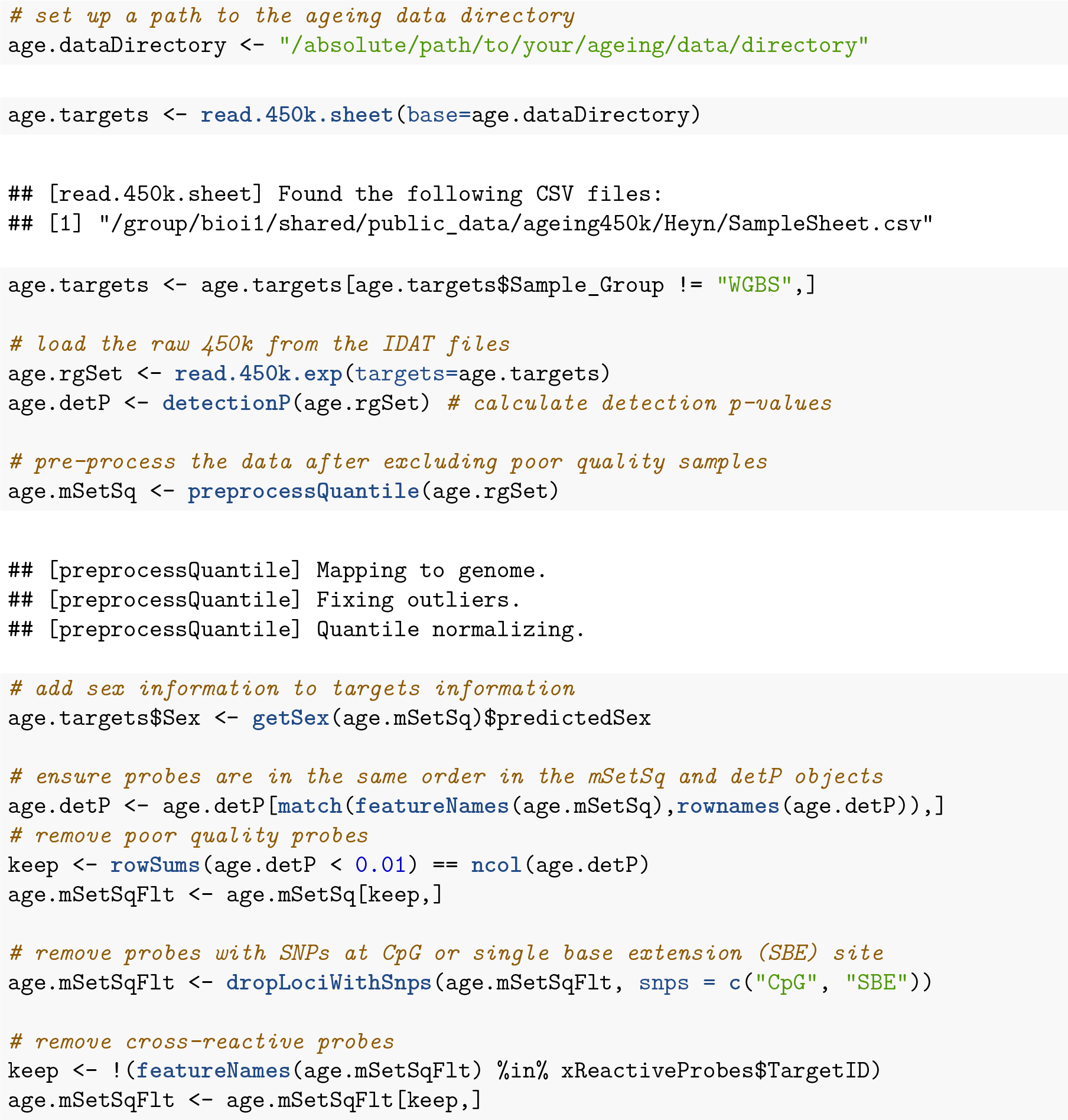

As this dataset contains samples from both males and females, we can use it to demonstrate the effect of removing sex chromosome probes on the data. The MDS plots below show the relationship between the samples in the ageing dataset before and after sex chromosome probe removal. It is apparent that before the removal of sex chromosome probes, the sample cluster based on sex in the second principal component. When the sex chromosome probes are removed, age is the largest source of variation present and the male and female samples no longer form separate clusters.

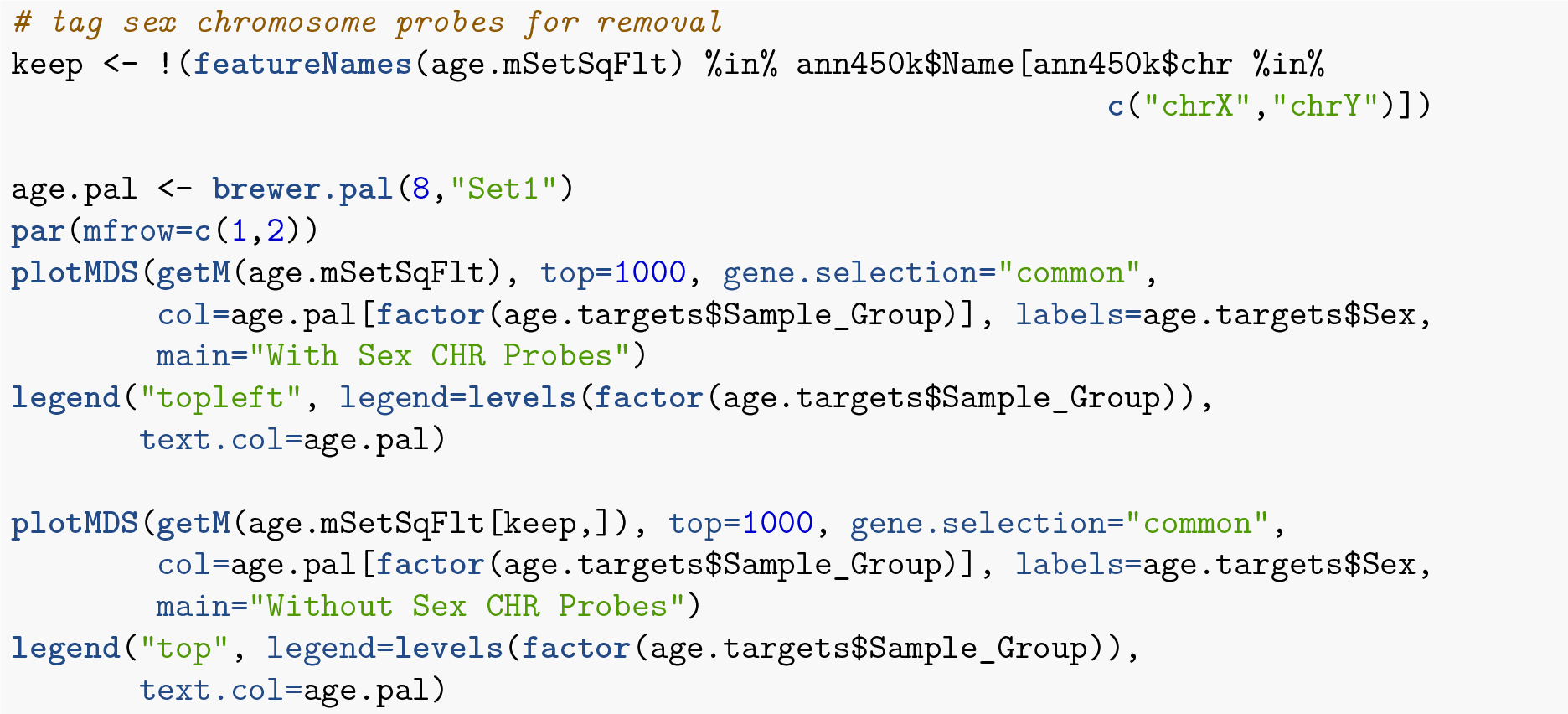

**Figure 13:**
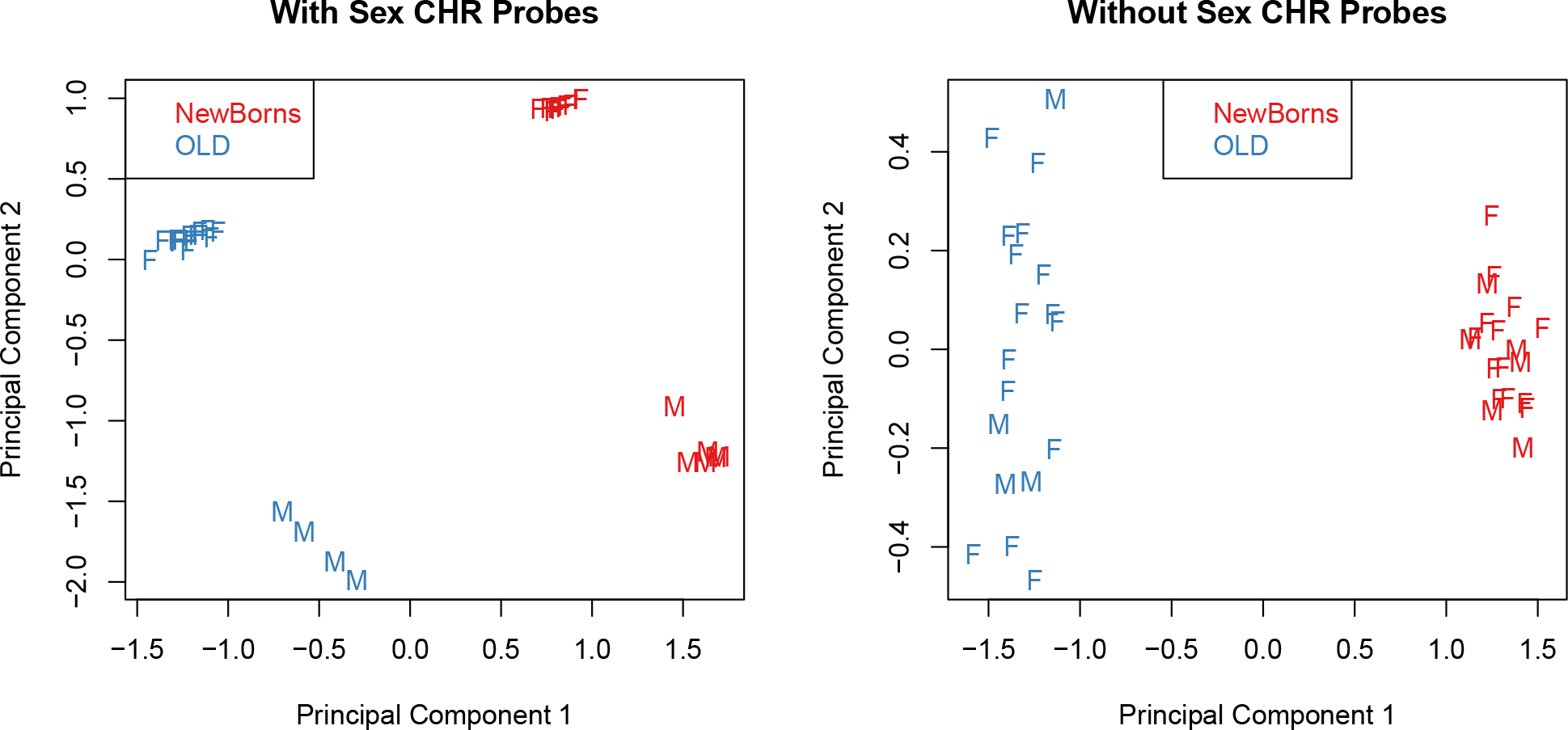
When samples from both males and females are included in a study, sex is usually the largest source of variation in methylation data.

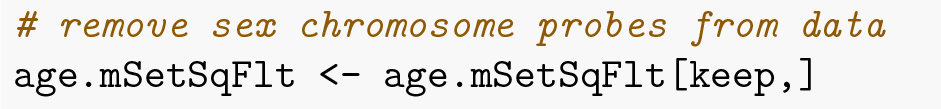

We can test for differentially variable CpGs using the **`varFit`** function in the *missMethyl* package. The syntax for specifying which groups we are interested in testing is slightly different to the standard way a model is specified in **`limma`**, particularly for designs where an intercept is fitted (see *missMethyl* vignette for further details). For the ageing data, the design matrix includes an intercept term, and a term for age. The **`coef`** argument in the **`varFit`** function indicates which columns of the design matrix correspond to the intercept and grouping factor. Thus, for the ageing dataset we set **`coef=c(1,2)`**. Note that design matrices without intercept terms are permitted, with specific contrasts tested using the **`contrasts.varFit`** function.

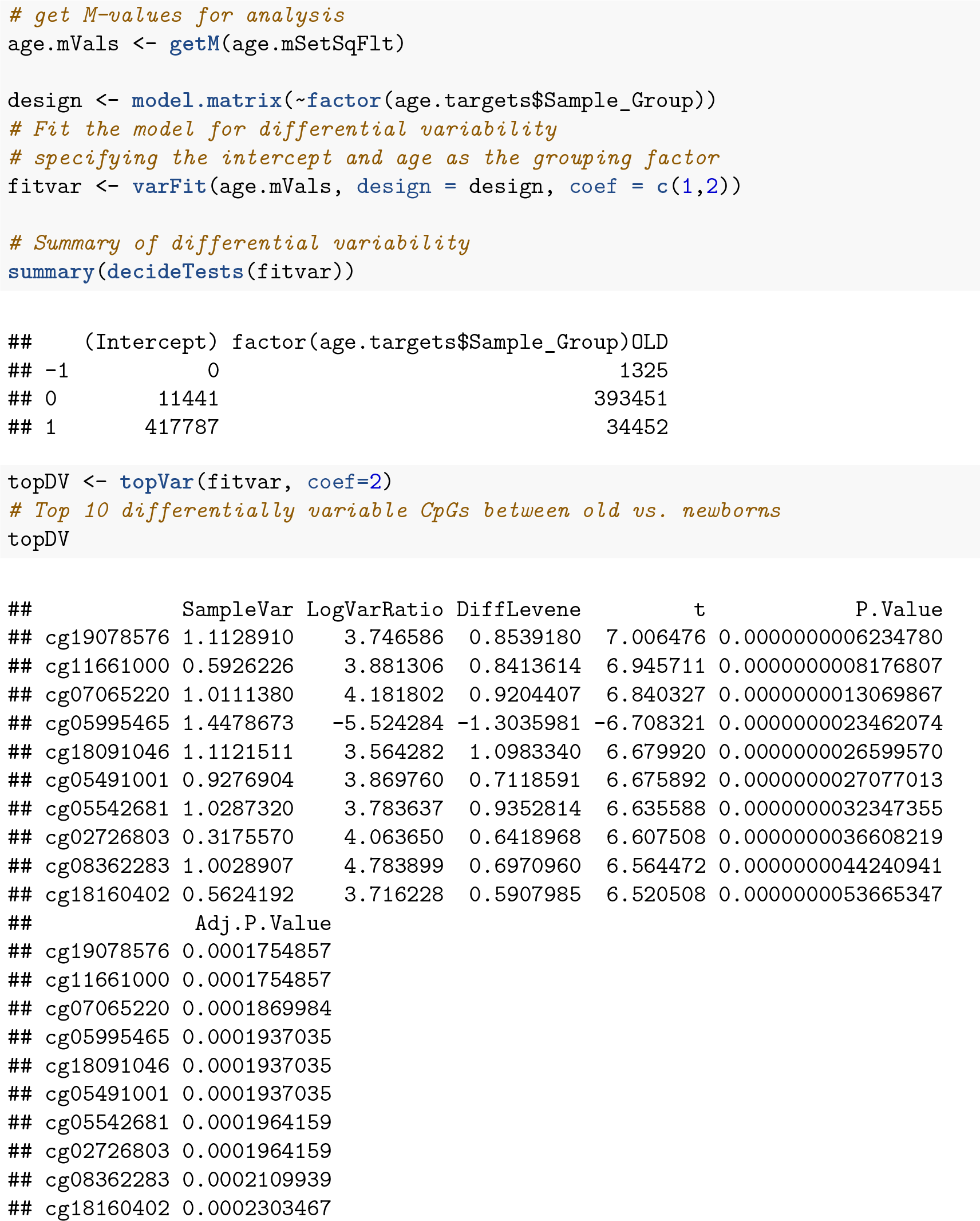

Similarly to the differential methylation analysis, is it useful to plot sample-wise beta values for the differentially variable CpGs to ensure the significant results are not driven by artifacts or outliers.

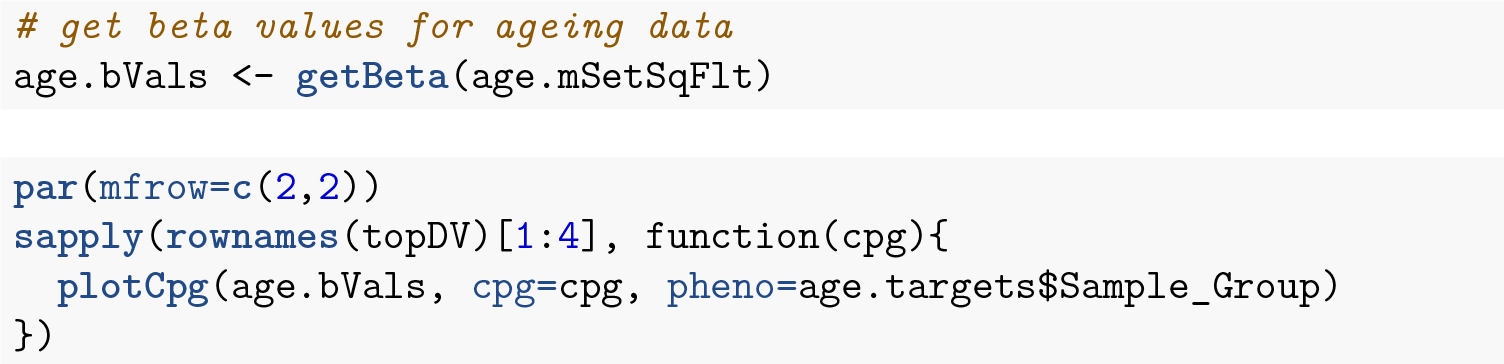

**Figure 14:**
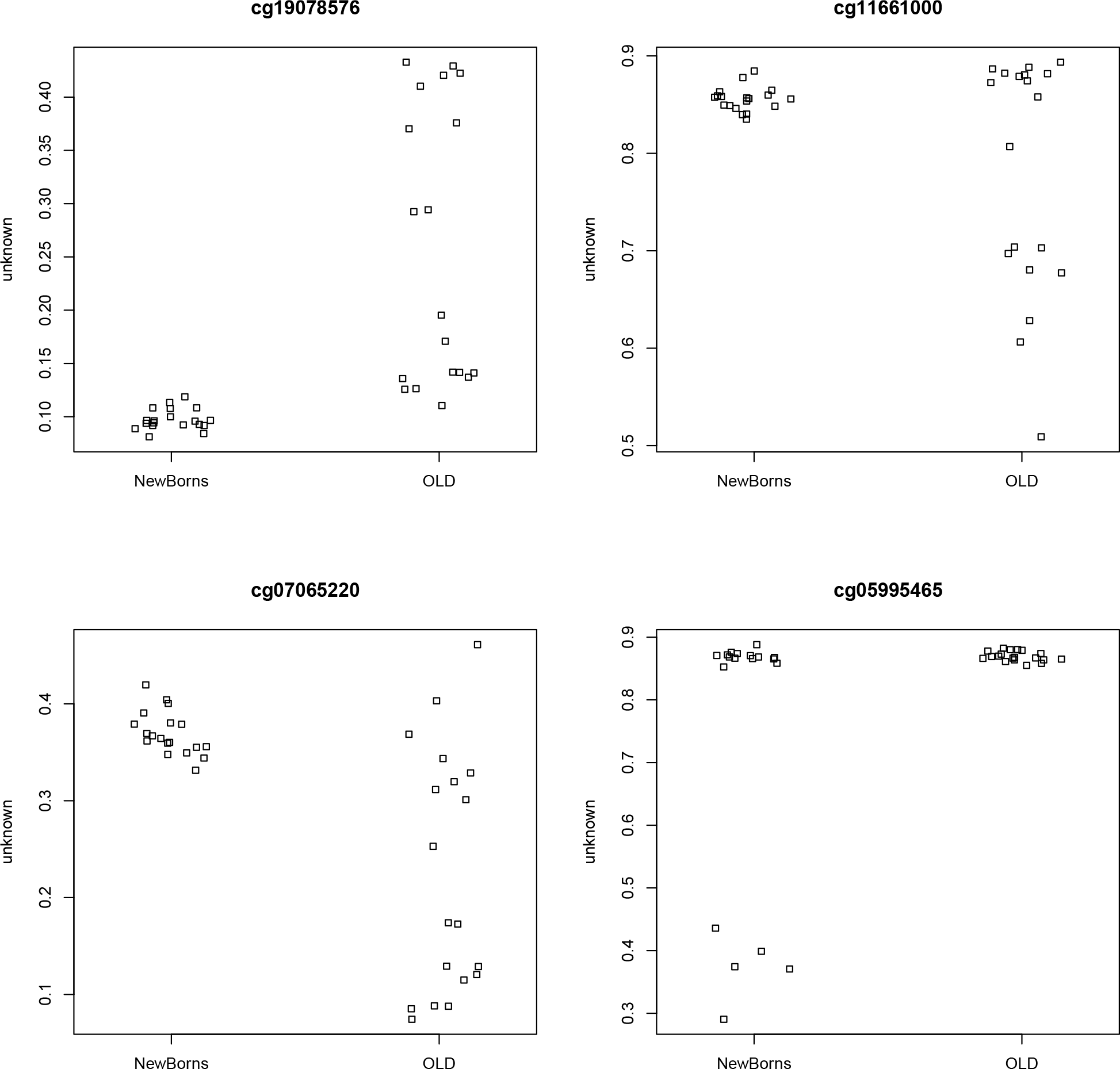
As for DMPs, it is useful to plot the top few differentially variable CpGs to check that the results make sense.

An example of testing for differential variability when the design matrix does not have an intercept term is detailed in the *missMethyl* vignette.

### Cell type composition

As methylation is cell type specific and methylation arrays provide CpG methylation values for a population of cells, biological findings from samples that are comprised of a mixture of cell types, such as blood, can be confounded with cell type composition (Jaffe and Irizarry 2014). The *minfi* function **`estimateCellCounts`** facilitates the estimation of the level of confounding between phenotype and cell type composition in a set of samples. The function uses a modified version of the method published by Houseman et al. (2012) and the package **`FlowSorted.Blood.450k`**, which contains 450k methylation data from sorted blood cells, to estimate the cell type composition of blood samples.

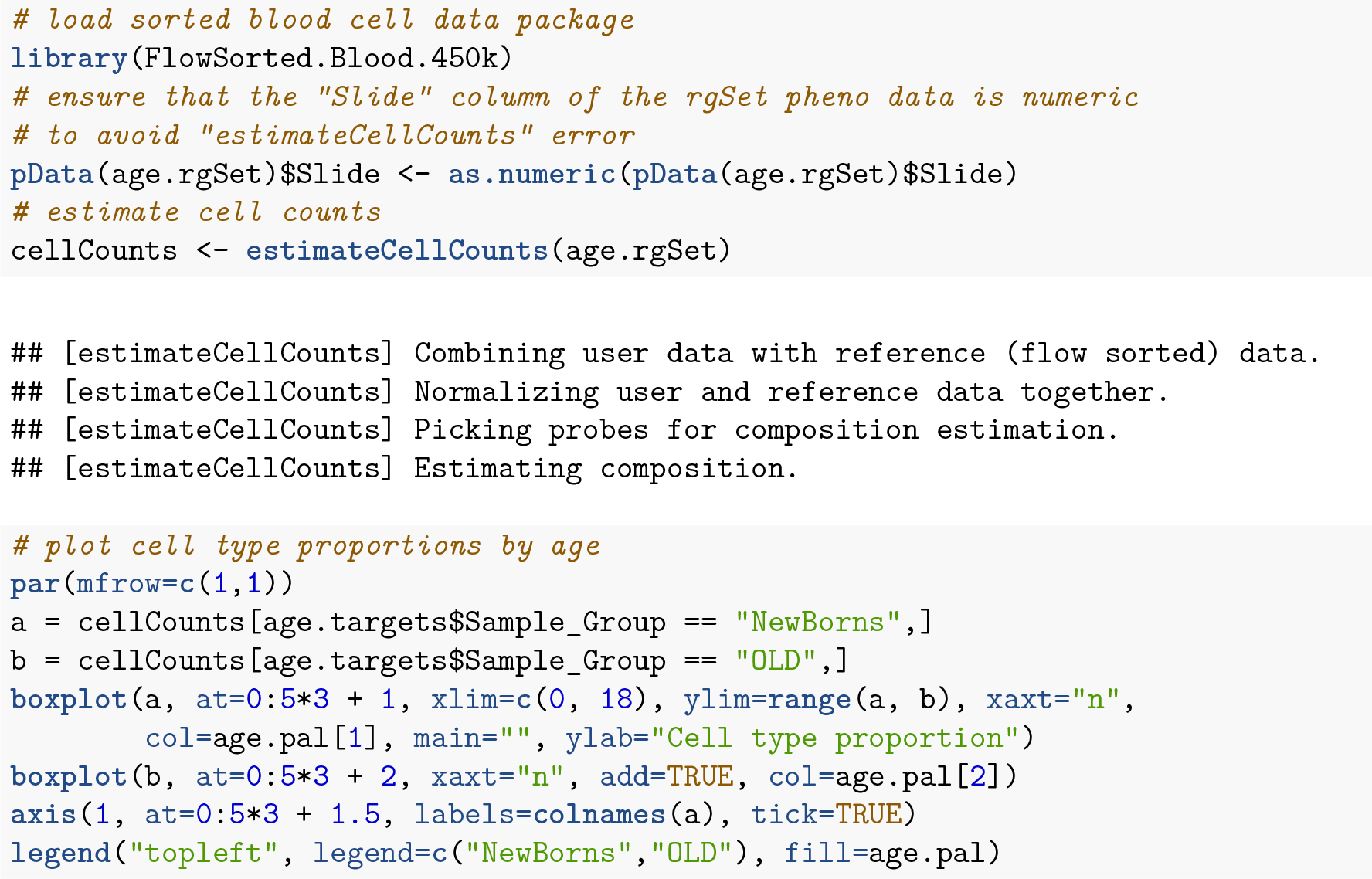

As reported by Jaffe and Irizarry (2014), the preceding plot demonstrates that differences in blood cell type proportions are strongly confounded with age in this dataset. Performing cell composition estimation can alert you to potential issues with confounding when analysing a mixed cell type dataset. Based on the results, some type of adjustment for cell type composition may be considered, although a naive cell type adjustment is not recommended. Jaffe and Irizarry (2014) outline several strategies for dealing with cell type composition issues.

## Discussion

Here we present a commonly used workflow for methylation array analysis based on a series of Bioconductor packages. While we have not included all the possible functions or analysis options that are available for detecting differential methylation, we have demonstrated a common and well used workflow that we regularly use in our own analysis. Specifically, we have not demonstrated more complex types of analyses such as removing unwanted variation in a differential methylation study (Maksimovic et al. 2015; Leek et al. 2012; Teschendorff, Zhuang, and Widschwendter 2011), block finding (K. D. Hansen et al. 2011; Aryee et al. 2014) or A/B compartment prediction (J.-P. Fortin and Hansen 2015). Our differential methylation workflow presented here demonstrates how to read in data, perform quality control and filtering, normalisation and differential methylation testing. In addition we demonstrate analysis for differential variability, gene set testing and estimating cell type composition. One important aspect of exploring results of an analysis is visualisation and we also provide an example of generating region-level views of the data.

**Figure 15:**
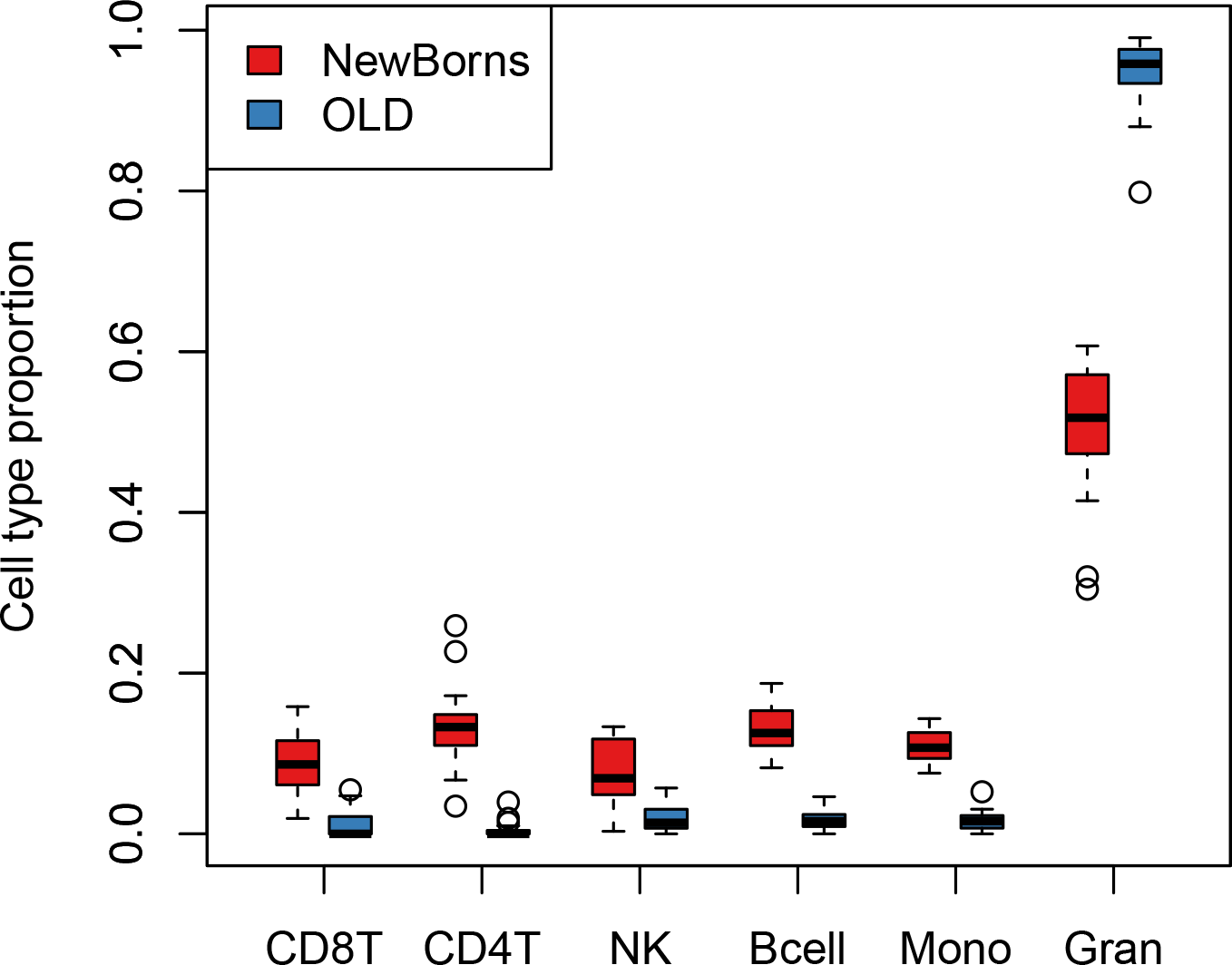
If samples come from a population of mixed cells e.g. blood, it is advisable to check for potential confounding between differences in cell type proportions and the factor of interest.

## Software versions

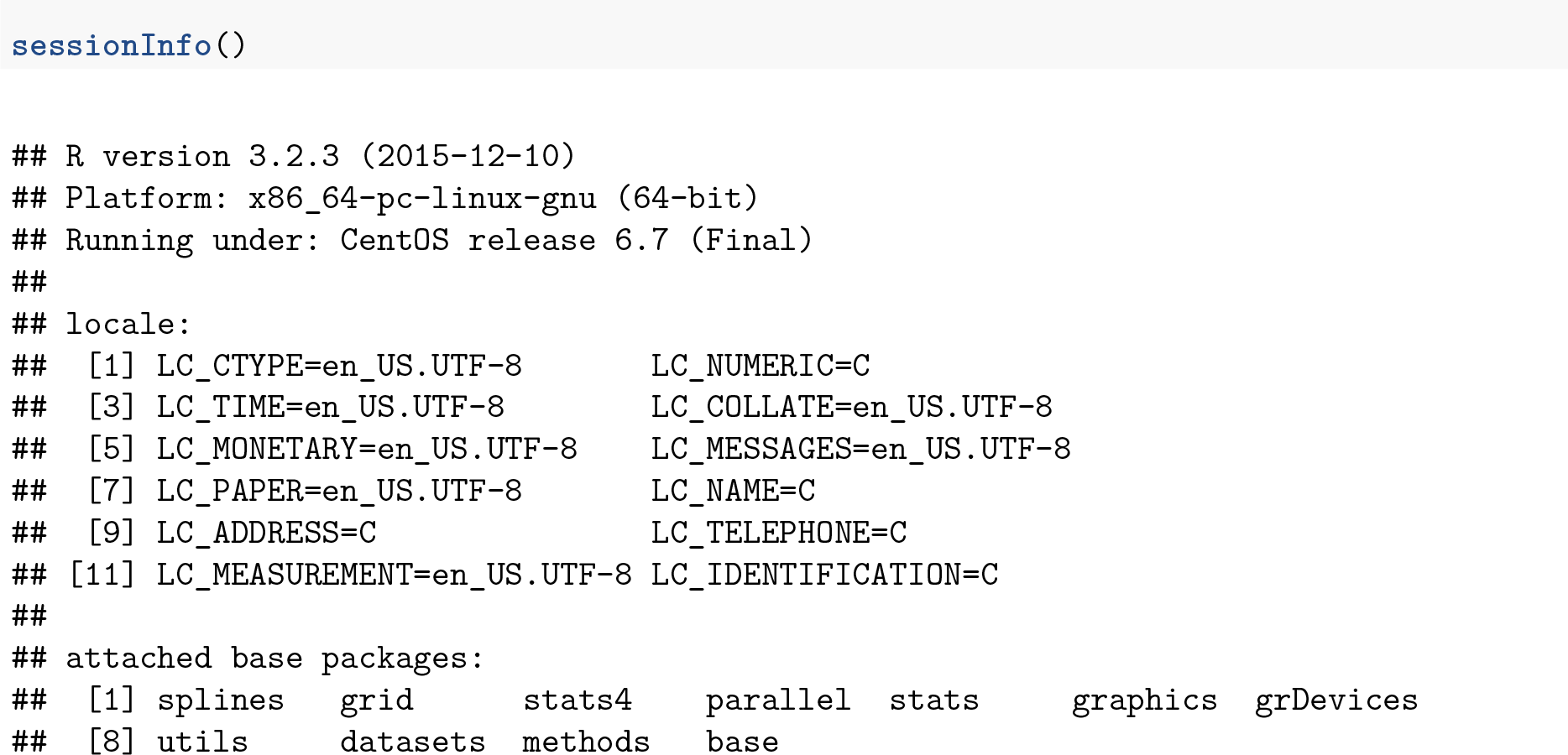

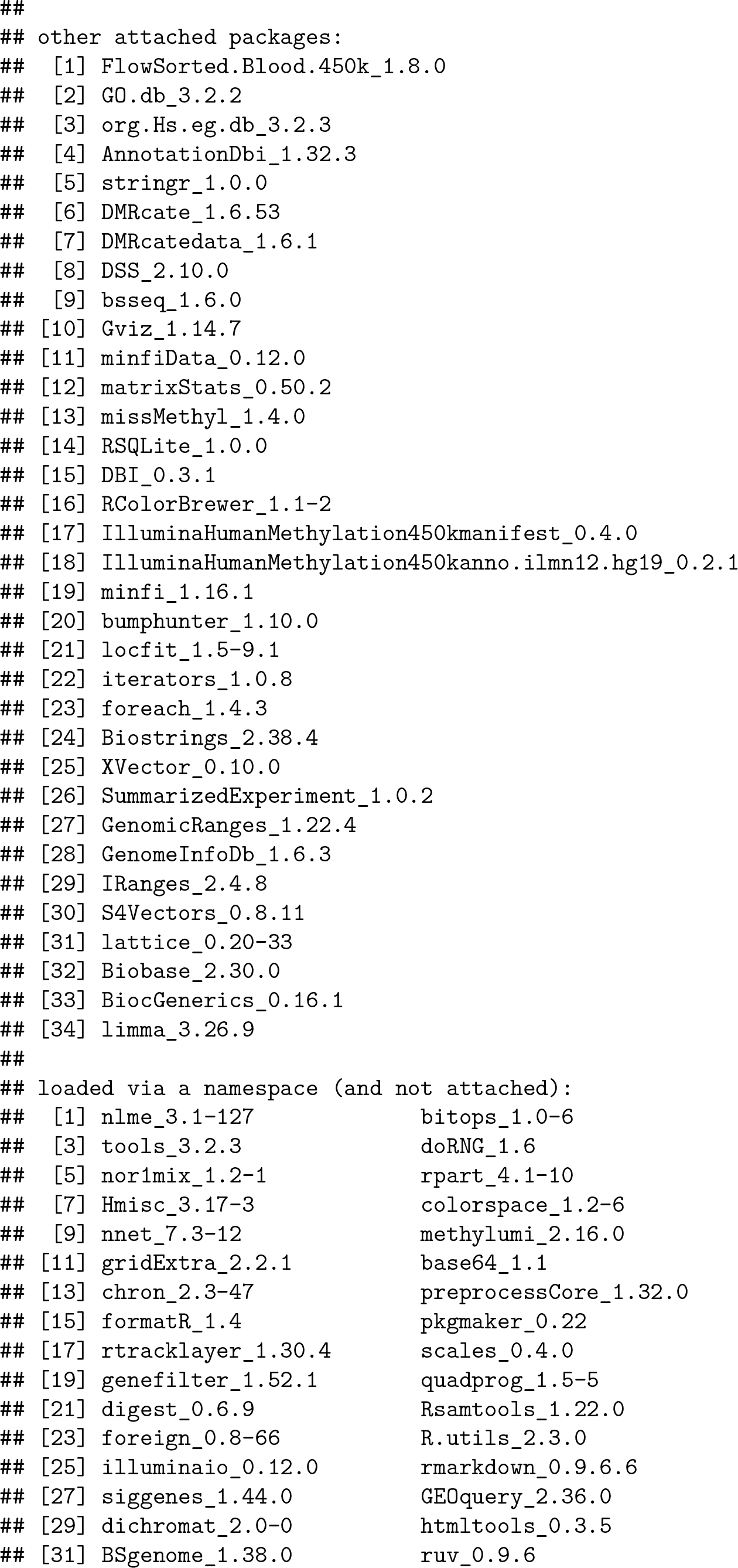

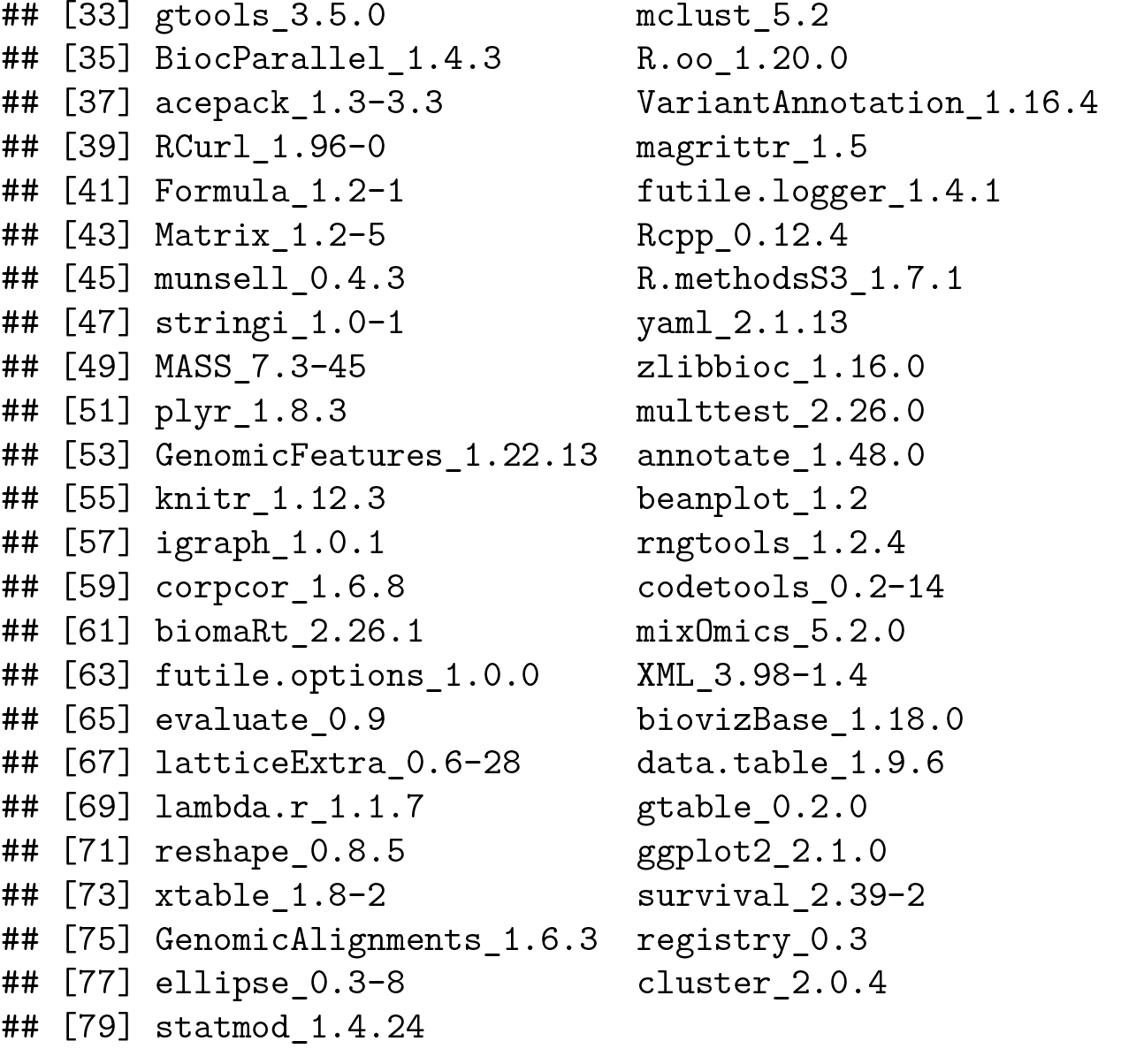

## Author contributions

JM and BP designed the content and wrote the paper. AO oversaw the project and contributed to the writing and editing of the paper.

